# Mapping the free energy landscape of K-Ras4B dimerization

**DOI:** 10.1101/2025.04.22.650040

**Authors:** Panagiotis I. Koukos, Nastazia Lesgidou, Sepehr Dehghani-Ghahnaviyeh, Camilo Velez-Vega, José S. Duca, Zoe Cournia

## Abstract

KRAS-4B regulates cellular proliferation and differentiation via its GTPase activity, and it is often mutated in human tumors. Deregulation of the MAPK/ERK pathway as a result of K-Ras4B mutations leads to uncontrolled proliferation, with the dimer/multimerization of K-Ras4B on the plasma membrane believed to be the initiating event for subsequent MAPK/ERK signaling. While K-Ras4B proteins are known to cluster on the plasma membrane, whether they associate through well-defined dimerization interfaces remains an open question. Here, we present the dimerization landscape of active, GTP-bound wild-type and G12D mutant K-Ras4B using coarse-grained unbiased and enhanced sampling molecular dynamics simulations. We recover the experimentally-reported K-Ras4B interfaces, and additionally unveil rugged free energy landscapes with many-yet uncharacterized-minima that feature c-Raf-mediated dimerization interfaces. We further explore whether wild-type or G12D K-Ras4B present different dimerization states, revealing that the G12D mutant is more likely to form diverse dimers compared to WT K-Ras4B. Our work presents evidence that K-Ras4B proteins likely interact through multifaceted interfaces that may enable controlled dimerization in different conformations from a single system, efficiently promoting nanoclustering. Although many weak, non-specific interfaces are forming, the most dominant interfaces occur with nanomolar affinity, offering a structural basis for the design of ligands able to modulate K-Ras4B dimers.

## INTRODUCTION

Kirsten rat sarcoma (*KRAS*) is the most frequently mutated gene of the RAS family^1^, with *KRAS4B*, which encodes the K-Ras4B protein, being the dominant splice variant over *KRAS4A*^2^. All *RAS* genes encode for small GTPases, which act as binary switches alternating between active, GTP-bound and inactive, GDP-bound states^1^. Recent analyses show that the frequency of *KRAS* mutants in human cancers range between 15-30%.^3,4^ KRAS-associated mutations still cause significant disease burden since they are the most frequently encountered mutations in 92%, 49% and 35% of pancreatic ductal adenocarcinoma, colorectal cancer and non-squamous non-small cell lung cancer cases, respectively, with G12D being the most frequent mutation.^5^ While long considered an undruggable target^6^, recently developed covalent allosteric inhibitors targeting the G12C mutation have been granted FDA approval,^7,8^ with more bioactive molecules targeting the K-Ras4B pathway in active development.^9,10^

Disease phenotypes as a result of *KRAS* mutations usually manifest through the ERK/MAPK or PI3K/AKT/mTOR signaling cascades, with the ERK/MAPK pathway being the best characterized to date.^11^ Signaling through the ERK/MAPK pathway is triggered by the multi/dimerization of the GTP-bound, active form of Ras, which acts as a scaffold for the dimerization and subsequent activation of Raf kinase.^12^ The Raf domains that are most relevant for the interaction with Ras are the Ras binding (RBD) and cysteine rich domains (CRD). The RBD interacts directly with Ras, whereas the CRD interacts both with Ras and the plasma membrane anionic lipids.^12^ The initiating event for this signaling pathway is thought to be the spatial gathering of 4-7 Ras protomers on the plasma membrane, forming a Ras nanocluster, ^13–15^ where lipid-protein interactions are believed to be the driving force for the formation of the nanocluster.^16–18^

Over the years, multiple interacting surfaces have been proposed as candidate dimerization interfaces. The most commonly reported interfaces are the α4-α5 interface,^19^ the α3-α4 interface,^20^ and the interfaces involving beta sheets and switches: the β2-switch1^20^ being observed in the absence as well as the presence of inhibitory compounds and the β1-switch2,^21–24^ which is observed only in the presence of inhibitors. More recently, an alternative G12D-specific dimerization interface has been described with α-β interacting regions^25^ and another where α4-α5 helices of one K-Ras4B monomer interact with the GTP of the other monomer.^26^ Figure 1 shows an illustration of the secondary structure nomenclature of Ras proteins.

**Fig. 1.**
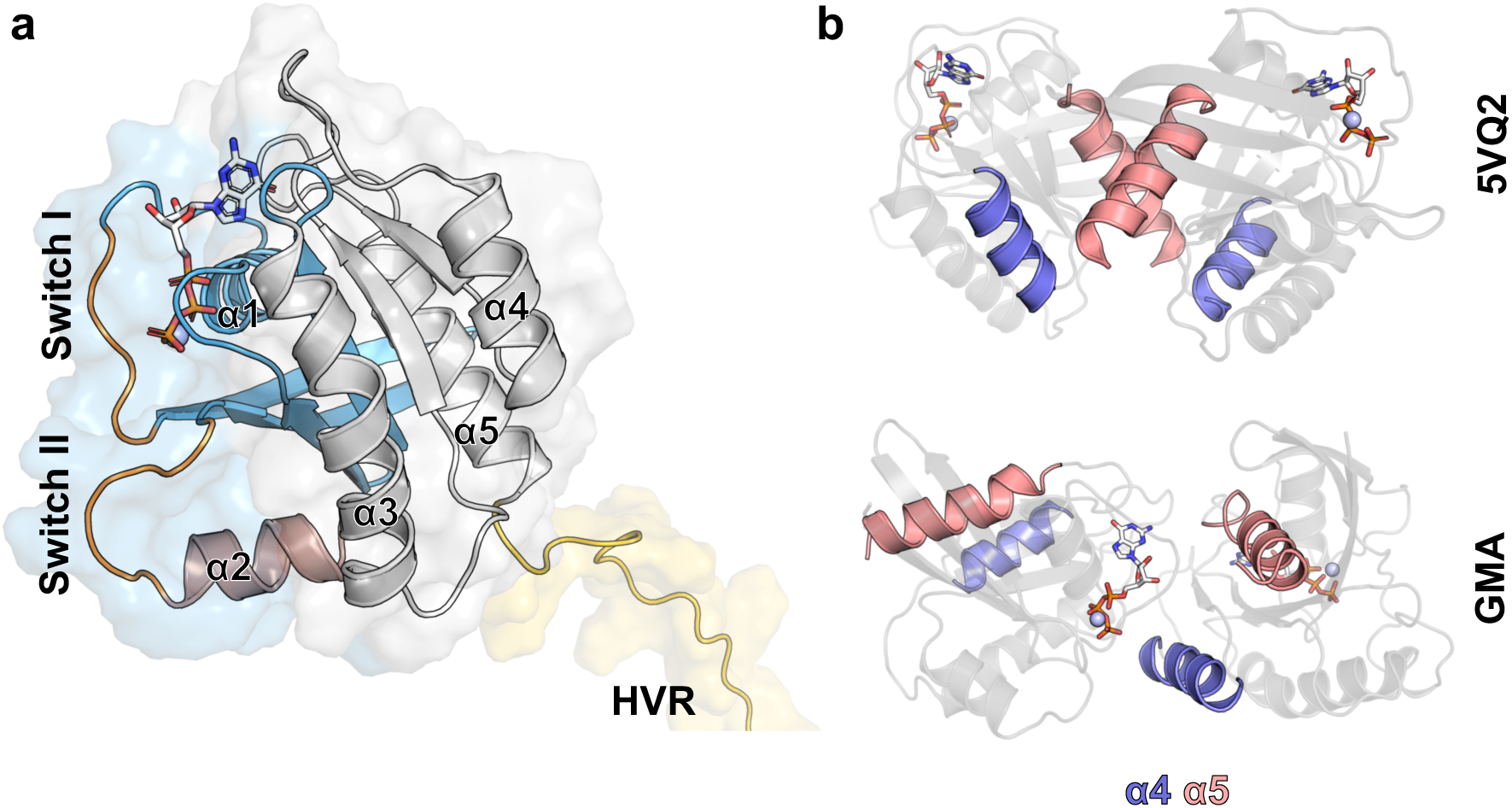
Secondary structure nomenclature of K-Ras4B. (a) Cartoon representation of K-Ras4B with helices α1, α2 and α3-α5 colored sky blue, light brown and light grey, respectively. Switch I and II are colored light orange and the HVR light yellow. GTP is represented as sticks and the Mg^2+^ ion as a light blue sphere. (b) Starting structures for the simulations presented herein: PDB ID: 5VQ2^36^ (top) and the structure from Mysore et al.^26^ (bottom) with helices α4 and α5 colored slate blue and salmon, respectively.

Of these candidate dimerization interfaces, the α4-α5 interface has received the most attention due to a study^19^ showing that mutants of oncogenic K-Ras4B variants carrying mutations which are located on the α5 helix resulted in impaired downstream signaling through the MEK/ERK pathway. However, follow-up studies have failed to replicate these findings^27,28^. On the opposite end, a study by Chung and colleagues,^29^ showed that K-Ras4B remains monomeric across a range of density values, artificial membrane compositions and type of nucleotide bound. More recent findings have identified WT and mutant K-Ras4B nanoclusters in native membranes, using diverse techniques, such as transmission electron microscopy^16,18,30^ and fluorescence-assisted microscopy^31^. However, neither of these techniques is capable of yielding atomic-, residue- or even domain-level insights regarding the interacting surface areas of the K-Ras4B proteins. K-Ras4B dimerization and nanocluster formation has also been approached computationally with all-atom, lipid-bilayer molecular dynamics simulations of K-Ras4B in the presence and absence of its Raf [RBD-CRD] effectors,^32^ and multiscale modelling of K-Ras4B in a native-like membrane bilayer.^33^ Both studies emphasized the significance of protein-lipid interactions for the formation of dimers and oligomers, with the findings of the latter being corroborated by follow-up single cell particle tracking experiments.^34,35^

Here, we employ coarse-grained enhanced sampling molecular dynamics (MD)-based protocols for the study and characterization of the dimerization free energy landscape of the K-Ras4B bound to its c-Raf [RBD-CRD] effector using Martini 3^37,38^ and parallel tempering metadynamics in the well-tempered ensemble (PT-MetaD-WTE) simulations.

We first highlight the capability of our approach to identify previously-reported experimentally or computationally determined K-Ras4B homodimerization interfaces. We then demonstrate that, contrary to popular belief, the clustering of two K-Ras4B proteins does not arise solely due to the association of two conserved K-Ras4B G-domain interfaces but, instead, is a consequence of diverse dimerization interfaces, between K-Ras4B dimers as well as K-Ras4B and c-Raf [RBD-CRD] subunits. We calculate the 2D free energy homodimerization landscape of the WT and G12D mutant K-Ras4B systems and extract representative structures from local minima of the free energy landscape of each simulation. The comparison of the identified minima structures with published K-Ras4B complex structures shows that many of the previously observed interaction modes are part of the free energy landscapes of our simulations, thus supporting our methodology. Additionally, many novel interaction modes are revealed. The comparison of minima from different parts of the landscape reveals a complex and rugged dimerization free energy landscape, with highly diverse dimerization modes. While most identified interfaces are medium-to-low affinity, high-affinity dimers are also present, including c-Raf-mediated interactions that are found to be more frequent compared to the ones featuring exclusively K-Ras4B contacts (K-Ras4B – K-Ras4B). We conclude by analyzing protein-lipid interactions, the diffusion rate of protein complexes, and the membrane curvature induced onto the bilayer by the presence of the K-Ras4B-c-Raf protein complex. Overall, although many weak, non-specific dimer interfaces are forming, the most dominant interfaces occur with nanomolar affinity, especially for the dimers without effectors, which may allow structure-based design of ligands capable of modulating the formation and activity of K-Ras4B dimers on the membrane with therapeutic potential against cancer.

## RESULTS

### Unbiased CG simulations of two K-Ras4B monomers on a lipid bilayer reveal novel interaction modes

We initially performed ten independent unbiased coarse-grained simulations of two K-Ras4B monomers each bound to c-Raf RBD and CRD domains (henceforth referred to as c-Raf [RBD-CRD] with effectors) in the presence of a DOPC:DOPS:PIP_2_ membrane at 75:20:5% mol. concentration, for 10 μs per replica (see Methods). For all ten replicas, we then computed the interface root mean square deviation (RMSD) relative to the α4-α5 interface of PDB ID: 5VQ2 and the α4-α5-GTP interface recently proposed by Mysore et al.^26^ (GMA, Figure 1b). The interface RMSD values of the plots clearly indicate that these two interfaces are not visited in any of the ten replicas (Figure S1). Clustering the trajectory of each replica independently (see Methods) yielded a total of 17 representative structures. Figure S2 shows the time evolution of RMSD values compared to the representative structure(s) for each replica. The similarity of all representative structures was quantified by calculating the RMSD between all pairs of representative structures, grouping the structural pairs in groups of high (RMSD < 5 Å), acceptable (5 Å ≤ RMSD < 10 Å), and low similarities (10 Å ≤ RMSD < 15 Å), shown in Figure S3 matrix as red-, blue- and light grey-colored cells, respectively. Almost 90% (122 out of 136) of the interfaces are completely dissimilar compared to representative structures (RMSD ≥ 15 Å, see Figure S3, white-colored cells). Of the remaining 10% (14 out of 136) of structural pairs, only four exhibit acceptable (5 Å ≤ RMSD < 10 Å) or high similarity (RMSD < 5 Å), with the remaining 10 structures classified as low-similarity (10 Å ≤ RMSD < 15 Å). Figure 2 shows representative structures along the membrane z-axis and the x-y plane. Perhaps the most surprising feature that these structures share is the absence of direct inter-protomer K-Ras4B interactions, and specifically the lack of a K-Ras4B – K-Ras4B protein-protein interface.

**Fig. 2.**
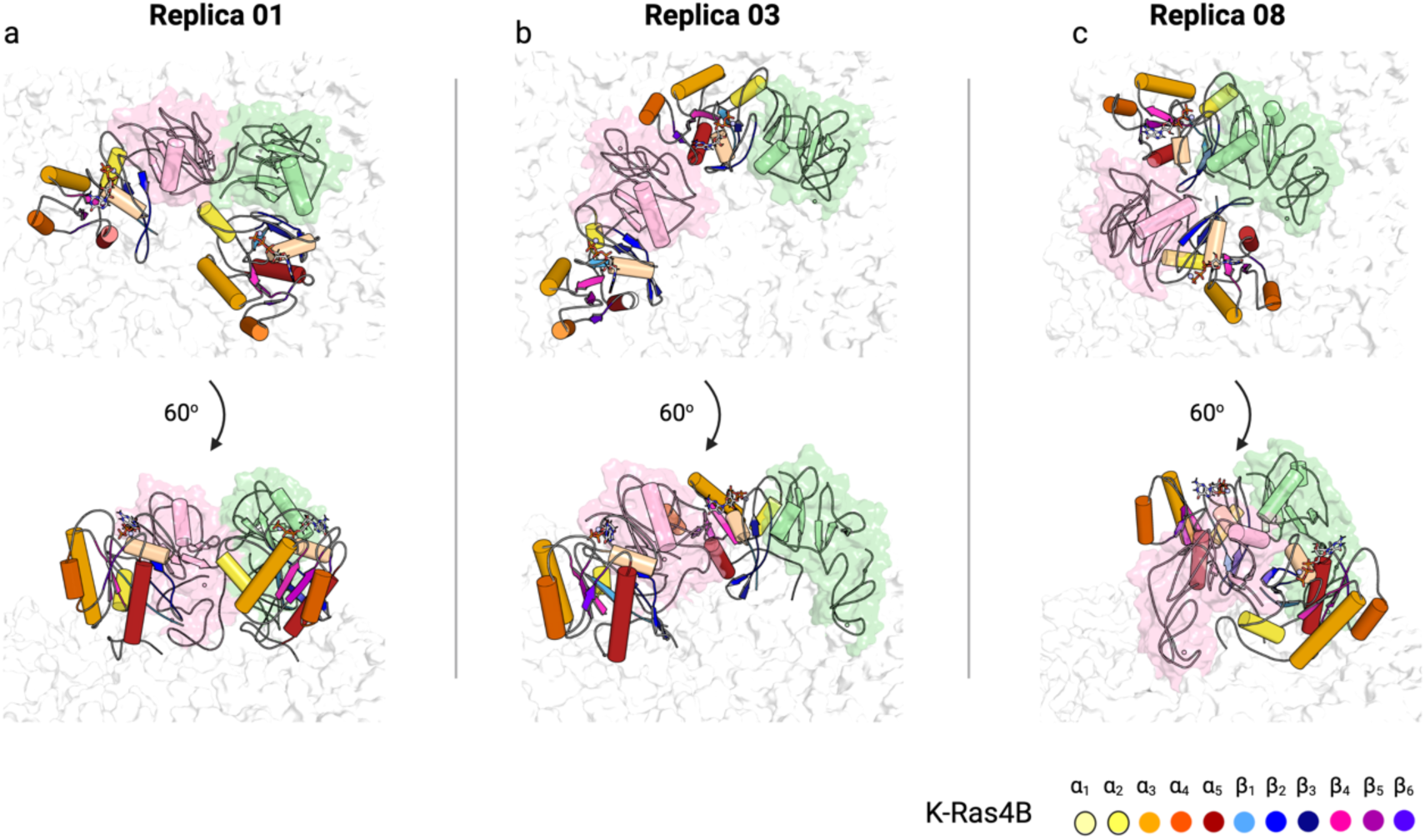
Representative structures from the unbiased simulations. Structure of the major cluster of replicas 1 (a, 84% population), 3 (b, 63% population) and 8 (c, 73% population) viewed along the membrane x-y plane (upper row) and the membrane-axis (lower row). K-Ras4B coloring is the same as for Figure 1a. Raf are shown as light pink and green cartoons, with the Zn^2+^ ions shown as dark gray spheres. K-Ras4B structural elements are colored according to the legend shown in the image. The membrane bilayer is represented as white spheres. The major cluster structures of replicas 1, 3, 8 were selected for visualization because they share the greatest degree of similarity to the remaining representative structures (Figure S3).

Even more surprisingly, we discovered that the majority of the inter-protomer interactions across all representative structures formed between the c-Raf [RBD-CRD] subunits or between the inter-protomer K-Ras4B and c-Raf [RBD-CRD] subunits, with only two of the 17 representative structures interacting through K-Ras4B interfaces. This finding is surprising because it contrasts the established notion that K-Ras4B forms homodimers that act as a scaffold for c-Raf kinase to dimerize itself.^32^

### 2D dimerization free energy landscape of WT and G12D K-Ras4B using CG-Metadynamics simulations

The findings of the unbiased simulations led us to investigate the free energy landscape of K-Ras4B in the presence of c-Raf [RBD-CRD] using CG simulations with Parallel Tempering Metadynamics in the Well-Tempered Ensemble (PT-MetaD-WTE)^39,40^ following a protocol previously developed in our group to study protein dimerization^38^. We considered six systems with initial configurations and components shown in Table 1.

**Table 1.** Simulated systems. The first and second columns indicate the name and details of each system simulated and the third the duration of the simulation in μs.

| Name | System | Simulation length (μs) |
| --- | --- | --- |
| WT without effectors | Wild type K-Ras4B without effectors starting from the α4-α5 interface of PDB ID: 5VQ2. | 15 |
| WT-GMA without effectors | WT K-Ras4B without effectors starting from the α4-α5-GTP interface recently proposed by Mysore et al.<sup>26</sup> | 15 |
| G12D without effectors | G12D K-Ras4B mutant without effectors starting from the α4-α5 interface (PDB ID: 5VQ2). | 15 |
| WT with effectors | WT K-Ras4B with c-Raf [RBD-CRD] effectors starting from the α4-α5 interface (PDB ID: 5VQ2). | 15 |
| WT-GMA with effectors | WT K-Ras4B with c-Raf [RBD-CRD] effectors starting from the GMA interface. | 15 |
| G12D with effectors | G12D K-Ras4B with c-Raf [RBD-CRD] effectors starting from the α4-α5 interface (5VQ2). | 15 |

We have used two Collective Variables (CVs): a) the RMSD from a given reference structure and b) intermonomer distance (see Methods). Simulation convergence was monitored through the diffusion of CV values and the evolution of the 1D free energy profiles with respect to the RMSD CV over time (see SI for details and Figures S4-S6).

### Labeling of the 2D free energy landscapes of K-Ras4B dimerization with and without effectors

Figures 3a and 4a show the 2D free energy landscapes for all six systems (Table 1, see also Figure S6 for simulation convergence and Figures S7 and S8 for fully labeled and unlabeled versions of the free energy landscapes, respectively). Minima are marked with lower and uppercase letters (Figures 3a, 4a and S7). Labels of the minima correspond to their relative ranking with the global minimum labeled “A”, the subsequent 25 minima in order of increasing energy labeled “B” to “Z”, and the remainder labeled with lowercase letters (up to “y” for the WT-GMA simulation with effectors, which features the most minima). Minima with the same label across different simulation do not necessarily correspond to the same structure. In our previous work,^38^ we had shown that Martini 3 force field is able to predict dimer crystal structures, but cannot always rank the identified minima correctly. Therefore, here we only considered minima whose energy is <-6 kcal/mol (see “Identification of minima from the 2D free energy landscapes” section in “Methods”).

**Fig. 3.**
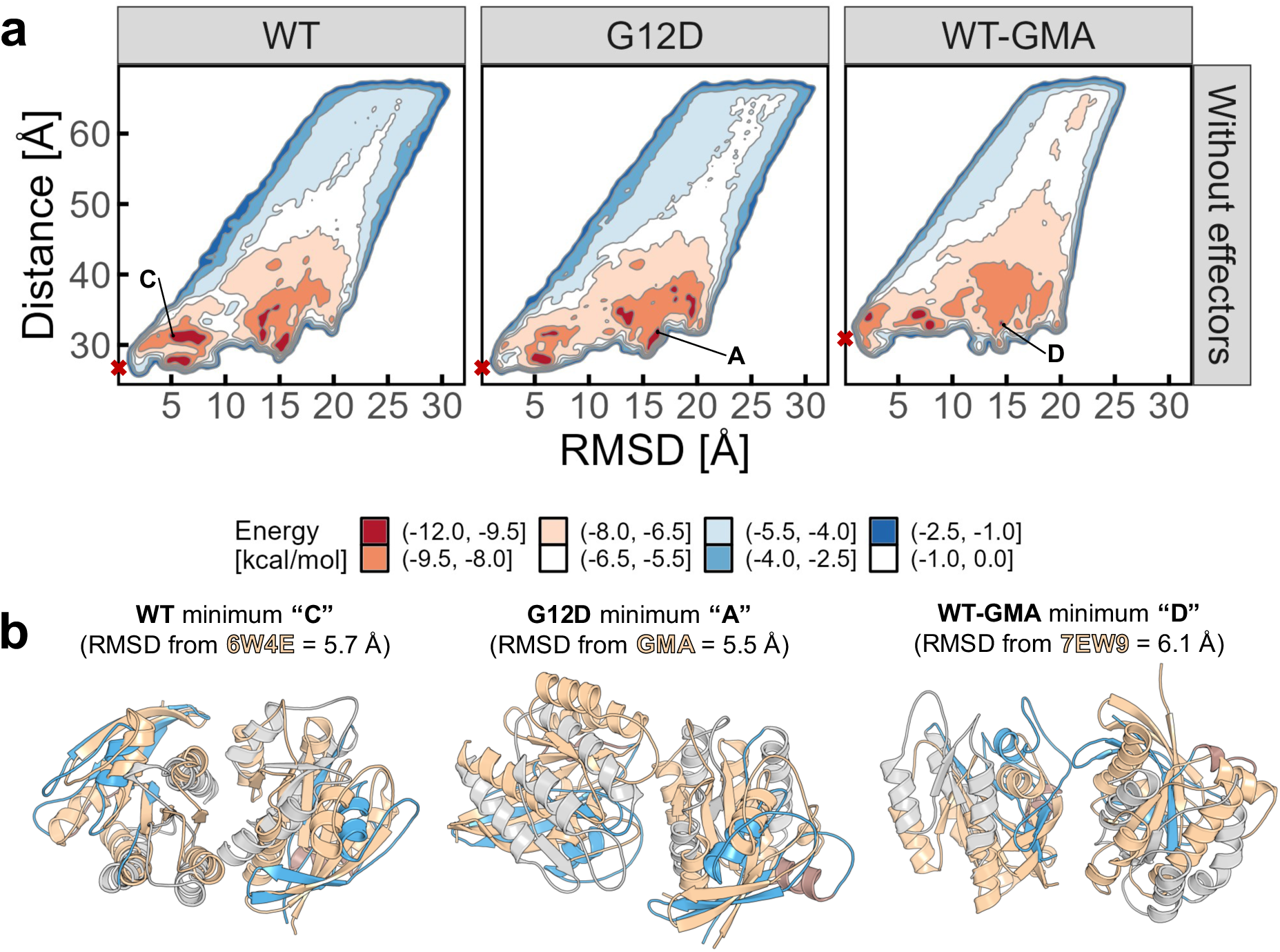
(a) 2D free energy landscapes of K-Ras4B without c-Raf [RBD-CRD].) Red crosses indicate the starting coordinates (in CV space) for the three simulations, WT and G12D starting from PDB ID: 5VQ2^36^ and WT-GMA starting from the structure of Mysore et al.^26^ Local minima are labeled according to their energetic ranking from A (lower energy, global minimum per simulation) through Z and a through y (higher energy). Energy values have been normalized according to the lowest energy observed across all systems (global minimum of WT-GMA without effectors). The same label does not necessarily correspond to the same minimum across simulations, as it only reflects the energetic ranking, which is carried out on a per-simulation basis. (b) Comparisons of minima with reference structures. Minima structure coloring is the same as in Figure 1a and reference structures are shown as wheat-colored cartoons. GTP and Mg^2+^ are not shown for clarity. Loops are shown simplified for clarity.

**Fig. 4.**
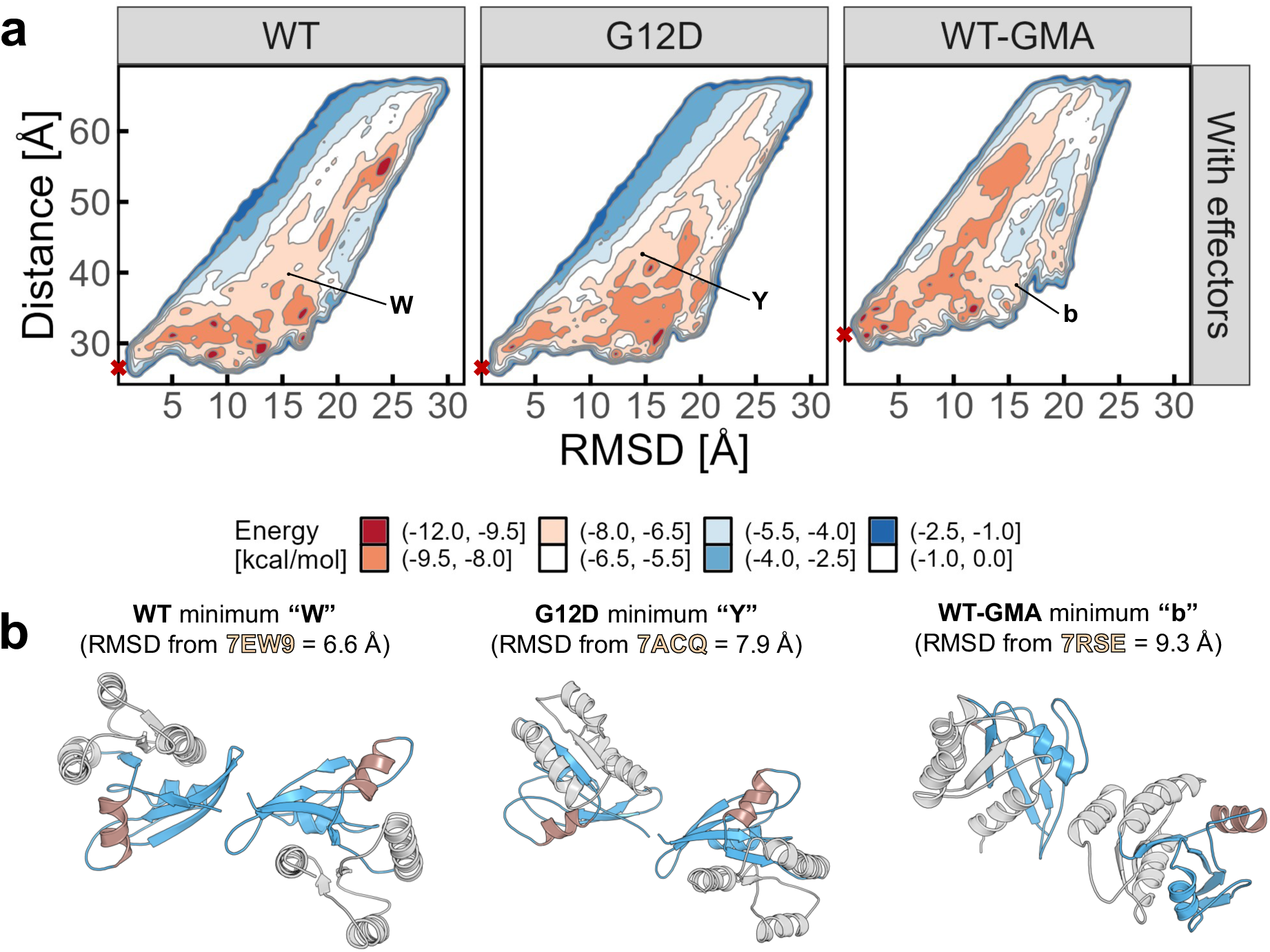
(a) 2D free energy landscapes of K-Ras4B with c-Raf [RBD-CRD]. Red crosses indicate the starting coordinates (in CV space) for the three simulations, WT and G12D starting from PDB ID: 5VQ2^36^, WT-GMA starting from the structure of Ref.^26^ Local minima are labeled according to their energetic ranking from A (lower energy, global minimum per simulation) through Z and a through y (higher energy). Energy values have been normalized according to the lowest energy observed across all systems (global minimum of WT-GMA without effectors). The same label does not necessarily correspond to the same minimum across simulations, as it only reflects the energetic ranking which is carried out on a per-simulation basis. (b) Structures resembling known interfaces. Reference structures are not shown for clarity - see Figure S12 for a version of this figure, which also includes the superimposed reference structures.

### Simulations without c-Raf [RBD-CRD] effectors identify existing interfaces and reveal interplay between interfaces

Minima “A” and “C” for the WT simulation without effectors are closest to the native state of the simulation (PDB ID: 5VQ2) with RMSD values equal to 6.8 and 5.8 Å, respectively; minimum “A” corresponds to PDB ID 5VQ2^36^ and minimum “C” to PDB ID 6W4E^41^ (see Figure 3b). For the G12D simulations, minimum “B” also corresponds to structure with PDB ID 5VQ2 and “G” to 6W4E, both with RMSD values equal to 6.5 Å, indicating that these interfaces occupy adjacent and closely related parts of the landscape.

Interestingly, the two deepest minima “A” and “B” of the G12D and WT simulations, respectively, have RMSD values from the GMA reference structure^26^ equal to 5.5 and 6.2 Å – a structure which neither simulation is being biased toward (see Figure 3b), and is actually far from the native state (PDB ID 5VQ2, RMSD > 10 Å). The fact that this structure is spontaneously detected as a nanomolar interface in the free energy landscape points to the conclusion that this is indeed a viable interface, which remains to be verified experimentally. However, it is noted that the “GMA” interface structure is not detected in the presence of effectors.

The 2D free energy landscape of the WT-GMA simulation without effectors reveals that when starting from this configuration, the system is relatively more constrained compared to the other two simulations. This constrained behavior is reflected in the absence of deep minima and an overall shallower landscape away from the native state (RMSD > 10 Å), in contrast to the WT and G12D simulations (See Figure 3a – WT-GMA, for more information see the SI). However, minima of the WT-GMA simulation resemble other experimentally resolved structures. For example, the dimer observed in the G12D PDB entry 7EW9^42^ (dimer formed by PDB entry chains B and C) has an RMSD of 6.1 Å with minimum “D” (see Figure 3b).

Similarly, in G12D simulation without effectors, minimum “D” resembles PDB entry 7ACA^24^. Interestingly, both crystal structures with PDB IDs 7EW9 and 7ACA were solved in the presence of small molecule inhibitors, which target the active and inactive states of G12D K-Ras4B.^24,42^

Figure S9 shows the comparison of all identified minima against crystallographic structures of interest. Structural comparisons among representative minima structures that share similar structural features across simulations are summarized in Figure S10, whereas in Figure S11 the comparisons are further broken down by similarity into high (RMSD < 5 Å), acceptable (5 Å ≤ RMSD < 10 Å), low (10 Å ≤ RMSD < 15 Å) and no similarity (RMSD ≥ 15 Å). This analysis reveals the multiple points where the WT and G12D simulations without effectors converge with a high degree of similarity: most convergence points resemble the α4-α5 interfaces (PDB entries 5VQ2 and 6W4E, with 5VQ2 being the structure both simulations are being biased toward) or the GMA interface. The single point of convergence that moderately resembles another reference structure is minima “I” and “O” from the WT and G12D simulations, respectively, whose RMSD values from PDB entry 7ACA are 7.8 and 8.9 Å, respectively. Another significant observation is the fact that even though none of the reference α4-α5 interfaces are anywhere close to the minima of the WT-GMA simulation (see Figure S9), parts of the conformational landscape of the 5VQ2-based (WT and G12D) and GMA-based (WT-GMA) simulations overlap (see Figures S10 and S11). This is best illustrated by the high similarity between minima “R” and “a” of the WT and G12D without effector simulations, respectively, with minimum “I” of the WT-GMA simulation (Figure S11 matrices and inset), as well as the 19 out of 31 distinct WT-GMA minima that share at least acceptable (RMSD < 10 Å) similarity with one of the WT or G12D minima.

### Simulations with c-Raf [RBD-CRD] effectors show significance of c-Raf-mediated interactions

The rugged 1D free energy profiles with respect to the RMSD CV (see Figure S6) for the simulations with effectors is also reflected in the 2D free energy landscapes as well (Figure 4a). Comparing the landscapes of the simulations with and without effectors (Figure 3a vs 4a) shows that simulations without effectors explore deeper minima around the state towards which each simulation is being biased, compared to the simulations with effectors, which explore shallower minima in the same part of the landscape. The landscapes of the simulations with effectors have more distinct minima compared to no effectors (141 vs 102), indicating the presence of more diverse interactions (see Discussion for details).

In terms of similarities with known interfaces, overall, the percentage of minima that have at least one match among the experimentally-determined K-Ras4B structures has increased for the WT simulation with effectors when compared with the simulation without effectors (29% vs 43%) and down for the G12D simulation (46% vs 30%). No significant differences were observed for the WT-GMA simulations (23% vs 20% for the simulation with and without effectors, respectively). Interestingly, none of the minima for the WT and G12D simulations with effectors resemble the GMA interface, unlike their counterparts without effectors. For more discussion of the similarities with known interfaces and the WT-GMA simulation free energy landscape, see the SI.

Unlike the simulations without effectors, the WT and G12D simulations with effectors explore a completely different part of the conformational landscape compared to the WT-GMA simulation. This is illustrated by the percentage of WT-GMA minima structures that have no similarity to the minima of the WT or G12D simulations, which is approaching 100% (see Figure S10). The minima of the simulations with effectors resemble less each other than their counterparts without effectors, with percentages of minima of each simulation that are dissimilar (RMSD > 15 Å) to all other minima of the same simulation equal to 81.8, 88.7 and 81.6% for WT, G12D and WT-GMA, respectively, further reinforcing the more diverse nature of the minima of the simulations with effectors. The minima of the G12D simulation with effectors stand out as the ones bearing the smallest similarity to each other.

The landscapes of the simulations with effectors are also more complex relative to those without effectors. For example, we observe the presence of diverse structures even at distances greater than 40 or 50 Å between the two K-Ras4B monomers, in particular for the WT simulations. This large distance does not mean that the two protomers are dissociating, but rather that the inter-protomer interaction is being mediated by c-Raf instead of K-Ras4B monomers. Figure 5 highlights six minima with K-Ras4B distances > 40 Å along with a 3D visualization of their free energy landscapes (see Figure 4a for 2D versions of the same landscapes). The presented minima structures are the two deepest minima from each simulation for which there is no contact between the K-Ras4B subunits, excluding any similar interfaces.

**Fig. 5.**
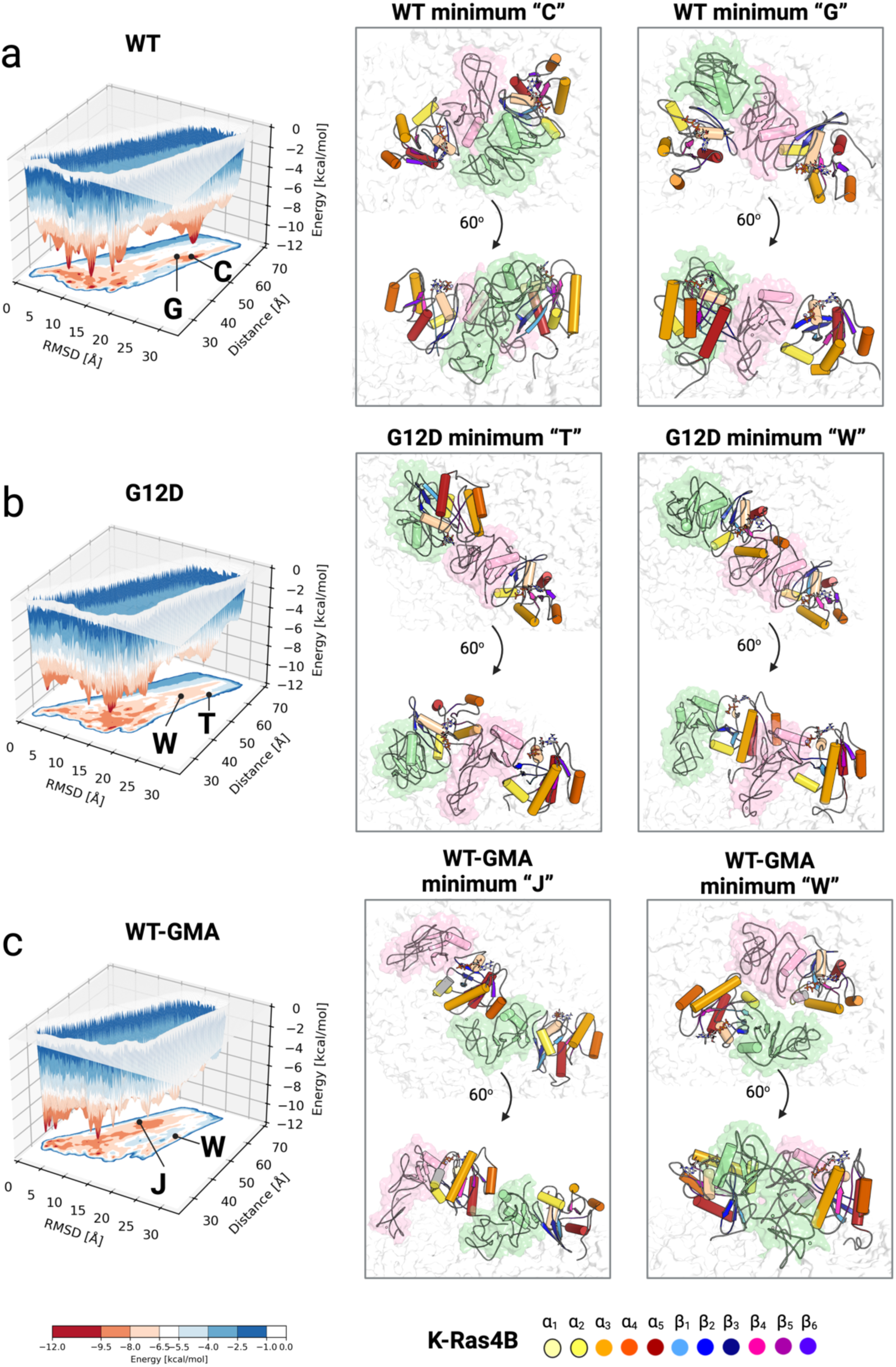
3D landscapes for the simulations with effectors and selection of minima without direct inter-monomer K-Ras4B – K-Ras4B interactions. C-Raf [RBD-CRD] are colored in green and pink and K-Ras4B structural elements are colored according to the labels shown in the image. The minima labels reflect the per-simulation ranking of minima and the same label does not necessarily correspond to the same structure.

Figures 5a-c show interactions where one or both c-Raf monomers interact with the partner K-Ras4B and c-Raf monomers or interactions where the inter-protomer interface is only shared between one K-Ras4B subunit and the c-Raf subunit of its partner.

We then calculated the contact profile for all identified minima by analyzing the inter-protomer contacts and classified them in three categories, depending on which subunits were making contact: a) RAS-RAS, when the contact occurs between K-Ras4B residues of both partners, b) RAS-RAF, when a K-Ras4B residue of one partner contacts a c-Raf residue of the other partner, c) RAF-RAF, when c-Raf residues of both partners are interacting(see Figure S13 for a per-minima breakdown of the contact-making residues). The results of the analysis are shown in Figure 6a, where it becomes obvious that the RAS-RAS and RAS-RAF contacts are almost equally represented in the WT and G12D simulations with percentages ranging between 45 and 50%. Also, the WT-GMA simulation is significantly skewed in favor of the RAS-RAF and RAF-RAF interactions. For a more detailed analysis see the caption of Figure S13.

**Fig. 6.**
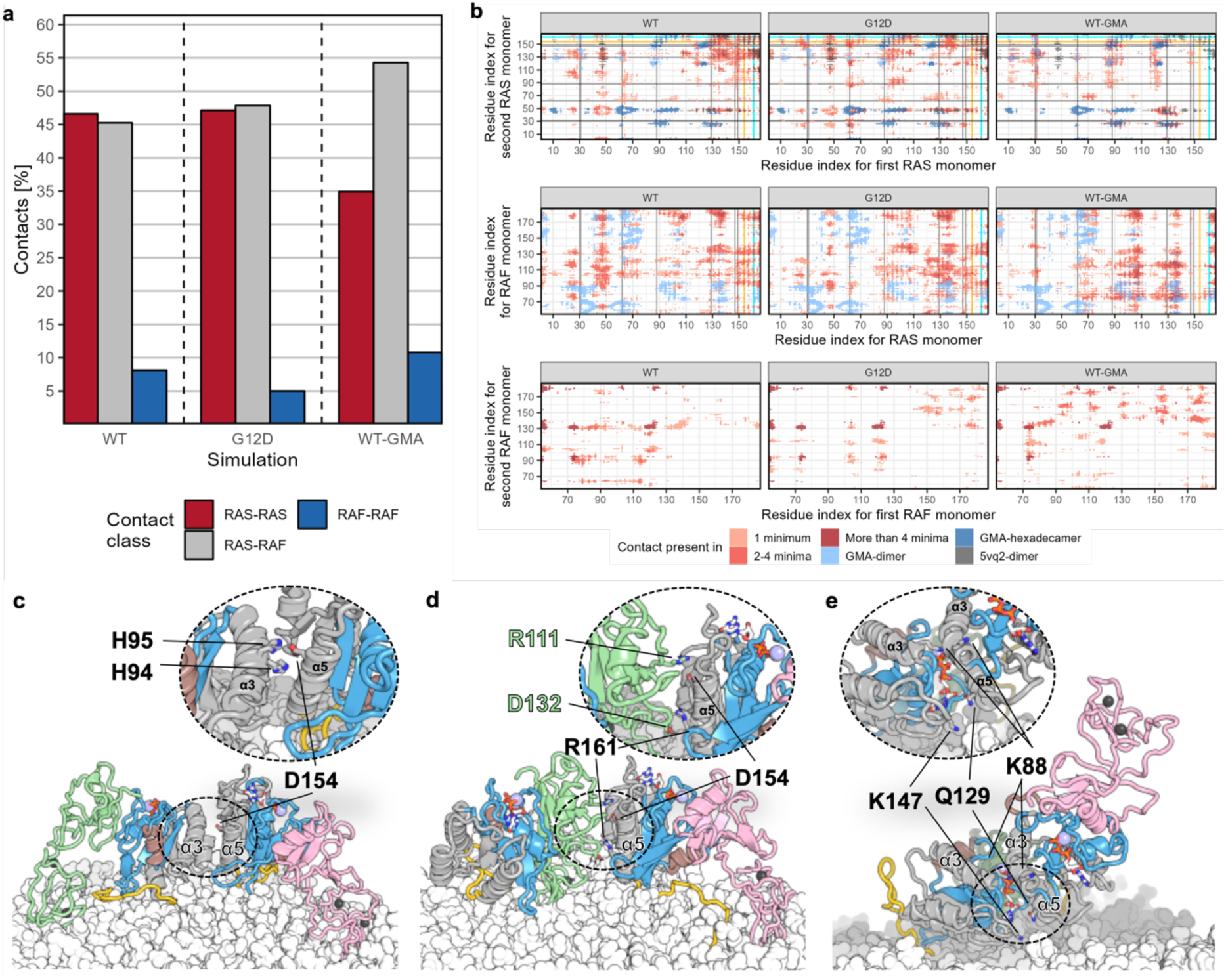
Contact analysis for the simulations with effectors. (a) Percentage of contact class for simulations with effectors, shown as colored bars. (b) Contact map for all minima structures, simulation reference structures and the proposed hexadecameric “signalosome” from Mysore et al.^26^ The cyan (residue 161), black (residues 30, 31, 62, 88, 129, 147 and 149) and orange (residue 154) lines indicate the positions of K-Ras4B residues, which have been used in mutagenesis experiments. Red indicates the number of minima structures in which each contact is present. Light, medium and dark red indicate the presence of each contact in one, two to four and more than four minima structures, respectively. Contacting residues for the GMA dimer, hexadecamer and PDB ID 5VQ2 are colored light blue, blue and grey, respectively. (c-e) Minima structures in which residues of interest from panel a can be found in interfaces other than those they were tested for. Coloring of structures for panels c-e is the same as for Figure 1 for K-Ras4B; c-Raf [RBD-CRD] are colored in green and pink.

This finding is consistent with the observation from the unbiased simulations that the majority of inter-protomer interfaces were not mediated through the direct interaction of K-Ras4B subunits, but through RAS-RAF interfaces instead. The fact that the majority of inter-protomer interfaces involve RAS-RAF or RAF-RAF interactions for all simulations with effectors, could indicate a significance of the interface interaction diversity for the multimerization landscape of K-Ras4B, as these interactions may collectively contribute significantly to the total free energy of the system in a highly configuration-dependent manner, thereby substantially modulating the kinetics and/or equilibria of a variety of biochemically important reactions (see Discussion).^35,43^

### Simulations show that multiple novel interfaces comply with existing experimental data

Using the contact data mentioned previously, we calculated the residue-based contacts of all accessible residues of all minima for all simulations with effectors (see “Calculation of inter-protomer contacts and contact class analysis” section in Methods) and identified the number of minima in which each contact was present. (Figure 6b).

The selected contacts were chosen from experimentally resolved structures. In Figure 6b the lines corresponding to D154 and R161 (shown as orange and cyan lines, respectively) are on the well-characterized α4-α5 dimerization interface (see also Figure S14 for a depiction).^19^ The black and orange lines of Figure 6b correspond to residues that were mutated experimentally for the characterization of the GMA model, which were shown to either impede the primary dimerization interface (α4-α5 – GTP) or the secondary/tertiary interfaces (residues 88, 129 and 149)^26^: residues 30, 31, 62, 88, 129, 147, 149 (black lines) and 154 (orange line).

Interactions involving the charge-reversal mutations of the α4-α5 helices (K-Ras4B residues 154 and 161, shown as orange and cyan-colored lines, respectively) and regions of K-Ras4B other than α4-α5, highlight RAS-RAS interactions through alternative interfaces. The existence of these interfaces suggests that the observed deleterious impact of these mutations (with respect to nanocluster assembly and downstream signaling) could be explained through interactions of these residues between interfaces other than α4-α5 (for example, see Figure 6c-e). The same holds true for the mutations examined in the context of the GMA interface affecting the same part of K-Ras4B (residues 147 and 149, black lines). This finding signifies that residues, which have been previously proposed to play a key role in the dimerization of specific interfaces, may in fact participate in multiple dimerization interfaces. Importantly, most of the K-Ras4B residues analyzed here are also shown to liberally interact with c-Raf as well, as can be seen by the intersection of the various black lines (black, cyan or orange) with high-density areas (medium-dark red) of the contact maps for the RAS-RAF interactions (see Figure 6a, middle row and Figures 6c-e). For a more detailed structural explanation of Figure 6, see the SI.

However, since the GMA model dictates the placement of K-Ras4B monomers in a helical arrangement with c-Raf binding their respective Ras partner laterally, we also identified the interfaces of the GMA hexadecamer (eight K-Ras4B and eight c-Raf monomers), to determine whether the interactions we observe here were also present in the GMA model as well, or are entirely novel. These are shown as dark blue regions in the contact map plot (Figure 6a). Even accounting for the possibility that some of the contacts might be shifted between the simulated systems and the GMA hexadecamer (see for example the region around residues 50-70 in the RAS-RAS interactions of the WT simulation), the overall overlap is low, in particular for the RAS-RAF and RAF-RAF interactions. The low overlap indicates that the majority of interactions observed in the extracted minima are novel.

Importantly, none of these minima structures (Figure 6c-e) resemble any of the experimental structures that we have included in our analysis (see Figure S9). This result signifies that there are multiple structures that comply with existing experimentally important residues that are shown to interact. Therefore, multiple “solutions” exist that satisfy experimental data, albeit not with a unique structure.

We also compared the novel complexes identified through the clustering analysis of the unbiased simulations with the minima from the biased simulations and the results can be found in the SI. Figure S15 shows the similarity (in terms of RMSD) between the representative structures of the unbiased simulations and the minima of the biased simulations while S16 highlights three instances where the biased and unbiased simulations converge. Figure S17 shows the similarity (in terms of RMSD) on a per-minimum/representative structure basis.

### The dimerization landscape of K-Ras4B includes few high-affinity and plethora of intermediate-low affinity binders

We then calculated the binding affinities for all minima dimer structures from the 2D free energy landscapes (Figure 7) as a function of the cluster population over the total population of clustered structures. The analysis shows that simulations without effectors reach fewer and deeper (higher-affinity) minima compared to the simulations with effectors that result in few high-affinity (K_d_ < 500 nM) minima (left of dashed blue line, Figure 7). These minima structures primarily resemble interfaces that have been previously observed, such as the α4-α5 interface of PDB entry 5VQ2 or the α3-α4 interface exemplified by PDB entry 3KKN (see Figure S9 for a complete comparison with known interfaces). However, they also include previously-uncharacterized interfaces, such as minima “C” and “D” of the WT simulations, with and without effectors, respectively. The overwhelming majority of minima for all simulations though occupy a lower-affinity part of the landscape (right of dashed blue line, Figure 7), indicating a set of interchangeable dimerization interfaces. The lowest nanomolar affinity interfaces are depicted in Figure S18 with some examples of known interfaces shown and others omitted for diversity. Known interfaces calculated to be of <500 nM affinity are shown in Figure S19. All backmapped structures can be also found in the Zenodo repository (see data availability section).

**Fig. 7.**
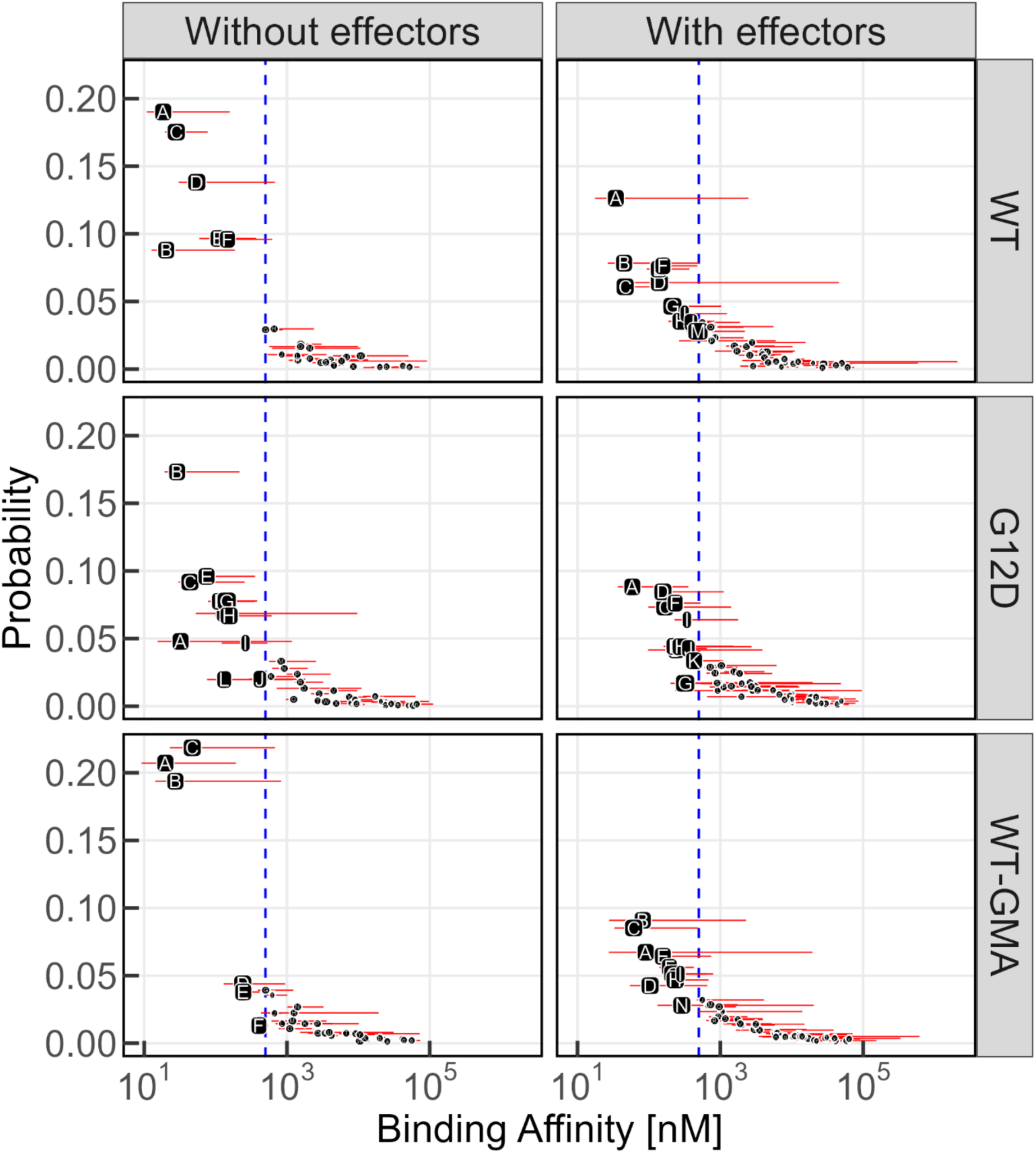
Binding affinity values for all members of all clusters identified through the minima obtained from the PT-MetaD-WTE simulations. The horizontal red lines indicate the extent of binding affinity values observed per cluster and the black points indicate the median binding affinity value of a given cluster. The dashed blue line corresponds to a binding affinity of 500 nM, and is used to identify high-affinity minima (left of the line). Probability (Y axis) indicates the population of each cluster as a ratio of the total number of trajectory frames that were grouped in clusters. Labels of clusters whose median affinity is ≤ 500 nM (left of blue line) are shown with increased font size. The X axis for all panels is in a log10 scale.

### Protein-lipid interactions

In addition to assessing the dynamics and energetics of the protein-protein interactions, we have also characterized protein-lipid interactions because they are a significant driving force for the formation of signal-conducing nanoclusters.^16,18,28,33^ We examined three metrics of protein-lipid interactions:

1. The population of lipid species close to K-Ras4B HVR and c-Raf residues through radial distribution function (RDF) analysis.
2. The diffusion of the K-Ras4b – c-Raf complex across different dimerization conditions as well as lipid diffusion in the presence and absence of the protein.
3. The membrane curvature induced by the protein.

The first analysis was performed on the biased simulations, while the second and third on the unbiased simulations.

### K-Ras4B HVR interacts more with PIP_2_ compared to DOPC and DOPS lipids

We analyzed the proximity of PIP2, DOPC, DOPS to threonine 183 of K-Ras4B HVR using RDFs in the PT-MetaD-WTE simulations with and without effectors and lysine 148 of c-Raf CRD in the PT-MetaD-WTE simulations with effectors, as these regions are known to interact with the membrane^28,44^ (Table 2, Figures S20-S21).

**Table 2.** Number of PO4 DOPC, DOPS and PIP2 beads past the second shell at a distance of 11 Å of each originating bead (K-Ras4B T183 BB and c-Raf K148 SC2 beads for the first six and bottom three rows, respectively). Numbers in parentheses reflect the percentage of each lipid type within the respective solvation shell for the protein-facing leaflet. See Figures S20 and S21 for the RDF distributions.

| **g(r)** | **BB bead T183 K-Ras4B – PO4 bead of DOPC (%)** | **BB bead T183 K-Ras4B – PO4 bead of DOPS (%)** | **BB bead T183 K-Ras4B – PO4 bead of PIP<sub>2</sub> (%)** |
| --- | --- | --- | --- |
| WT without effectors | 1.92 (0.5) | 1.1 (1) | 1.66 (6.4) |
| G12D without effectors | 1.91 (0.5) | 1.08 (1) | 1.66 (6.4) |
| WT-GMA without effectors | 2.02 (0.5) | 1.02 (1) | 1.3 (5) |
| WT with effectors | 1.87 (0.5) | 1.1 (1) | 1.71 (6.6) |
| G12D with effectors | 1.87 (0.5) | 1.08 (1) | 1.73 (6.7) |
| WT-GMA with effectors | 2.1 (0.5) | 1 (1) | 1.32 (5.1) |
| **g(r)** | **SC2 bead K148 c-Raf – PO4 bead of DOPC (%)** | **SC2 bead K148 c-Raf – PO4 bead of DOPS (%)** | **SC2 bead K148 c-Raf – PO4 bead of PIP<sub>2</sub> (%)** |
| WT with effectors | 1.61 (0.4) | 0.88 (0.8) | 0.74 (2.8) |
| G12D with effectors | 1.5 (0.4) | 0.81 (0.8) | 0.74 (2.8) |
| WT-GMA with effectors | 1.25 (0.3) | 0.71 (0.7) | 0.35 (1.3) |

Table 2 shows that all lipids cluster around the CRD of c-Raf and the HVR of K-Ras4B, with PIP_2_ clustering preferentially around these domains based on its low percentage in the membrane (see numbers in parentheses). Figure S22 shows the lipid distributions for the global minima of all simulations, demonstrating the propensity of PIP_2_ lipids to cluster around the protein.

### No differences are observed in lateral diffusion between monomeric and unstable dimeric states for the protein protomers-dimers

We calculated mean squared displacement (MSD) values for the protein and the lipids and estimated their lateral diffusion coefficient, *D_t_,* in the unbiased simulations. To examine different states of the protein, for the monomer state we selected replicas 1, 6 and 8, and determined the *D_t_* before the stable complex was formed (see Figure S2). We used replicas 2 and 3 to estimate the *D_t_* of complexes undergoing multiple association/dissociation events. Figures S23 and S24 show the MSD plots, for the monomeric and metastable dimeric states, respectively, from which the *D_t_* was calculated, after performing a linear fit on the highlighted regions. The derived *D_t_* are 10.3 ± 3.7 μm^2^/s and 8.8 ± 0.3 μm^2^/s for the monomeric and unstable dimeric states, respectively, indicate no significant differences in the diffusive regimes of the two states.

### Negatively-charged lipids diffuse faster in the absence of the protein

Comparing the lipid *D_t_* between the unbiased simulation replicas, where a stable dimer formed (see Figure S2) and a set of simulations where the same bilayer was simulated in absence of the protein, reveals differences in the lateral diffusion of the negatively charged species. Specifically, the *D_t_* of DOPC, DOPS and PIP_2_ lipids were 72.7 ± 1.3, 66.9 ± 2.6 and 50.9 ± 3 μm^2^/s, respectively, in the unbiased simulations where the protein was present in the bilayer. In the protein-free bilayer simulations, the respective values were 71.3 ± 2.9, 71.8 ± 4 and 59.4 ± 7.7 μm^2^/s. The MSD plots from which *D_t_* was calculated are shown in Figures S25-S26, while distributions of the *D_t_* values in the form of boxplots are shown in Figure S27. While the neutral DOPC has the same *D_t_* in presence and absence of the protein, the negatively charged DOPS and PIP_2_ lipids are affected by the presence of the protein, with the more negatively charged PIP_2_ (nominal charge −4) more strongly impacted compared to DOPS (nominal charge −1). However, none of the measured diffusion constant differences are significant across simulation conditions.

### The K-Ras4B-c-Raf protein complex induces changes in membrane curvature

We calculated the mean membrane curvature of the bilayer in the presence and absence of the K-Ras4B-c-Raf protein complex after the dimer complexes have formed (4-10 μs) using the unbiased simulations (Figure S28). While the protein-free bilayer has almost zero curvature, the protein-associating bilayer undergoes significant deformation along the membrane normal, resulting in the formation of locally concentrated areas of curvature. These mainly manifest as concave areas (shaded blue), where the bilayer is depressed as a result of its interaction with the protein complex, the location of which is indicated by the cross and circle. This effect is more evident in replicas 1, 4 and 8 and to a lesser extent in replica 7, and it is almost entirely absent in replica 10. The areas of deformation match the arrangement of the protein complex with the representative structure of replica 1 showing a structure with the K-Ras4B and c-Raf of the two protomers side-by-side, interacting with the same membrane patch, resulting in a single area of deformation. Oppositely, the representative structure of replica 4 features an arrangement, where the inter-protomer interface is formed by the K-Ras4B protein of one protomer and the K-Ras4B and c-Raf proteins of the other, forcing the two c-Raf protomers to be distant from each other, matching the two areas of deformation shown in the mean curvature plot.

Convex (red) and flat (white) areas, are not as extensive, and in the case of the convex areas, do not result in the same magnitude of deformation, when compared with the protein-induced concave areas (blue). Conclusively, K-Ras4B dimerization induces mostly concave curvatures with respect to the pure lipid membrane bilayer, with a different degree of bending depending on the nature of the dimer interface (RAS-RAS, RAS-RAF, RAF-RAF).

## DISCUSSION

In the present work, we present the free energy landscape of K-Ras4B dimerization with or without the RBD and CRD domains of its effector protein c-Raf in the WT and G12D mutant states using the advanced free-energy technique of coarse-grained metadynamics simulations.

Initially, we performed unbiased simulations that revealed limited direct interactions between K-Ras4B protomers, specifically highlighting the absence of a K-Ras4B-KRas4B protein-protein interface. Most inter-protomer interactions across all representative structures occurred either between the c-Raf [RBD-CRD] subunits or between the K-Ras4B and c-Raf [RBD-CRD] subunits (Figure 2), with only two out of the 17 representative structures involving K-Ras4B interfaces. This result was initially unexpected, as it challenges the widely accepted view that K-Ras4B forms homodimers, which act as a scaffold for c-Raf kinase dimerization;^19,41,45^ however, upon closer examination of the recent literature,^18,27,28,31,33–35^ it is apparent that an increasing body of evidence suggests that other schemes for K-Ras4B dimerization exist.

Simulations without c-Raf [RBD-CRD] effectors identify existing interfaces and additionally reveal a plethora of novel interaction modes that the systems can access regardless of mutational status or structure toward which they are being biased.

The minima identified in these simulations highlight the fact that the three simulations share part of the conformational landscape they explore, as indicated by the presence of the GMA interface among the top-ranking minima of the WT and G12D simulations (see Figures 3b and S9-S11), but also lower ranked minima (see Figure S11 and inset).

The simulations with K-Ras4B effectors reveal that when c-Raf effectors are bound to K-Ras4B, the complexity of the derived conformational landscapes increases significantly (compare Figures 3a and 4a). The simulations explore parts of the landscape which have been previously characterized. However, these known structures present shallower minima compared to the simulations without effectors, and the K-Ras4B with effectors system is more rugged and features more distinct minima compared to the K-Ras4B without effectors system, suggesting a wider range of interactions. Interestingly, the minima of the WT simulation with effectors resemble known interfaces more frequently than those of the equivalent simulation without effectors (29% vs 43%), while the opposite is true for the G12D systems (43% vs 30%, see Figure S9), leading us to hypothesize that the presence of effectors pushes the WT system to sample more high-affinity interfaces, in contrast to the G12D simulation with effectors that shows the least similarity between its minima. Importantly, our results show that interaction modes for simulations with effectors may not only be through the direct interaction of K-Ras4B monomers, but also through the RAS-RAF and RAF-RAF interfaces. The fact that the majority of inter-protomer interfaces involve RAS-RAF or RAF-RAF interactions for all simulations with effectors, demonstrates the significance of these interactions for the multimerization landscape of this system (see below).

Analysis of the contact network of all free energy minima revealed novel interfaces in which residues of the α4-α5 or the GMA interface participate, thus resulting in the identification of additional dimerization interfaces beyond the proposed α4-α5 / GMA dimerization interfaces (Figure 6). This finding suggests that residues, which have been previously proposed to play a key role in the dimerization of specific interfaces, may in fact participate in multiple dimerization interfaces. Another key observation was the multitude of RAS-RAF interactions, a finding which is in agreement with other studies that have honed in on the role of RAS-RAF interactions.^44,46^

The calculation of binding affinities for the dimer structures reveals that simulations without effectors reach fewer but deeper (higher-affinity) minima, compared to simulations with effectors, which yield fewer high-affinity (Kd < 500 nM) minima. The high-affinity dimers primarily resemble known interfaces, such as the α4-α5 interface from PDB ID 5VQ2^36^ and the β interface seen in the inhibitor-bound PDB ID 6GJ7^22^ (Figure S19). However, these nanomolar interfaces also include previously unidentified interfaces such as minima “C” in the WT simulations without effectors and “D” in those with effectors. A model of the nanomolar interfaces of K-Ras4B dimers with and without effectors can be seen in Figure 8, which clearly shows the diversity of high-affinity interactions that K-Ras4B is capable of. Despite these high-affinity cases, most of the minima across all simulations occupy a lower-affinity region of the landscape, suggesting a variety of interchangeable dimerization interfaces. Perhaps this finding is unsurprising given the fact that protein–protein interactions that are fundamental to most cellular processes are often of surprisingly low affinity (*K_d_* values in the mM to μM range), such as the low-affinity complexes and transient complexes between proteins involved in electron transfer, enzyme–substrate complexes, and weak protein self-association.^47^ Weak, non-specific interactions may contribute very little to the free energy of the system when considered only on a pairwise basis. However, in physiological environments these interactions may collectively contribute significantly to the total free energy of the system in a highly configuration-dependent manner, thereby substantially modulating the kinetics and/or equilibria of a variety of biochemically important reactions.^43^ In fact, the versatility of K-Ras4B interfaces may stem from the fact that K-Ras4B has multiple heterodimer partners such as SOS1, RAF, PDEδ, Grb2, PIP5K1A and itself.^48^ The main advantage of using non-specific interactions to bind K-Ras4B dimers is that they can enable controlled dimerization in different conformations by achieving simultaneous delivery of several conformations from a single system. Moreover, binding K-Ras4B with non-specific interactions may improve their diffusion, which is are crucial for their effective nanoclustering. Additionally, we find that dimerization induces membrane curvature compared to the pure lipid bilayer, and this curvature depends on the nature of the dimer interface (RAS-RAS, RAS-RAF, RAF-RAF), which may also aid in adopting different membrane orientations to facilitate oligomerization. It is in fact known that the activities of several membrane enzymes are modulated by membrane curvature, e.g. the enzyme tafazzin requires defects in the membrane, i.e. perturbations in the packing order induced by either positive or negative curvature, to enhance its catalytic activity.^49^

**Figure 8.**
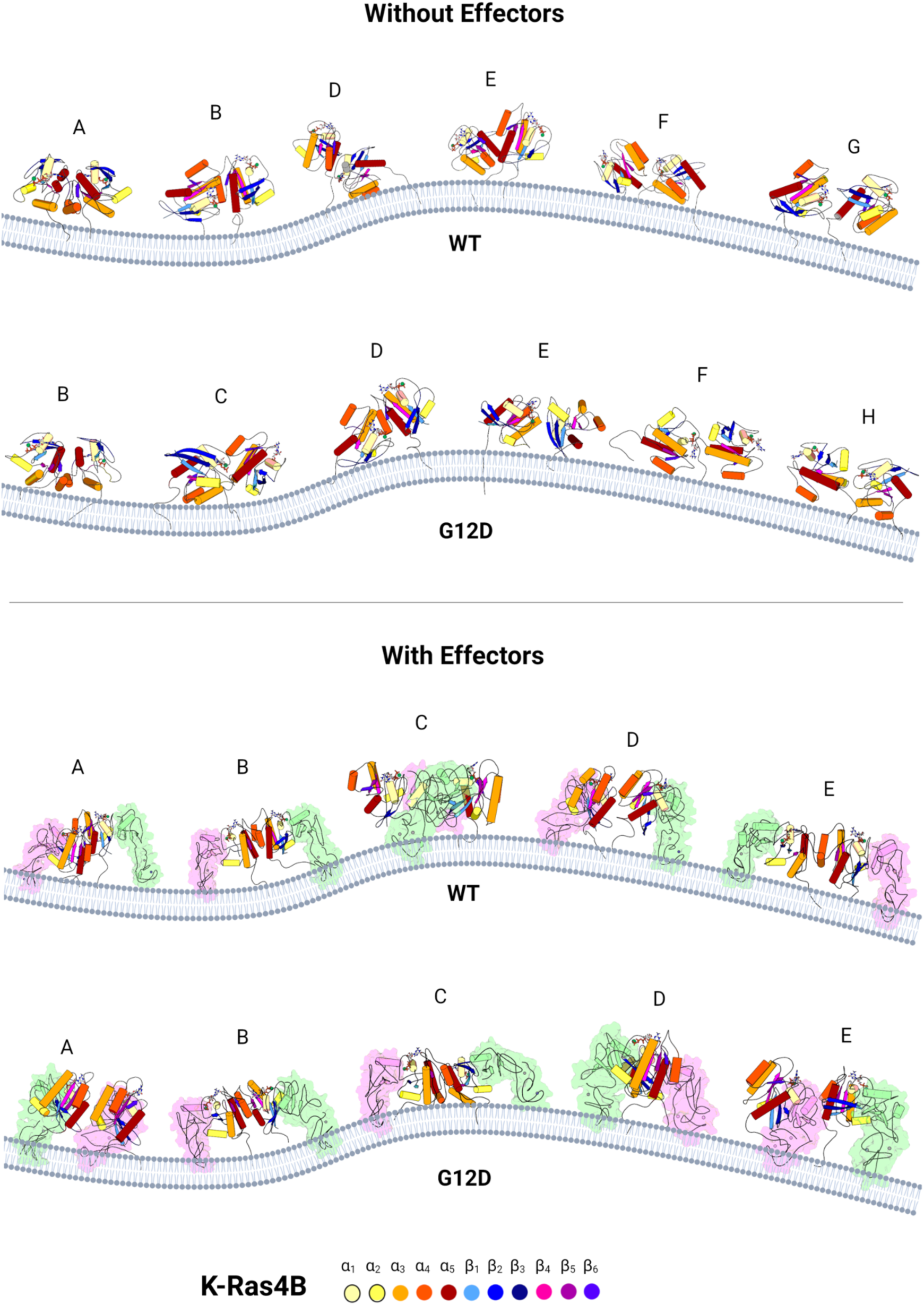
Nanomolar interfaces (Kd <500 nM) of K-Ras4B dimers with and without the c-Raf [RBD-CRD] effectors. Existing and novel interfaces are present. WT without effectors: Minimum A corresponds to PDB ID 5VQ2 (*Kd* = 10nM), B to the GMA structure (Mysore et al., *Kd* = 12 nM), E to PDB ID 6W4E (*Kd* = 60 nM) and the rest of the shown interfaces are novel. G12 without effectors: Minimum A corresponds to PDB ID 5VQ2 (*Kd* = 19 nM), D to the GMA structure (Mysore et al., *Kd* = 53nM), E to PDB ID 7ACA (*Kd* = 53 nM) and the rest of the shown interfaces are novel and below 100 nM. WT with effectors: Minima B and E corresponds to PDB ID 6W4E (*Kd* = 26 nM and 94 nM) and the rest of the shown interfaces are novel. G12D with effectors: only minimum C corresponds to PDB ID 5VQ2 (*Kd* = 98 nM) and the rest of the shown interfaces are novel and below 300 nM. For a detailed picture and *Kd*s of each model, see Figure S18. c-Raf [RBD-CRD] effectors are colored in green and pink and K-Ras4B structural elements are colored according to the labels shown in the image. The affinity values reported in the figure correspond to the highest-affinity structure of each minimum. The structure corresponds to the representative structure of each minimum.

The fact that G12D K-Ras4B mutant occupies the most diverse free energy dimer structures compared to the WT K-Ras4B may indicate that if more ways exist to complete a puzzle, the more versatile the protein becomes to perform its function, pointing into an evolutionary advantage of G12D to enable c-Raf dimerization and thus facilitate downstream signaling. These findings are corroborated by experimental results showing that weak intrinsic affinities between RAS–RAS complexes on the cell membrane can impact RAS downstream signaling^50^ and recent findings from other computational approaches, which highlight the promiscuous nature of K-Ras4B interaction profiles^33^ as well as experiments that found no evidence for reduced oncogenic KRAS signaling as a result of mutations in the proposed α4-α5 dimerization interface^27,51^. Further, a consensus arises around the hypothesis that K-Ras4B dimerization is the result of two protomers being in close proximity as opposed to actually forming a tightly bound dimer with one well-characterized interface^27,28,33^. Similarly to the results presented herein, recent reports have highlighted the likely low-affinity and non-specific nature of K-Ras4B – K-Ras4B interactions^14,27,28,52^ and have also honed in on the potential use of existing and novel biologics for targeting the K-Ras4B system.

In conclusion, our results offer unprecedented structural insights into the dimerization mechanism of K-Ras4B with and without its c-Raf effectors. The dimeric structures resolved in our study open unexplored routes for the regulation of the activity of these proteins through the structure-based design of ligands capable of modulating the formation of dimers as a novel way to prevent the downstream signaling cascades that lead to the development of cancer. These novel c-Raf-mediated interfaces of nanomolar affinity could constitute new avenues for targeting K-Ras4B in the future, as opposed to the more typical approach of targeting the c-Raf kinase domain.^53,54^

## METHODS

### Unbiased simulations of the K-Ras4B system

For the unbiased simulations, we set up a system consisting of two K-Ras4B monomers with each monomer bound to c-Raf RBD and CRD domains (henceforth referred to as c-Raf [RBD-CRD] effectors). The two K-Ras4B – c-Raf [RBD-CRD] protomers were then placed in a simulation box containing a pre-equilibrated lipid bilayer. The pre-equilibrated bilayer composition was chosen in a way that would mimic the composition of the inner leaflet of the plasma membrane as closely as possible with 1,2-Dioleoyl-sn-glycero-3-phosphocholine (DOPC): 1,2-dioleoyl-sn-glycero-3-phospho-L-serine (DOPS): Phosphatidylinositol 4,5-bisphosphate (PIP_2_) represented at 75:20:5% mol, across both leaflets. The two protomers were then anchored to the same side of the bilayer via the farnesylated C-terminal cysteines of K-Ras4B, and laterally translated along the XY plane of the bilayer, so that the distance between their closest beads was no less than 4 nm. We then simulated the system across ten independent replicas for 10 μs per replica, for a total simulation time of 100 μs across all replicas.

### Coarse-grained metadynamics simulations

#### Generation of starting conformations

The systems without c-Raf [RBD-CRD] effectors contained only two K-Ras4B monomers, whereas the remaining three systems (see Results, systems 4-6), in addition to the K-Ras4B monomers, also contained the RBD and CRD domains of the c-Raf kinase along with the flexible linker between the two domains (c-Raf [RBD-CRD] effectors). The generation of the starting conformation for the systems has been described in detail previously^32^, but a brief recap is in the SI.

#### Coarse-graining of the initial conformations to Martini 3

The all-atom starting conformations were converted to the Martini 3 CG representation using the *martinize2* tool (v0.7.3)^55^. We used our recently developed mapping scheme and parameters for the C-terminal farnesylated cysteine^56^. For the systems with c-Raf [RBD-CRD] effectors, the K-Ras4B – c-Raf [RBD] interface was maintained in its starting conformation through the elastic network of restraints created during the coarse graining of the initial all-atom coordinates. We used the default value for the elastic network potentials (500 kJ/mol/nm^2^, distance cut-off 0.9 nm), and applied it on each protomer individually, meaning the elastic network only constrained the relative movement of each c-Raf protein with respect to its bound K-Ras4B protein. For the systems without c-Raf [RBD-CRD] effectors, the elastic network was applied to each K-Ras4B protein separately, and for the systems with c-Raf [RBD-CRD] effectors, the elastic network was applied to the individual K-Ras4B proteins and their associated c-Raf proteins. We also enabled side-chain fixes. Secondary structure was assigned prior to coarse-graining, with DSSP^57,58^. We manually defined restraints between the GTP cofactor, Zn^2+^ and Mg^2+^ ions, and their coordinating residues, to prevent them from moving away from their starting positions and into the solvent during the simulation (GROMACS bond potential type 6, same force as for the elastic network). The restraints active on the GTP molecules and Mg^2+^ ions of both K-Ras4B monomers, and the Zn^2+^ ions of c-Raf can be seen in Tables S1 through S3. Martini 3 parameters for GTP were shared by Paulo C. T. de Souza in a private communication.

#### Generation and equilibration of the lipid bilayer

We used the INSANE^59^ tool (v1.1.dev0) to generate the initial lipid bilayer structure with with the same composition as in the unbiased simulation in both leaflets. For details on the construction and equilibration of the bilayer see the SI.

#### Generation of the production system

The last frame from the bilayer equilibration and the CG structures of the protein starting conformations were manually combined in PyMOL^60^ by placing the proteins on top of the equilibrated bilayer in such a way that the lipid tails of the farnesylated cysteines embedded themselves in the hydrophobic core of the lipid bilayer. The entire system was then solvated with INSANE in a simulation box with approximate dimensions equal to 19*19*15 nm (X*Y*Z axes, respectively). The solvated systems were then energy minimized with the steepest descent algorithm (up to 1000 steps) using the same settings as previously for the bilayer. For the equilibration, we only included enough Na^+^ ions to counterbalance the negative charge of the bilayer and protein components. The rest of the simulation settings remained the same as during the bilayer equilibration, with the exception of the barostat, for which we switched to Parrinello-Rahman^61^ with τP equal to 12 ps and compressibility values equal to 3 x 10^-4^ bar^-1^. All production simulation runs were carried out with GROMACS (v2021.5) patched with PLUMED (v2.8.0)^62–64^. We also simulated three independent replicas of the bilayer in the absence of any proteins using the same simulation conditions as for the production runs in the presence of the proteins. These were used to compare the diffusion coefficient of the bilayer lipids and membrane curvature in the presence and absence of the K-Ras4B-c-Raf protein complex.

#### Collective variables

We used two CVs: Root Mean Square Deviation (RMSD) from a given reference structure and K-Ras4B intermonomer distance. For the 5VQ2-based WT and G12D mutant systems with and without effectors, the PDB structure against which RMSD values were calculated was the CG model based on PDB entry 5VQ2. For the two WT systems (with and without effectors) based on the GMA interface, RMSD comparisons were carried out against the CG starting model based on the GMA interface. RMSD values for all systems were calculated over the backbone beads of all residues in stable secondary structure elements (helices and strands, as determined from the 5VQ2 structure) in the G domain of the K-Ras4B protein. Intermonomer distance was calculated between the centers of geometry (CoG) of the backbone beads of the G domain of both K-Ras4B monomers. Both CVs are physical units so the smallest value they can obtain is zero but we also soft-capped the upper value of the RMSD CV at 4.5 nm, as well as the lower and upper value of the inter-monomer distance CV between 1 and 6.5 nm, respectively, to prevent the systems from fully dissociating during the simulations.

#### CG parallel tempering metadynamics in the well-tempered ensemble

For the PT-MetaD-WTE simulations, we used eleven replicas over a range of 20 K (from 300 to 320 K with a step of 2 K), with exchanges between neighboring replicas attempted every 100 steps. During the metadynamics runs, energy, in the form of gaussian distributions, was deposited in the system every 100 steps (gaussian deposition frequency) with an initial gaussian height of 0.1 kJ/mol. The width of the deposited distributions for both CVs for all systems was set to 0.03 kJ/mol and the bias factor set to 2. The grid spacing for both CVs was set to 0.01 nm for the 5VQ2-based WT and G12D mutant systems without c-Raf [RBD-CRD] effectors and 0.006 nm for the remaining four systems. This difference had no impact on the outcome of the simulations and only affected the computational efficiency with which the bias and CVs were calculated at each step. The grid extended from 0 and 1 to 10 and 10 nm for the RMSD and intermonomer distance CVs, respectively.

### Analysis of production trajectories

Standard GROMACS tools (‘gmx trjconv’) were used for the treatment of all trajectories and their post-processing (removal of periodicity effects, centering, clustering and extraction of representative structures). All analyses for all metadynamics and unbiased simulations were carried out after 15 and 10 μs (for all ten replicas) of simulation, respectively.

#### Interface RMSD calculation for the unbiased simulations

The interface RMSD values were computed over the trajectories of all ten independent replicas from the unbiased simulations, using the CG initial structures for the 5VQ2-based and GMA-based PT-MetaD-WTE simulations as reference structures. The comparison was carried out on the backbone beads of the interface residues. The interface was defined, in accordance with the criteria used for the assessment of the accuracy of predicted protein-protein interfaces^65^, as the set of K-Ras4B monomer residues whose atoms are within 1 nm of any atom of the other K-Ras4B monomer. For more information, see the SI.

#### Clustering analysis for the unbiased simulations

We performed a clustering analysis for each of the ten independent replicas and identified the major clusters for each trajectory, from which we extracted representative structures. We used the GROMOS clustering algorithm^66^ as implemented in GROMACS (‘gmx cluster-method gromos’) with a cut-off value of 1 nm. The RMSD values for the clustering were computed using all backbone beads of K-Ras4B (minus the HVR residues) and c-Raf [RBD-CRD] effectors. Clusters are numbered according to cluster size, with the first cluster being the most populated one and the last cluster the least populated.

#### Comparison of unbiased simulation frames against representative structures

Using the same settings as for the clustering analysis above, we calculated the RMSD of all simulation frames against the representative structure(s) of each replica, for all replicas. The results of the analysis are shown in Figure S2.

#### CG PT-MetaD-WTE convergence monitoring CV diffusion, 1D and 2D free energy calculations

The convergence of the CG PT-MetaD-WTE simulations were monitored through the diffusion of CVs over time and the amount of energy being deposited into the system. CV diffusion was monitored by dumping CV values (RMSD and intermonomer distance) every 2 ps in a text file through PLUMED. The values presented in Figure S4 were obtained every 200 ps.

The ‘sum_hills’ utility of PLUMED was used for the calculation of the 1D free energy profile as a function of both CVs (RMSD from reference structure and K-Ras4B intermonomer distance) as well as for the 2D free energy landscape (RMSD vs intermonomer distance). These calculations were carried out for the entire trajectory including increasingly bigger parts (in steps of 500 ns), until the entire trajectory was included in the analysis. All calculations were carried out on the replica with the physiological temperature (310 K). The last few microseconds (12.5 – 15 μs) of each trajectory were used to quantify the convergence of the simulations through the RMSD-based free energy profiles (see Figure S6). 2D free energy landscapes were generated at 15 μs.

#### Identification of minima from the 2D free energy landscapes

We processed the 2D free energy landscape, which was calculated from the full trajectory for each system, using an in-house implementation of the following algorithm:

1. Identification of the 2D (RMSD, intermonomer distance) coordinates of the lowest energy value. This point corresponds to the global minimum of the free energy landscape for each system.
2. Proceed to the point with the next highest energy and identify its coordinates.
3. Compute the 2D Euclidean distance (in units of RMSD and intermonomer distance) between this point and all previously identified minima. For the first iteration of the algorithm, only the distance between this point and the global minimum is computed, since no other minima have been identified.
4. If this point is within a predetermined 2D distance cut-off of any previously identified minimum (including the global minimum), then this point is ignored since it is determined to be too close to a previously identified minimum.
5. If this point is not within a predetermined 2D distance cut-off of any previously identified minimum, then this point is identified as a new minimum of interest.
6. Steps 2 through 5 are repeated until all relevant minima below the desired energy cut-off value (−6 kcal/mol) have been sampled.

After testing various values for the distance cut-off of the aforementioned algorithm, we eventually settled on using a 2D distance cut-off of 2 units, assuming both RMSD and intermonomer distance are in Å.

Following identification of all relevant minima, we then analyzed each trajectory with PLUMED, computing RMSD and intermonomer distance CV values for each frame of the trajectory written to disk. To isolate representative structures for all minima, we filtered all trajectory frames by RMSD and intermonomer distance values and identified the frames that belonged in each minimum. We used the same clustering settings as for the analysis of the unbiased simulations (see “Clustering analysis for the unbiased simulations” section in Methods).

To ensure the identified minima represented relevant and non-redundant parts of the conformational landscape we carried out two additional rounds of filtering. We first calculated the buried surface area (BSA)^67^ for all coarse-grained structures identified by the above algorithm so as to only include complexed structures in the subsequent analyses. Subsequently, we also cluster the structures and selected representative entries from each cluster to remove redundant structures from the analysis. Information on these calculations can be found in the SI.

#### Comparison of minima with PDB entries and unbiased representative structures

In order to carry out comparisons with atomistic structures of K-Ras4B complexes deposited in the PDB, we backmapped the CG models to atomistic resolution using the ‘CG2AT2’ tool.^68^ All structural comparisons between the identified minima structures and PDB entries and unbiased simulations representative structures, were performed with the program ProFit (Martin, A.C.R., http://www.bioinf.org.uk/software/profit/), which implements the McLachlan fitting algorithm.^69^ We used the alpha carbon atoms of the subset of residues that were resolved in the 5VQ2 structure for these comparisons (residues 2-33, 37-58 and 69-166). Overall, we used PDB entries 5VQ2, 6W4E, 7rse^25^, 3KKN^70^, 6GJ7^22^, 6QUV^71^, 7ACA/F/Q^24^ and 7EW9^42^ and the GMA structure as reference structures in these comparisons. For the crystallographic entries of interest that were deposited in the PDB as monomers, we reconstructed the dimers after applying crystallographic symmetry operations and selecting the most viable dimers, following their visual inspection. The dimers that were reconstructed this way are those with PDB identifiers 3KKN and 6GJ7. For 3KKN, we reconstructed the dimer in a way that recreated the α3-α4 interface and for 6GJ7 in a way that recreated the small-molecule-stabilized β-β interface. We also selected the small-molecule-stabilized interface for entry 7acq. We used the same subset of residues (residues 2-33, 37-58 and 69-166) of K-Ras4B for the comparison of the minima structures and representative structures identified in the unbiased simulations.

We assigned the same similarity classes for all comparisons (against PDB entries and unbiased simulation representative structures) as for the analysis of the unbiased simulations, using the same RMSD cut-off values (see “Similarity analysis for the unbiased simulations” section in Methods).

#### Calculation of inter-protomer contacts and contact class analysis

We calculated inter-protomer contacts using the CG structures of the minima previously identified to avoid the ambiguity introduced by the placement of the side-chain atoms during the backmapping procedure, using a distance cutoff of 0.5 nm. We excluded all contacts involving the HVR residues (residues 167-185) of K-Ras4B due to their flexible nature, all ions (Mg^2+^ of K-Ras4B and Zn^2+^ of c-Raf) and the GTP cofactor. After calculating all contacts, we classified each in one of three categories: RAS-RAS, RAS-RAF and RAF-RAF, depending on which subunit the residues involved in the interaction belonged to.

For the creation of the contact maps, we used a longer distance cutoff equal to 1.2 nm, to capture more distant contacts in the minima structures. First, we determined the contacts of all residues and calculated the number of minima in which each contact was present. For the calculation of contact maps for the GMA hexadecamer, we used the supplementary structure Mysore et al.^26^ made available and using the same residue ranges as for the simulated systems and a distance cut-off value equal to 1 nm, since this is an atomistic system. We did the same for the dimeric GMA complex that was used as the biasing structure for the WT-GMA simulations and the structure of PDB entry 5VQ2 which was the biasing structure for simulations WT and G12D.

#### Transformation of free energy to binding affinity values

We transformed energy values obtained from the 2D free energy landscapes (see Figures 3a, 4a, S7 and S8) to binding affinities using the following equation:

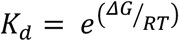

Where K_d_ indicates the binding affinity, ΔG the binding free energy, R the Boltzmann constant and T the simulation temperature (310 K). Binding affinity values were calculated for each clustered frame and the median as well as the total range of values are shown in Figure 7.

#### Radial distribution function analysis

We calculated the radial distribution functions with the rdf module of GROMACS (‘gmx rdf’) using two sets of beads as reference positions: the backbone bead (BB) of threonine 183 of K-Ras4B, and the terminal side chain bead (SC2) of lysine 148 of c-Raf. The latter calculation was only carried out in the simulations with c-Raf [RBD-CRD] effectors, whereas the former in both simulations, with and without effectors. In both calculations we used the phosphate beads (PO4) of the three lipids, for the positions of the lipid heads. The lipids were further limited so as to only include lipid heads of lipids belonging to the leaflet with which the protein complex was interacting (upper leaflet). The number of lipids presented in Table 2 were calculated after integrating the area under the distributions past the second shell, at a distance cutoff of 1.1 nm. We used the PT-MetaD-WTE simulations for this analysis.

#### Diffusion coefficient calculations

The mean squared displacement module of GROMACS (‘gmx msd’) was used for the calculation of the lateral displacement with the time between the reference points for the calculations set to 10 ns. We used the center of mass of the full K-Ras4B-c-Raf complex and that of the individual lipids for the displacement calculation, both for the monomeric and unstable dimeric states. The displacement was then averaged across the number of molecules: two in the case of the protein complex, 792, 210 and 52 for the DOPC, DOPS and PIP_2_ lipids, respectively. After obtaining the MSD vs time plots, we fit a first-degree equation in the linear regime of each region of interest for each plot and extracted the slope. The diffusion coefficient was calculated with the following equation:

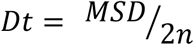

Where D is the diffusion coefficient, MSD is the slope determined from the linear fit described above and n is the number of dimensions in which the calculations were performed, in this instance 2. This analysis was performed on the unbiased simulation replicas 1, 6 and 8 (simulation time between 1 and 1.5 μs) for the calculation of the diffusion speed of the monomeric states, and replicas 2 and 3 (simulation time between 1 and 2 μs) for that of the unstable dimeric state.

#### Membrane curvature analysis

We used the MembraneCurvature^72^ module of MDAnalysis^73,74^ to analyze the curvature of the bilayer in the presence and absence of the K-Ras4B-c-Raf protein complex. We first separated the bilayer into two leaflets based on the Z-axis coordinates of the phosphate beads of all lipids. For the simulations with the protein complex present, the upper leaflet of the bilayer was the one associating with the protein complex. We then divided each leaflet into 100 square cells with approximate X and Y dimensions equal to 1.9 * 1.9 nm, respectively (total bilayer size ∼ 19 * 19 nm). With this 10 * 10 cell grid in place, we computed the average mean curvature of each cell, using the coordinates of the phosphate beads of all lipids contained within the cell. The resulting grid colormaps are shown in Figure S28.

As part of this analysis, we also computed the coordinates of the center of mass of the protein complex and its radius of gyration. We used GROMACS for both calculations, utilizing the trajectory (‘gmx traj’) and radius of gyration (‘gmx gyratè) modules, respectively. We used the entire protein complex for both calculations and only considered the radius of gyration around the Z axis (parallel to the membrane normal). Both analyses were carried out on replicas 1, 4, 7, 8 and 10 of the unbiased simulations and after stable dimeric complexes had formed (simulation time ≥ 4 μs). The entire production trajectories of all three replicas were used for the bilayer simulations in the absence of any proteins.

## DATA AVAILABILITY

All minima structures at the coarse-grained and all-atom resolution levels are deposited in Zenodo (https://doi.org/10.5281/zenodo.14169550, structures.zip). The Zenodo archive also contains input files for all simulations presented herein, as well as trajectory files and code/scripts used for the analyses carried out.

## ACKNOWLEDGMENTS

This work was supported by computational time granted from the Greek Research & Technology Network (GRNET) in the National HPC facility ARIS under Project ID pr012011/CG-KRAS, granted to ZC and PK. ZC and PK also acknowledge PRACE for granting computational time in Marconi100, CINECA, Italy.

## AUTHOR INFORMATION

### Author Contributions

ZC, CVV and PIK designed and supervised the study. PIK carried out simulations and performed calculations and analysis. PIK wrote the first draft. All authors analyzed and discussed the results and implications, wrote, revised, and approved the main text and the Supplementary information, discussed and evaluated the progress of the project.

