## Supporting Information for "Mapping the free energy landscape of K-Ras4B dimerization"

### Methods

#### Coarse-grained metadynamics simulations

##### Generation of starting conformations

For the WT system without effectors, we generated the K-Ras4B monomers from the 5VQ2 PDB entry, carrying out all the required mutations to conform to the WT sequence. We made use of additional PDB entries 4g0n[1] and 2msc[2] to model the switch II, and HVR and switch I regions of the protein, respectively. The terminal cysteine was farnesylated using the parameters available as part of the CHARMM36m force field.[3] For the WT system with effectors present, we modelled the c-Raf domains using the 4g0n and 1far[4] PDB entries as the basis for the RBD and CRD domains, respectively. We also modelled the flexible linker between them with a combination of 2-body docking and loop modelling, using the AIDA webserver.[5] For the equivalent G12D systems, we mutated residue 12 of both K-Ras4B monomers from glycine to glutamic acid. For the GMA systems, the monomers were generated in the same way as for the WT systems, but the starting orientation of the K-Ras4B monomers was changed after superimposing each monomer on the respective GMA interface monomer. The K-Ras4B monomers for all systems were loaded with GTP and $Mg^{2+}$ in the GTP binding pocket, and the CRD domain of c-Raf included two $Zn^{2+}$ ions.

##### Generation and equilibration of the lipid bilayer

The initial dimensions of the bilayer and its simulation box were set to 18 * 18 * 12 nm, along the X, Y and Z axes, respectively. We only included enough $Na^+$ ions to counterbalance the highly negative charge of the lipid bilayer, a result of the presence of $PIP_2$ lipids. The system was energy-minimized with the steepest descent algorithm for up to 1000 steps, followed by approximately 15 μs of equilibration. Velocities were generated and kept at 310 K for the entire equilibration run through the use of v-rescale thermostat[6]. Pressure target was set at 1 bar via a semi-isotropic barostat (Berendsen[7] with $\tau_P$ equal to 4 ps and compressibility values equal to 4.5 x $10^{-5}$ $bar^{-1}$). We used the recommended settings for the Martini 3 force field non-bonded interactions, i.e., a 1.1 nm cut-off for short-range electrostatics and van der Waals interactions, a reaction field for the electrostatics with a screening constant of 15 (set to 0 beyond the short-range cut-off) and shifting the van der Waals potential to 0 at the cut-off (with the Verlet cut-off scheme[8]). A 20 fs time step was used throughout. We used GROMACS (v2021.5) for the equilibration.[9,10]

**Interface RMSD calculation for the unbiased simulations**

We used the all-atom initial structures (prior to their coarse graining) for the calculation of the atom-atom distances. Atoms from the HVR were excluded from this due to their ability to form contacts with distant parts of the binding partner as a result of the highly flexible nature of the HVR. This yielded residues 2-4, 45-51, 82, 84, 108-116, 118-119, 123, 125-145 and 150-166 for the 5VQ2 interface and residues 10-18, 27-36, 57-64, 68, 82-92, 96, 116-125, 147-148 and 22, 46-47, 82, 84, 111-159, 161-162, 164-166 for the GTP-contributing and non-GTP-contributing monomers for the GMA interface, respectively.

**Similarity analysis for the unbiased simulations**

We used the same bead selection to compute an all-vs-all RMSD matrix of the representative structures to identify potential points of convergence between the independent replicas. The RMSD criteria we used for the determination of whether two minima structures were highly, acceptably, or lowly similar or entirely dissimilar and were determined after visual inspection of the superimposed representative structures, after taking into consideration the fact that the backbone beads of the backbone of the entire protein complex (minus the HVR K-Ras4B residues) was included in the calculation. The three labels (highly, acceptably, or lowly similar) reflect different levels of similarity with the former corresponding to two representative structures where all the crucial elements of the interaction converge with a high degree of accuracy, the middle corresponding to a pair of representative structures where the overall arrangement is similar but smaller-scale interactions such as residue-residue distances or interface composition diverges to some extent, and the latter corresponding to a pair where the only similarities can be found in some of the intermonomer interactions as some domains might have shifted significantly. Any pairs whose similarity is classified as "low" exhibit no meaningful similarities. The RMSD values we have used as cut-off ranges for the similarity classes are [0, 0], (0, 5), [5, 10), [10, 15), [15, ∞) Å, for identical, highly similar, acceptably similar, lowly similar, or dissimilar structural pairs, respectively.

**Identification of minima from the 2D free energy landscapes: filtering for non-redundant minima**

The BSA was computed with GROMACS after calculating the Solvent Accessible Surface Area (SASA) for the two protomers individually and together, yielding a total of three SASAs: The SASA of the complex, the SASA of the first protomer and the SASA of the remaining protomer. SASA values were computed with the sasa module of GROMACS (`gmx sasa`) using a probe with radius equal to 0.19 nm and sampling the surface area of the structures at a density of 100 dots. The van der Waals radii for regular, small and tiny beads were set to 0.264, 0.23 and 0.191 nm, respectively, and in agreement with Martini 3 parameters[11]. The BSA is defined and calculated as:

$$BSA = (SASA_{protomer\_1} + SASA_{protomer\_2}) - SASA_{complex}$$

Where BSA is the buried surface area, $SASA_{protomer\_1}$ and $SASA_{protomer\_2}$ are the SASAs of the two individual protomers and $SASA_{complex}$ the SASA of the complexed structure. It follows from the equation above that in the cases of structures where no part of the surface of either protomer is buried to form the dimer, the sum of the SASA values of the protomers will be equal to the SASA of the complex, thus equaling a BSA of zero. We only included complexes whose BSA was equal to or greater than 350 Å$^2$. This value was determined after consulting previous literature related to the computational study of protein-protein interactions[12,13]. In both studies, the smallest interface associated with an experimentally determined dimeric protein-protein complex structure was equal to 368 and 381 Å$^2$. Therefore, the identified structures whose BSA is smaller than 350 Å$^2$ represent parts of the landscape in which the two protomers have fully dissociated (BSA = 0) or are transient states ($0 < BSA < 350$ Å$^2$) between other, fully complexed states ($BSA \geq 350$ Å$^2$).

After filtering the identified minima to only include the fully complexes states, we proceeded to cluster them and select representative entries from each cluster to remove redundant structures from the analysis. We carried out the clustering with GROMACS using the backbone beads of residues 2-166 of both K-Ras4B monomers (thus excluding the HVR residues) for all systems. We used the GROMOS method with a cut-off value of 0.5 nm. For the structures that were clustered together we selected the structure with the lowest energy/rank (as assigned from PLUMED). All representative structures and structures that formed single-entry clusters were included in all subsequent analyses.

**Table S1**: Distance restraints between GTP molecules and coordinating protein residues. Table rows indicate the GTP and K-Ras4B protein residue beads each restraint it acting on as well as the target distance for the restraint.

| GTP | | Protein | | | | |
|---|---|---|---|---|---|---|
| Chain | Bead | Chain | Residue Index | Residue Name | Bead | Distance [nm] |
| A | PG | A | 16 | LYS | SC2 | 0.417 |
| A | PB | A | 16 | LYS | SC2 | 0.382 |
| A | PA | A | 15 | GLY | BB | 0.415 |
| A | RB1 | A | 117 | LYS | SC2 | 0.513 |
| A | RB2 | A | 28 | PHE | SC2 | 0.48 |
| A | SC1 | A | 117 | LYS | SC2 | 0.41 |
| A | SC2 | A | 119 | ASP | SC1 | 0.378 |
| A | SC4 | A | 145 | SER | SC2 | 0.389 |
| B | PG | B | 16 | LYS | SC2 | 0.42 |
| B | PB | B | 16 | LYS | SC2 | 0.381 |
| B | PA | B | 15 | GLY | BB | 0.415 |
| B | RB1 | B | 117 | LYS | SC2 | 0.511 |
| B | RB2 | B | 28 | PHE | SC2 | 0.479 |
| B | SC1 | B | 117 | LYS | SC2 | 0.415 |
| B | SC2 | B | 119 | ASP | SC1 | 0.377 |
| B | SC4 | B | 146 | ALA | SC1 | 0.394 |

**Table S2**: Distance restraints between $Mg^{2+}$ ions and coordinating protein residues. Table rows indicate the $Mg^{2+}$ and K-Ras4B protein residue beads each restraint is acting on as well as the target distance for the restraint.

| $Mg^{2+}$ | Protein | | | | |
|---|---|---|---|---|---|
| Bead | Chain | Residue Index | Residue Name | Bead | Distance [nm] |
| MG | A | 17 | SER | SC1 | 0.295 |
| MG | A | - | GTP | PG | 0.343 |
| MG | A | 57 | ASP | SC1 | 0.344 |
| MG | A | - | GTP | PB | 0.348 |
| MG | B | 17 | SER | SC1 | 0.289 |
| MG | B | - | GTP | PG | 0.351 |
| MG | B | 57 | ASP | SC1 | 0.351 |
| MG | B | - | GTP | PB | 0.355 |

**Table S3**: Distance restraints between $Zn^{2+}$ ions and coordinating protein residues. Table rows indicate the $Zn^{2+}$ and Raf protein residue beads each restraint is acting on as well as the target distance for the restraint.

| **$Zn^{2+}$** | | **Protein** | | | | |
|---|---|---|---|---|---|---|
| Ion | Bead | Chain | Residue Index | Residue Name | Bead | Distance [nm] |
| 1 | ZN | A, B | 152 | CYS | SC1 | 0.23 |
| 1 | ZN | A, B | 155 | CYS | SC1 | 0.23 |
| 1 | ZN | A, B | 176 | CYS | SC1 | 0.23 |
| 1 | ZN | A, B | 173 | HIS | SC3 | 0.21 |
| 2 | ZN | A, B | 165 | CYS | SC1 | 0.23 |
| 2 | ZN | A, B | 168 | CYS | SC1 | 0.23 |
| 2 | ZN | A, B | 184 | CYs | SC1 | 0.23 |
| 2 | ZN | A, B | 139 | HIS | SC3 | 0.21 |

### Results

#### Unbiased CG simulations of two K-Ras4B monomers on a lipid bilayer reveal novel interaction modes

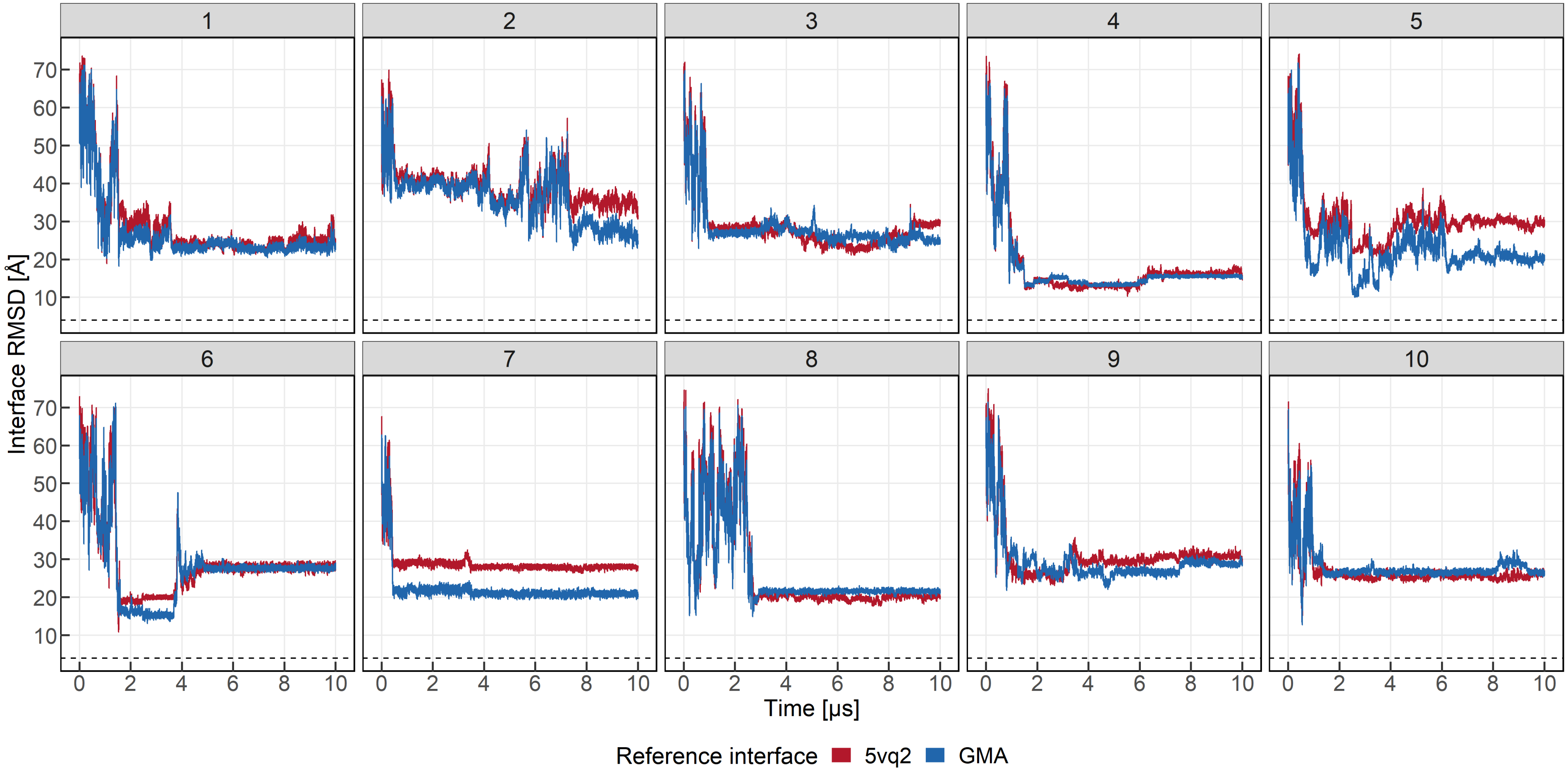


**Figure S1**. Interface RMSD of the K-Ras4B dimer when compared against the PDB entry 5VQ2 and the interface proposed recently by Mysore et al. Each box corresponds to an independent replica. The red and blue lines correspond to the interface RMSD values when using PDB entry 5VQ2 and the GMA interface, respectively. The dashed black line corresponds to an interface RMSD value of 4 Å, which signifies an acceptably similar interface.

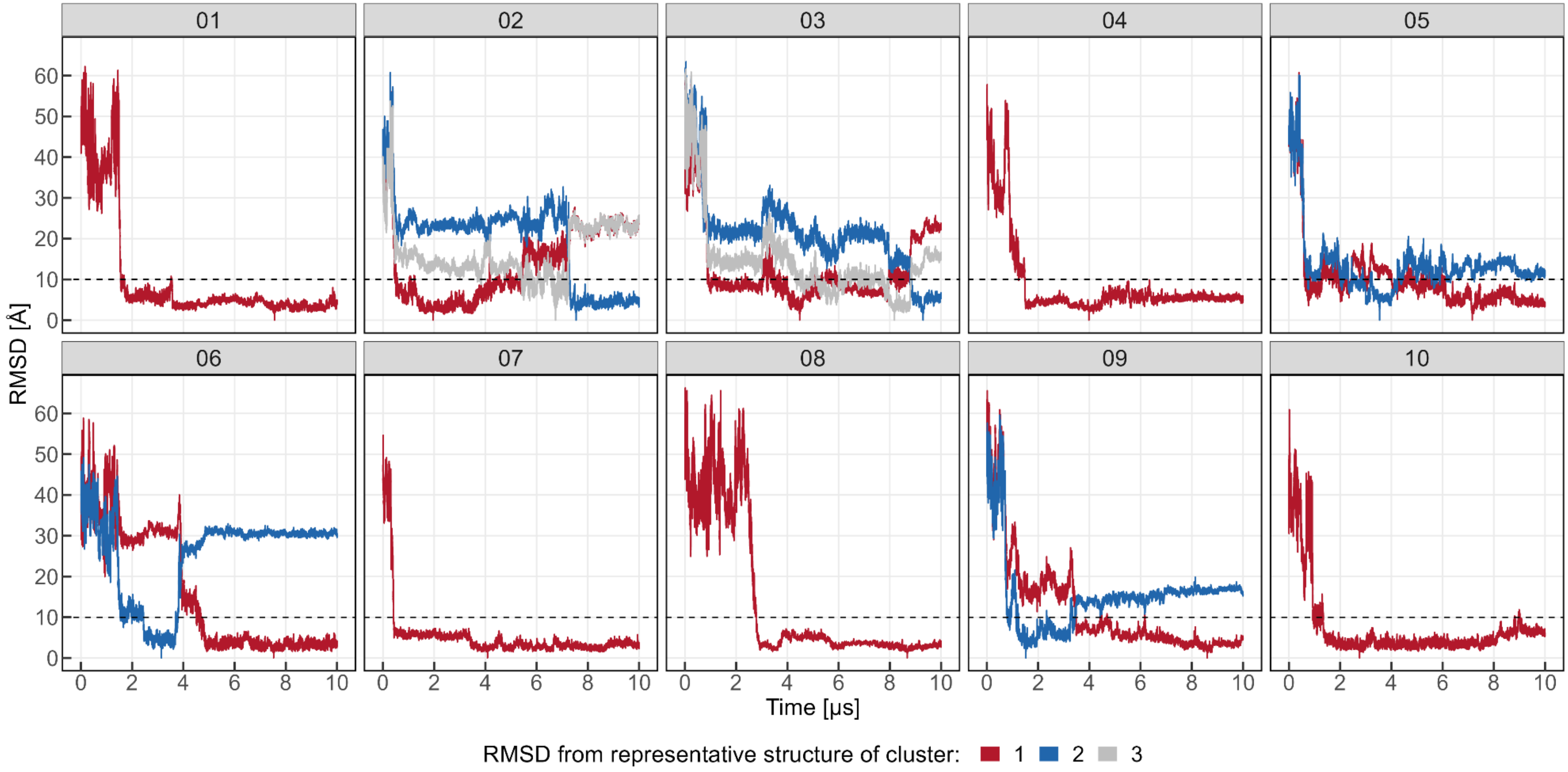


**Figure S2.** RMSD of the full K-Ras4B system (K-Ras4B + Raf [RBD-CRD] effectors) against the representative values from each replica. RMSD values have been calculated on all backbone beads of the proteins (minus the K-Ras4B HVR residues). Each box corresponds to an independent replica and the lines within each box are colored red, blue and light grey to indicate RMSD values from the first, second and third representative structure of each trajectory. The dashed black line indicates the RMSD value that was used for the clustering analysis (10 Å). The vertical lines which reach down to 0 on the Y axis indicate the timestamp of the representative structure from each major cluster.

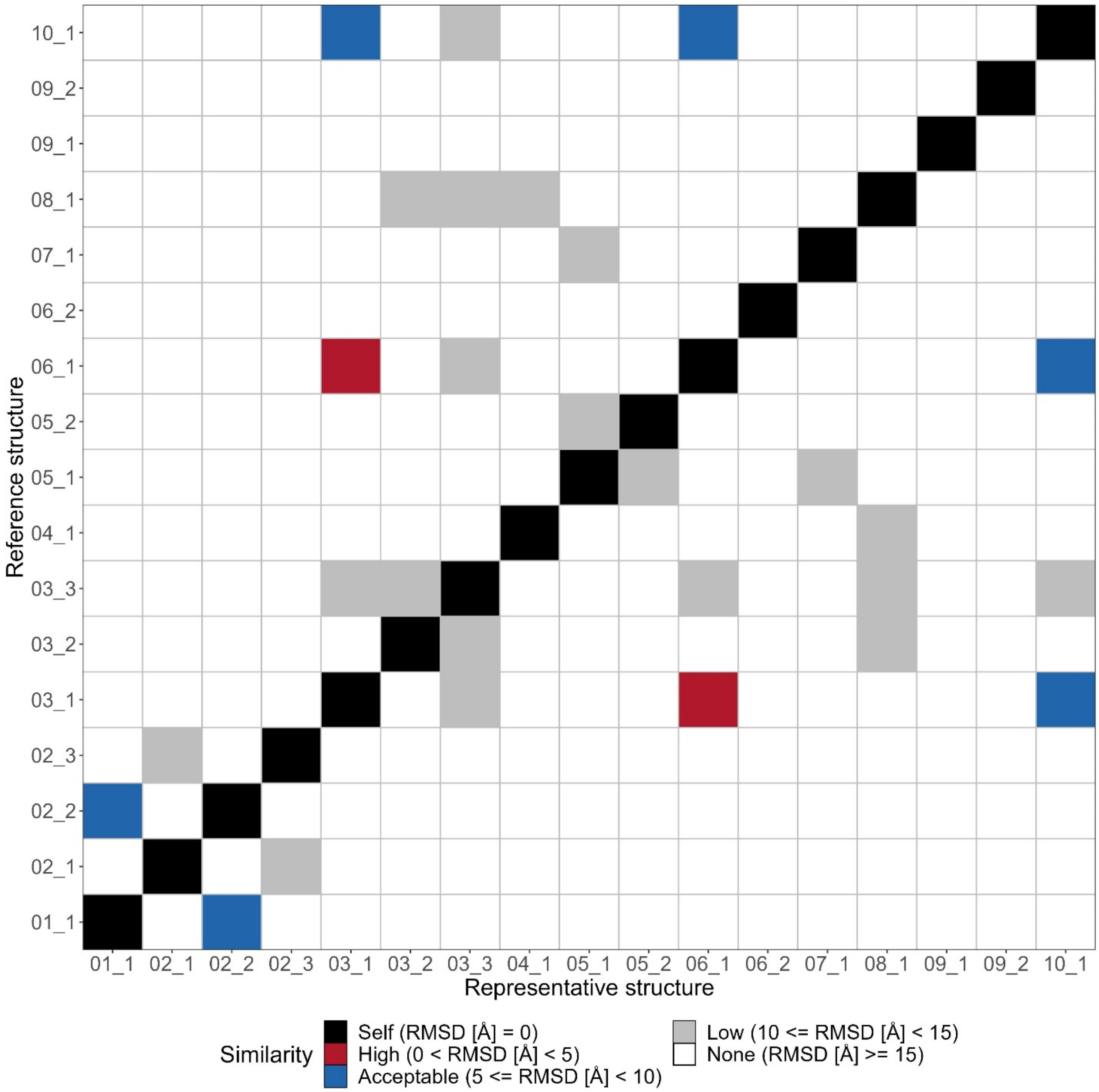


**Figure S3**. Similarity matrix of the representative structures from the clustering analysis of the unbiased simulations. Each cell of the matrix is colored according to the qualitative similarity of the two representative structures the cell corresponds to (X and Y axes). A black color corresponds to a cell with a self-comparison (RMSD = 0 Å), whereas red-, blue-, grey- and white-colored cells indicate high, acceptable, low and no similarity pairs, respectively. The RMSD ranges for the four similarity classes are (0-5), [5-10), [10-15) and [15-∞) Å, respectively, and have been computed over all backbone beads of the K-Ras4B (minus the HVR residues) and Raf proteins.

#### 2D dimerization free energy landscape of WT and G12D K-Ras4B using CG-Metadynamics simulations

##### Simulation convergence

Figure S4 shows the diffusion of both CVs over the course of the six simulations. All systems sample a broad range of values, with the observed values for the RMSD and intermonomer distance CVs ranging approximately between 0-25 and 30-70 Å, respectively. However, the range of observed values for both CVs for the GMA-based systems (both with and without effectors) is noticeably narrower than that of the other two systems, although not in a statistically significant way (see also the violin plots in Figure S5). This indicates the conformational landscape of the GMA-based systems is more constrained than that of the remaining ones. Specifically, focusing on the RMSD CV, there is a noticeably more populated segment (compared to the other two systems) below 5 Å, which corresponds to the native population of the simulation, indicating the systems favor the native state over exploring alternative states compared to the 5VQ2-based systems.

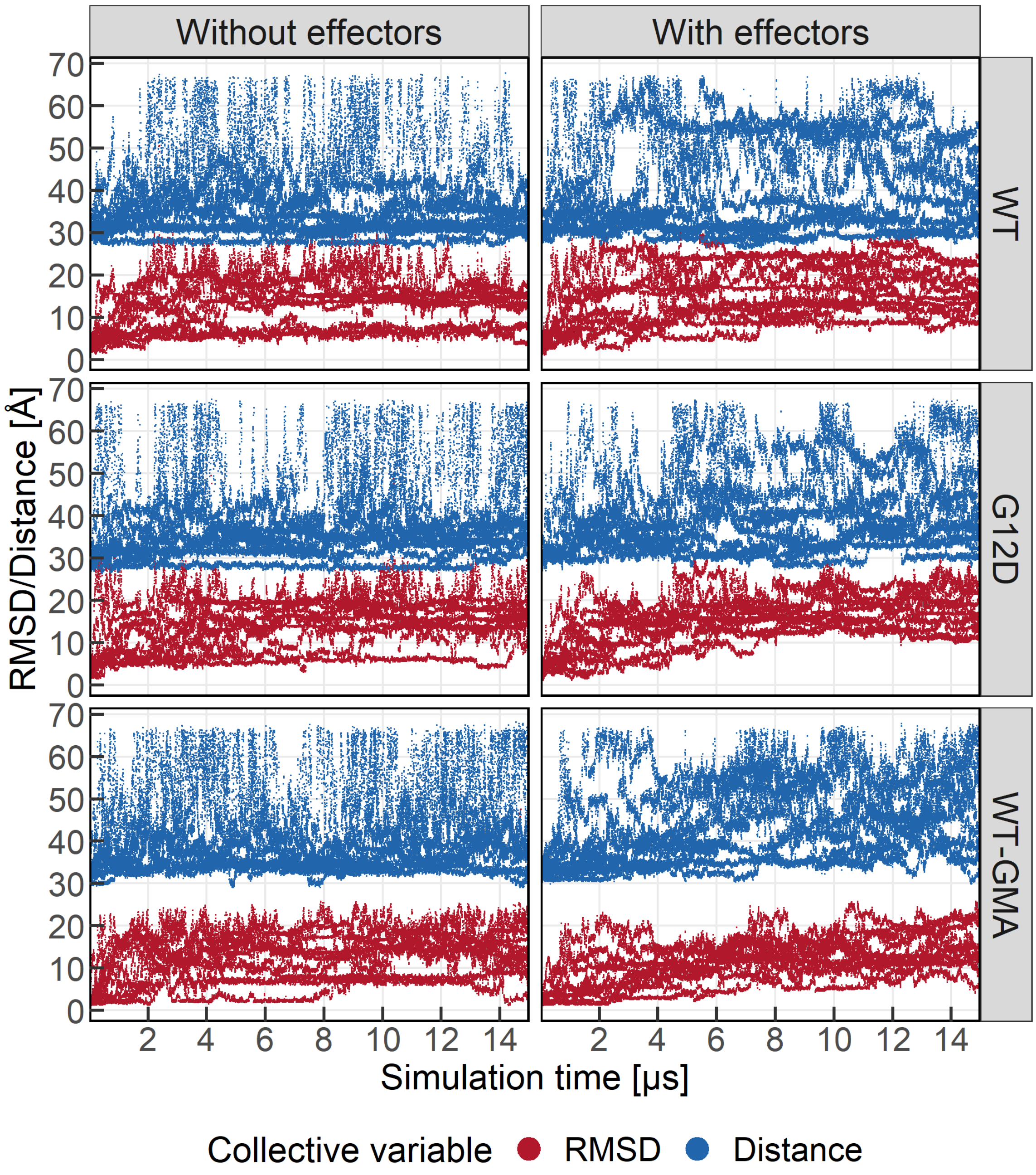


**Figure S4**. Diffusion plots for the two collective variables for all simulations. The values corresponding to the RMSD and Distance CVs are colored red and blue, respectively. The values have been grouped by presence of Raf effectors (vertically) and simulation (horizontally). RMSD/Distance values are plotted every 200 ps.

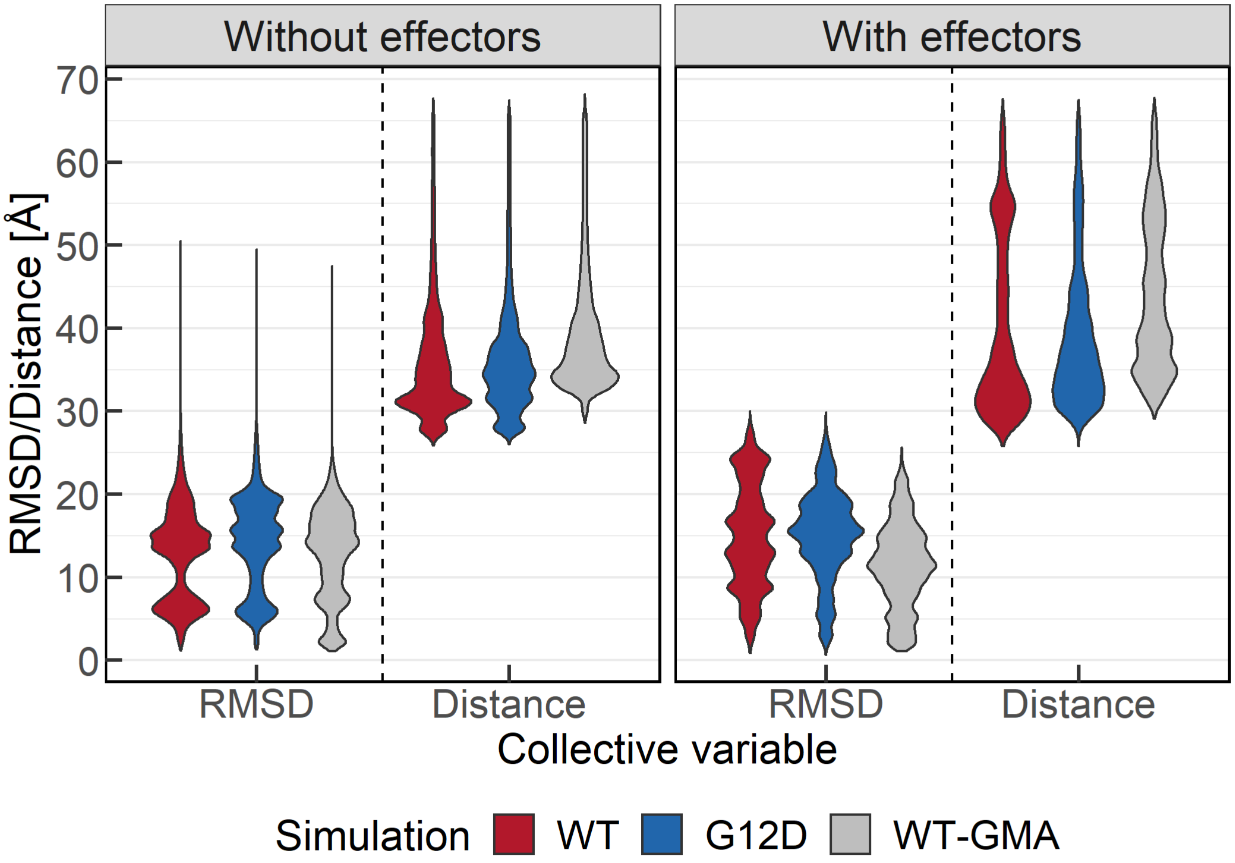


**Figure S5**. Violin plots of the CV values for all simulations. The distributions are colored red, blue and light grey for the WT, G12D and WT-GMA simulations, respectively.

The convergence of the simulations was monitored through the 1-dimensional (1D) free energy plots with respect to the RMSD CV. Figure S6 shows the 1D free energy profile for all simulations with some differences between the various systems easily noticeable. Comparing the systems with and without effectors (bottom and top row, respectively) reveals the profile of the systems without effectors to be significantly simpler with fewer minima and more pronounced sampling of the state toward which the simulation is being biased (5VQ2 for WT and G12D, and GMA for WT-GMA). This difference also extends to the convergence comparison with the systems without effectors showing no significant variations over the last few microseconds of trajectories, indicating that the simulations are converged. Another interesting observation is the comparison between the WT-GMA and remaining systems, which reveals the former mostly explored the part of the conformational landscape around the native state of the system (GMA), especially in the case of the system with effectors.

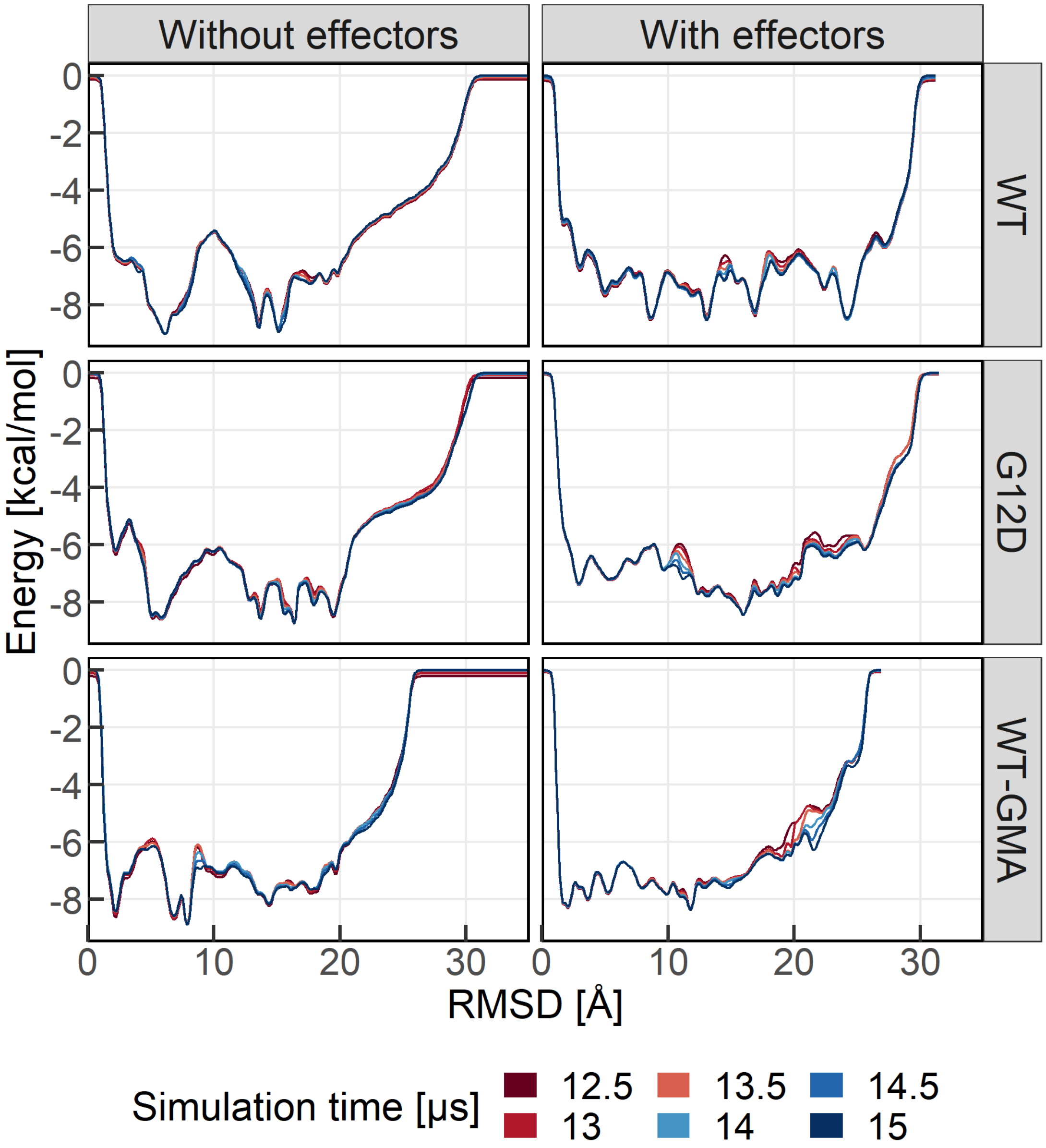


**Figure S6**. 1D free energy profile vs the RMSD CV for all simulations. The free energy has been computed up to specific simulation times (every half µs) and the last few µs of each simulation are shown ranging from 12.5 (dark red) to 15 µs (dark blue). The free energy values have been normalized according to the lowest energy observed across all systems (WT without effectors).

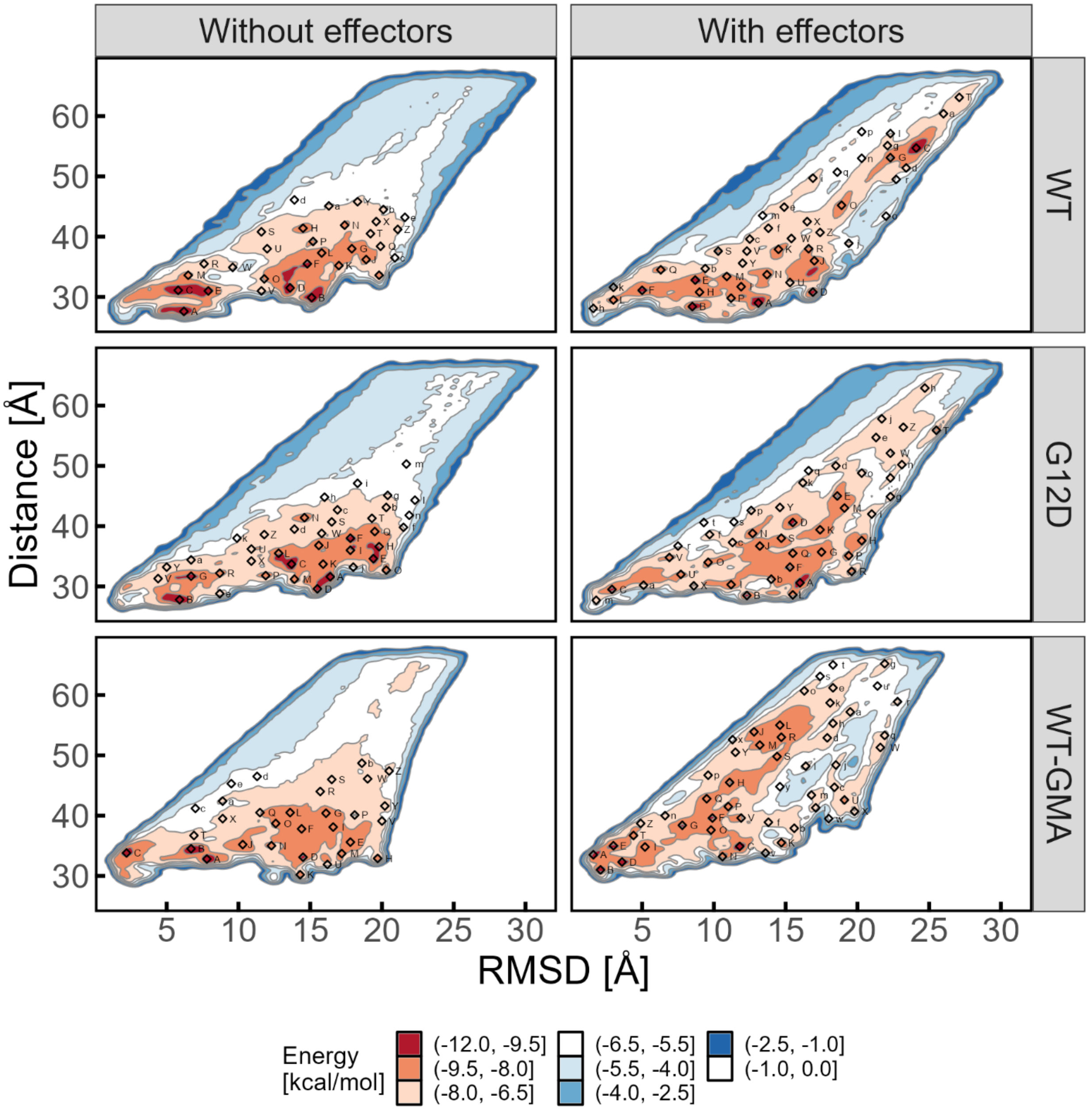


**Figure S7.** Labeled 2D free energy landscapes of K-Ras4B system. The simulations have been grouped vertically by the presence/absence of Raf effectors and horizontally by the system (WT, G12D and WT-GMA). The X and Y axes correspond to the RMSD and inter-monomer distance CV values, respectively. The coloring indicates the energy value associated with each coordinate and ranges from dark red (global minimum) through white and blue to white (0). Energy values have been normalized according to the lowest energy observed across all systems (WT-GMA without effectors, global minimum).

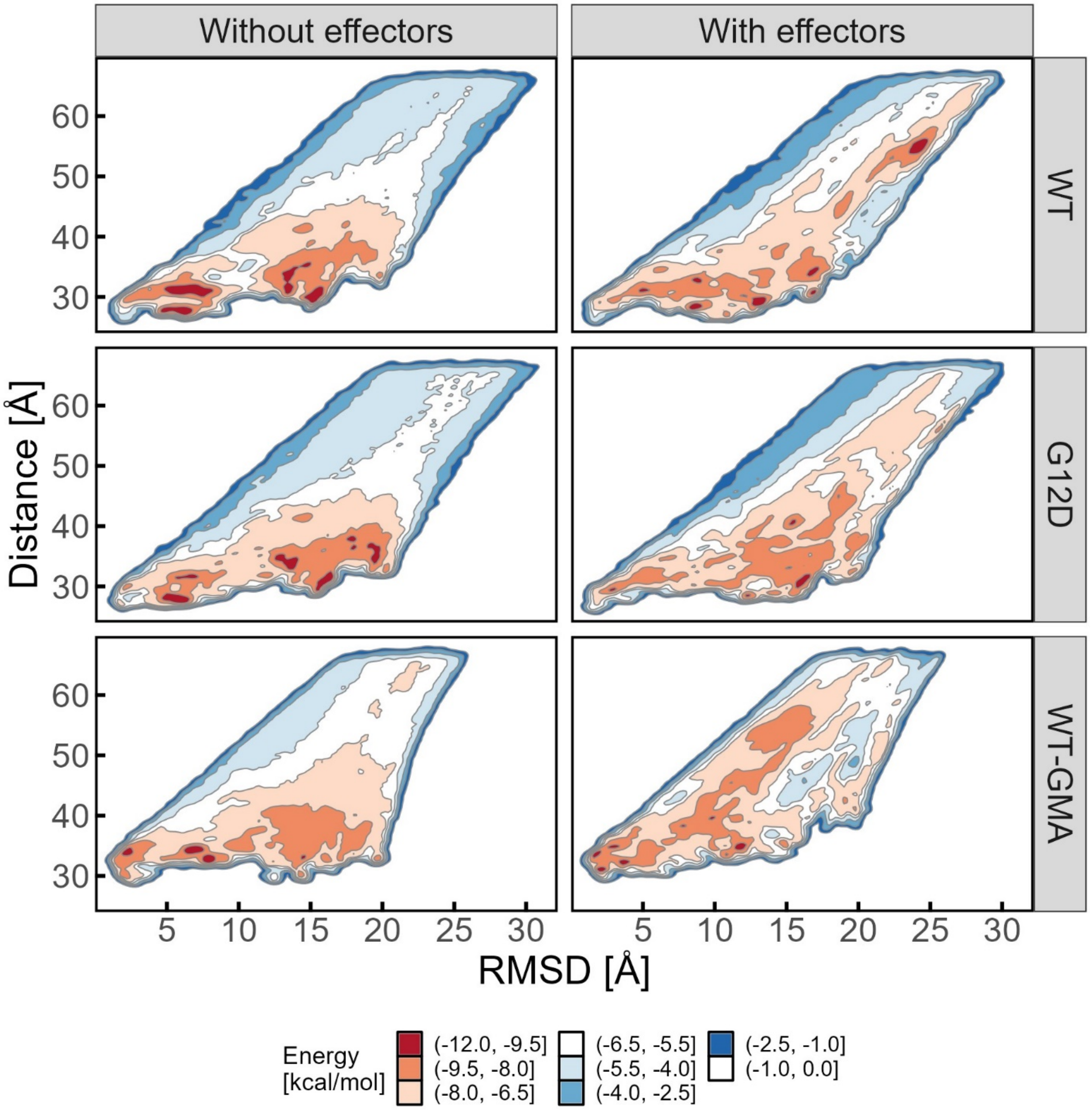


**Figure S8.** Unlabeled 2D free energy landscapes of K-Ras4B system. The simulations have been grouped vertically by the presence/absence of Raf effectors and horizontally by the system (WT, G12D and WT-GMA). The X and Y axes correspond to the RMSD and inter-monomer distance CV values, respectively. The coloring indicates the energy value associated with each coordinate and ranges from dark red (global minimum) through white and blue to white (0). Energy values have been normalized according to the lowest energy observed across all systems (WT-GMA without effectors, global minimum).

**Simulations without c-Raf [RBD-CRD] effectors identify existing interfaces and reveal interplay between interfaces**

The number of minima identified in the deepest part of the free energy landscape (free energy < -9.5 kcal/mol) is six, ten and four for the WT, G12D and WT-GMA simulations without effectors, respectively. Of these, three (50%), eight (80%) and one (25%), for the WT, G12D and WT-GMA simulations, respectively, are structurally far from the initial state with PDB ID 5VQ2(RMSD > 10 Å) of each simulation.

For the WT-GMA simulations without effectors, the three deepest minima of the WT-GMA simulation are located close to the native state (RMSD < 10 Å) with the third deepest minimum (representative structure of minimum "C") only 2.5 Å away from the structure of the native state of the simulation. Additionally, none of the WT-GMA simulation minima resemble either of the α4-α5 interfaces (PDB entries 5VQ2 and 6W4E), with the closest one being minimum "I", which has RMSD values of 11.1 and 12.2 Å, from PDB entries 5VQ2 and 6W4E, respectively.

Figure S9 shows the comparison of all identified minima against crystallographic structures of interest. This analysis further clarifies the similarities we noted earlier as well as differences. For example, in the simulations without effectors no minimum resembles PDB entry 3KKN[14] (RMSD < 10 Å), which we have selected as representative of the α3-α4 dimerization interface. The minima of the WT-GMA simulation do not resemble the reference structures as frequently as those for the 5VQ2-based simulations (WT and G12D). Specifically, only 23% of the WT-GMA minima (7/31) are similar (RMSD < 10 Å) to any reference structures. The respective numbers for the WT and G12D simulations are 29 (9/31) and 43% (17/40). Similarly, only 1.9% (7/372) of all structural comparisons for the WT-GMA simulation indicate acceptable similarity or better (RMSD < 10 Å) as opposed to 3 (11/372) and 5.2% (25/480) for the WT and G12D simulations, respectively.

The minima of the three simulations without effectors share comparable degrees of similarity, with the inter-simulation percentages of dissimilar structures (RMSD ≥ 15 Å) between WT and G12D, WT and WT-GMA, and G12D and WT-GMA equal to 75.6, 77.4 and 81.6%, respectively. Intra-simulation percentages of dissimilar minima (RMSD ≥ 15 Å) are 75.1, 74.9 and 69.9%, with the WT-GMA simulation standing out because the identified minima from that simulation share a higher degree of similarity with each other compared to the other two simulations.

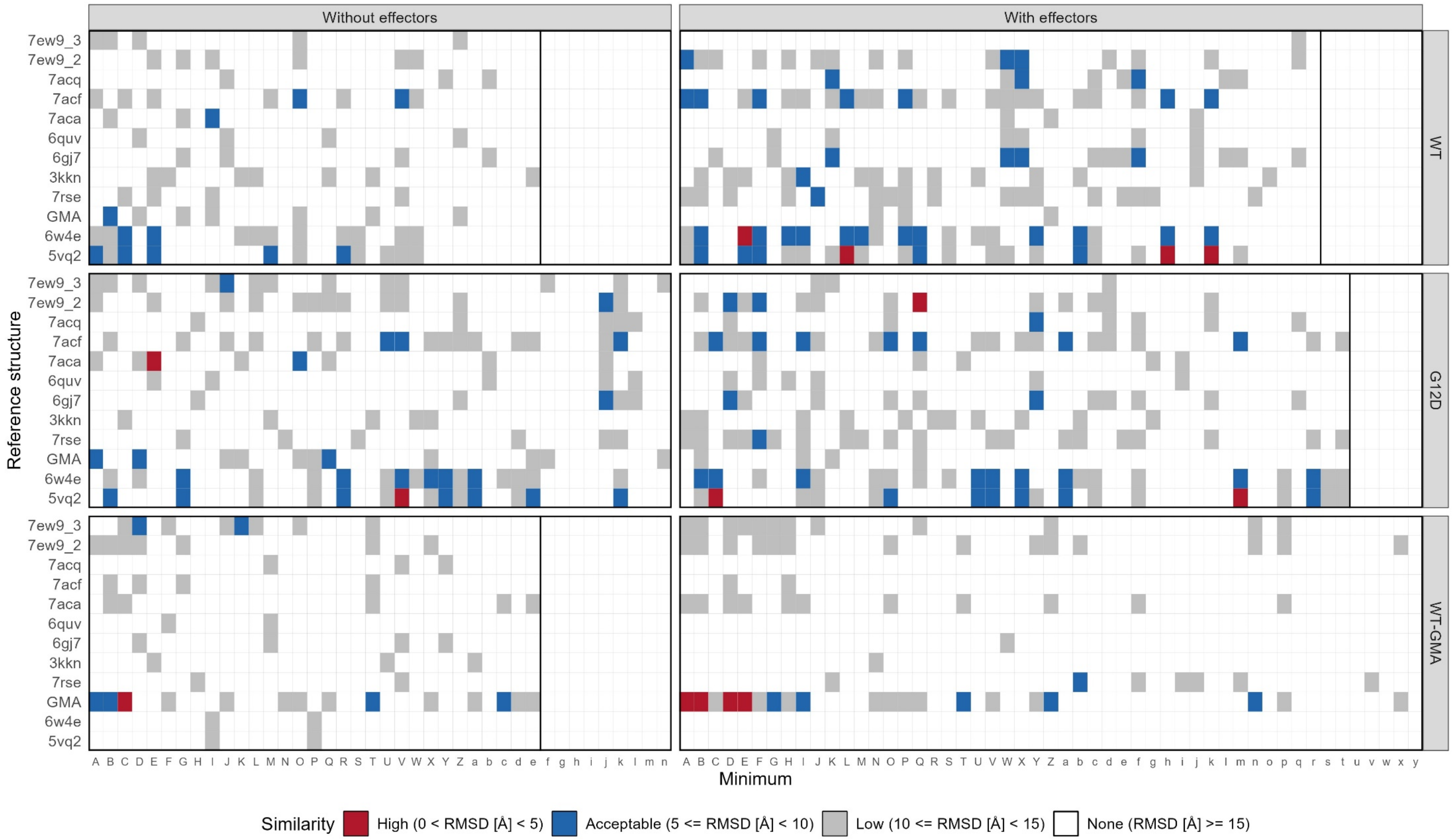


**Figure S9**. Similarity matrix of all representative minima structures against structures of interest. Each cell shows the similarity between a reference structure (Y axis) and a minimum from each of the six simulations (X axis = minimum rank). Cells colored red, blue, grey and white indicate highly, acceptably and lowly similar as well as completely dissimilar structural pairs, respectively. Vertical black lines indicate the number of minima identified per simulation. References for PDB entries 5VQ2, 6W4E, 3KKN, 6GJ7, 6QUV, 7RSE, 7ACA, 7ACF, 7ACQ and 7EW9_2 and 7EW9_3, and structure GMA in the main manuscript.

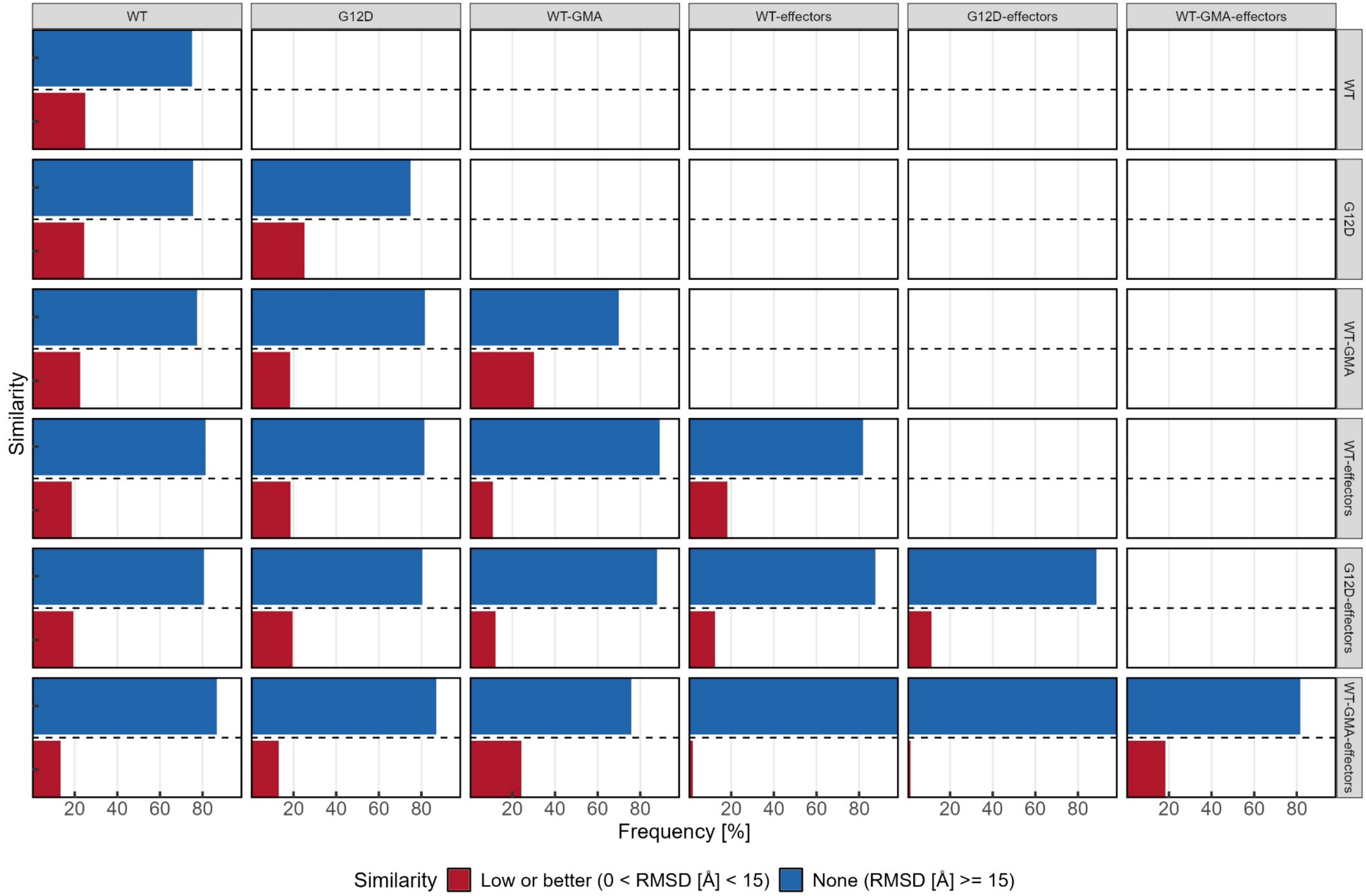


**Figure S10**. Percentage of representative minima structures that share similar structural features across simulations. Each panel shows a comparison between the minima of one simulation (grey box on top) and another (grey box on the right). The percentages of similar structures and dissimilar structures are represented as red- and blue-colored bar plots. The percentages have been calculated after excluding the self-comparisons in the boxes that compare the minima of a simulation with itself (diagonal). The cut-off value used for similarity is 15 Å, with structures whose RMSD is below that value considered similar. The RMSD values were computed over the backbone beads of the K-Ras4B (minus the HVR residues) and Raf proteins.

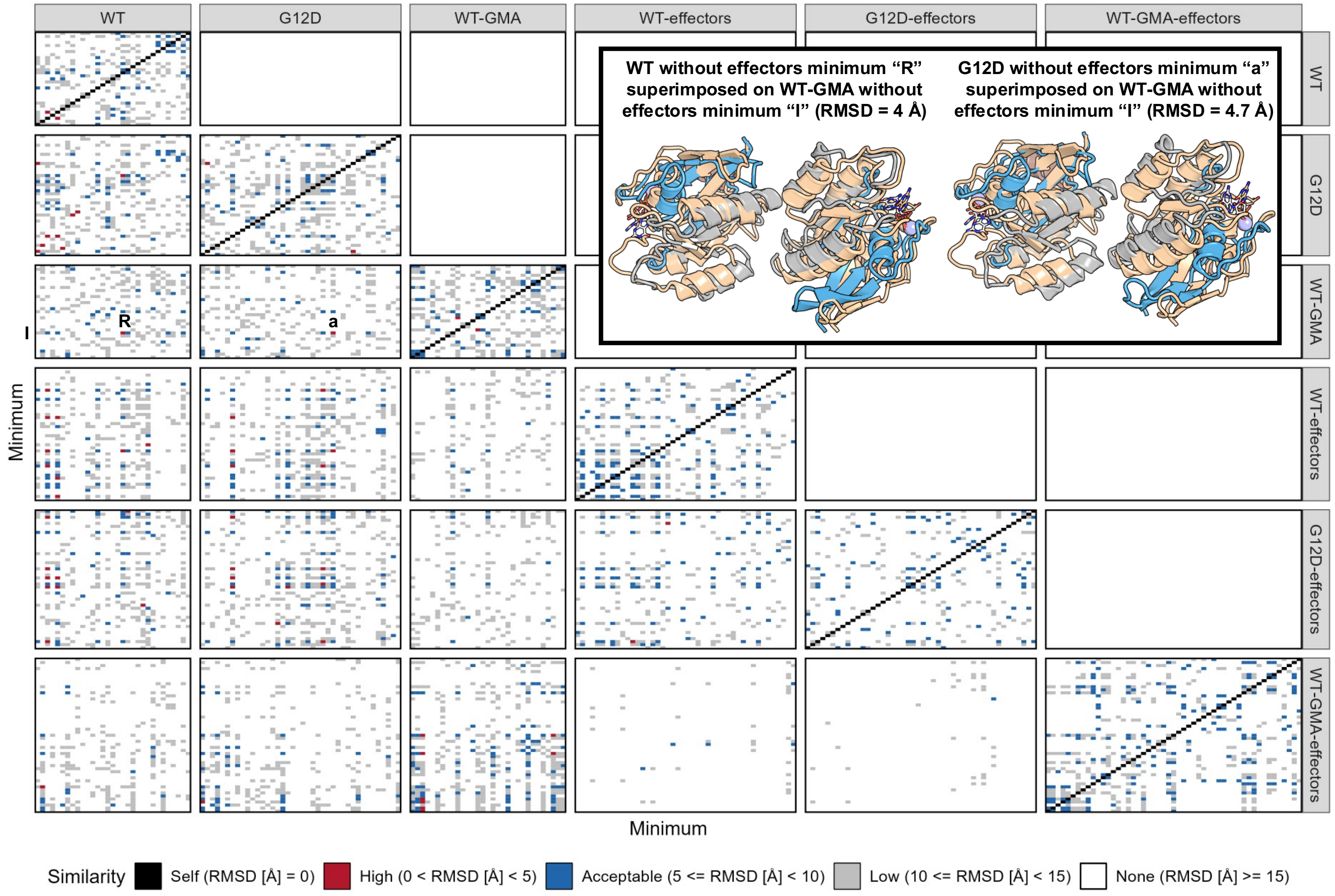


**Figure S11**. All-vs-all similarity matrices for all minima representative structures for all simulations. Same as for figure S8, each box compares the minima from one simulation (grey box on top) against the minima of another (grey box on the right). Each comparison is shown as a colored cell according to the RMSD value of the comparison with identical, highly, acceptably and lowly similar and completely dissimilar structures colored black, red, blue, light grey and white, respectively. The RMSD values were computed over the backbone beads of the K-Ras4B (minus the HVR residues) and Raf proteins. Comparisons of minima “R”, “a” and “I” of the WT, G12D and WT-GMA simulations without effectors, respectively, are indicated on the respective matrices. Inset shows structural comparison of these minima, with coloring the same as for Figure SXX.

**Simulations with c-Raf [RBD-CRD] effectors show significance of c-Raf-mediated interactions**

In terms of similarities with known interfaces, 32 minima closely resemble (RMSD < 10 Å) the interfaces the simulations are being biased toward, with PDB entries 5VQ2/6W4E, and structure GMA featuring prominently in the analysis of the WT/G12D, and WT-GMA simulations, respectively (see Figure S9, right column). Four of the remaining reference structures that resembled minima of the same simulation without effectors now match multiple minima (7ACF, 6GJ7, 7EW9_2 and 3KKN for the WT and G12D simulations). Reference structure 7ACA had a close match in the G12D simulation without effectors (minimum "E", RMSD < 5 Å), but it is not found in any of the minima of the simulations with effectors. In contrast, reference structure with PDB ID 7EW9_2 features three matches in the WT simulation with effectors (minima "A", "W" and "X", RMSD < 10 Å), but none in the same system without effectors, and structure with PDB ID 7RSE, has no matches in any of the simulations without effectors, but one match in each of the WT and G12D simulations with effectors (minima "J" and "F", respectively, RMSD < 10 Å).

The landscape of the WT-GMA simulation shows that the system is more constrained that the other two, with four of the top five and six of the top ten minima measuring less than 5 and 10 Å from the native state (GMA), respectively, in terms of RMSD. The same analysis for the other two simulations reveals only three and one of the top ten minima measure less than 10 Å RMSD from the native state of those simulations (5VQ2). Unlike the simulations without effectors, here the WT and G12D simulations seem to explore a completely different part of the conformational landscape compared to the WT-GMA simulation. This is illustrated by the percentage of WT-GMA minima structures that have no similarity to the minima of the WT or G12D simulations, which is approaching 100% (see Figure S10). Unsurprisingly, the only simulation with similar

(RMSD < 15 Å) minima to the WT-GMA simulation is the equivalent simulation without effectors, with all the minima that converge to structure GMA (see Figure S9) amounting to 19% of all pairs (see Figure S11).

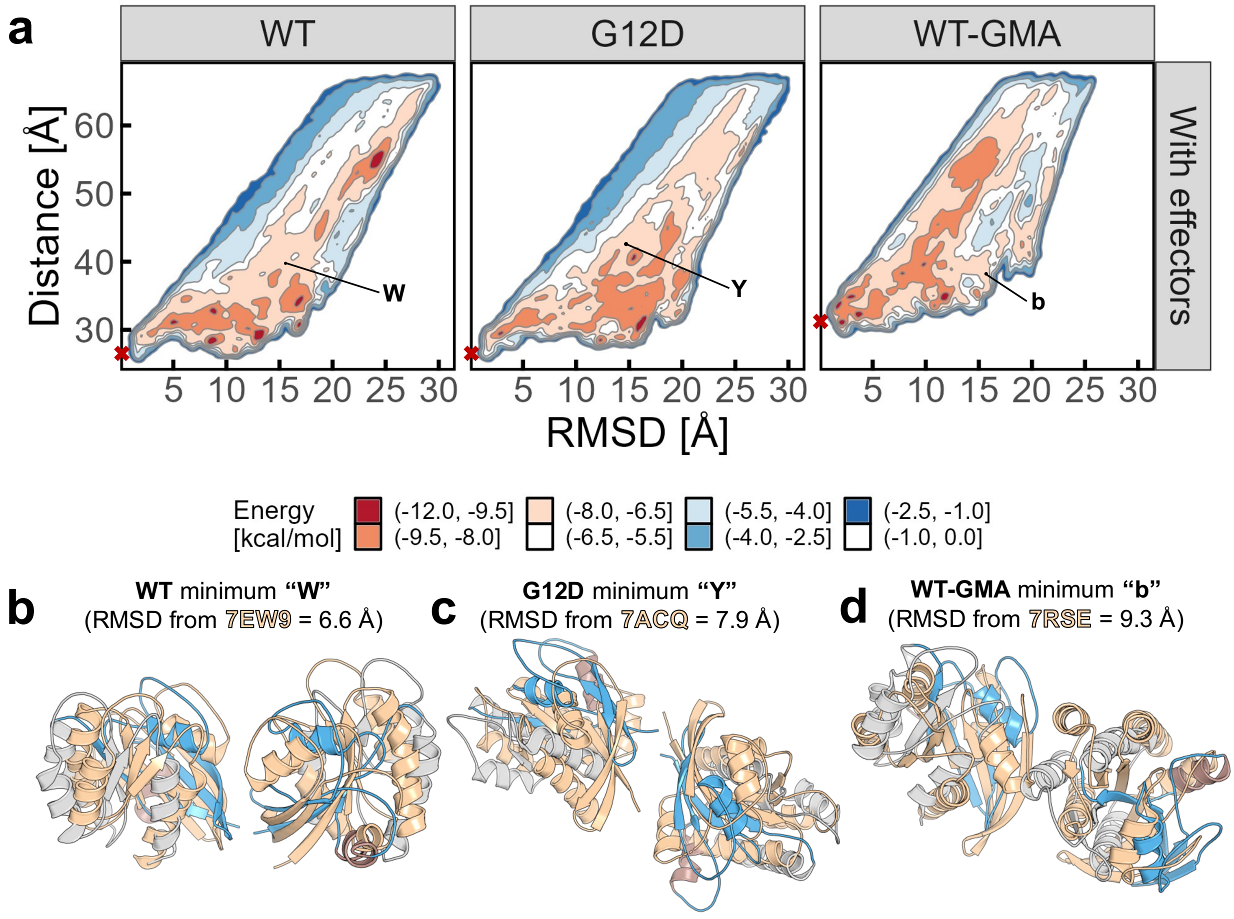


**Figure S12.** 2D free energy landscapes of K-Ras4B with c-Raf [RBD-CRD]. See Figure 3 for details. First conformer of the 7RSE ensemble chosen for visualization and RMSD calculation.

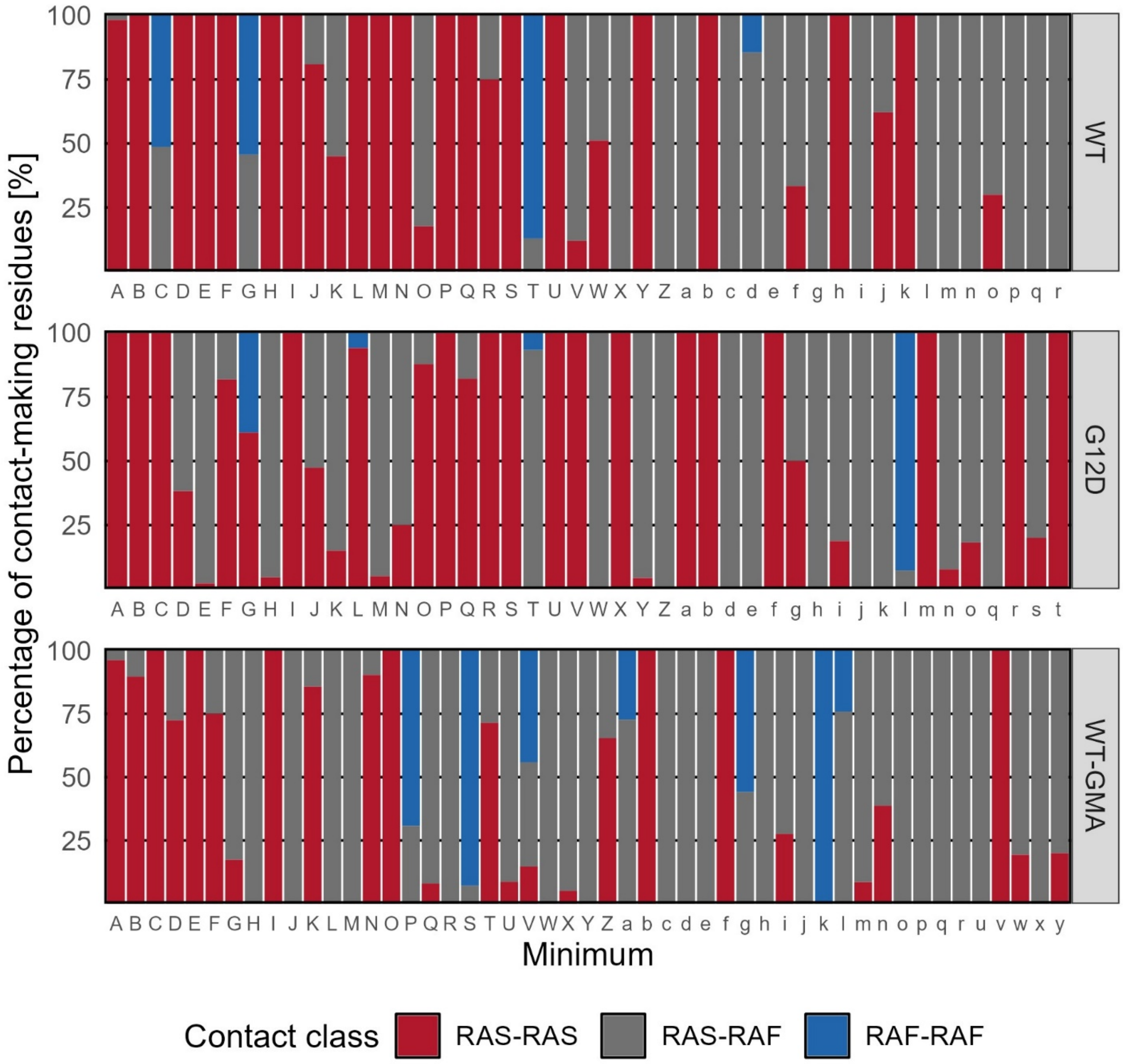


**Figure S13**. Percentages of the contact-class all the contact-making residues for each minimum structure belong to. Each vertical bar corresponds to the contact class percentages of the contact-making residues of one minimum structure (minimum rank on the X axis) for one simulation (indicated in the grey box on the right of each panel). The percentages are shown as stacked bars with their color indicating the contact class they belong to, with residues making RAS-RAS, RAS-RAF and RAF-RAF contacts colored, red, grey and blue, respectively. Vertical black lines for the WT and G12D simulations indicate the number of minima considered for these simulations. Minima “c”, “p” and “s”, “t” from the G12D and WT-GMA simulations respectively only form contacts through their HVR residues which are excluded from this analysis. Specifically, 47, 47 and 35% of the contacts are between K-Ras4B residues, for the WT, G12D and WT-GMA simulations, respectively. RAS-RAF contacts are 45, 48 and 54% of the total for the WT, G12D and WT-GMA simulations, respectively, and RAF-RAF contacting residues form 8, 5 and 11% of the contacts for the WT, G12D and WT-GMA with effector simulations, respectively.

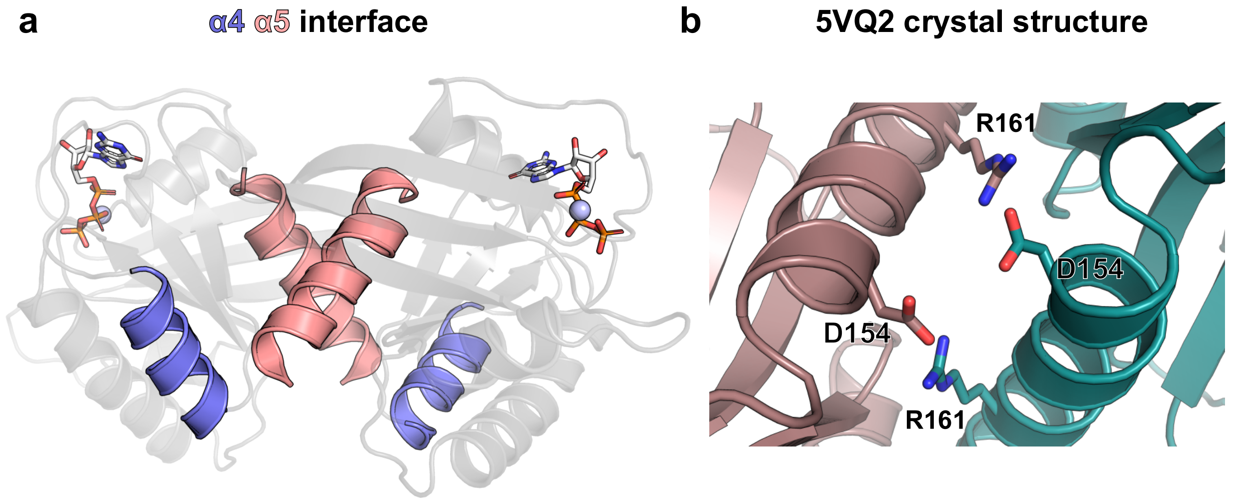


**Figure S14**. A depiction of the characterized α4-α5 interface, where R161 and D154 interact with two salt bridges in the PDB ID: 5VQ2.

**Figure 6 structural explanation**

In Figure 6b, the light blue and grey areas in the RAS-RAS contact maps (see Figure 6b, first row) indicate the regions towards which each simulation is being biased, because they represent the contact network of reference structures 5VQ2 (for the WT and G12D simulations) and GMA (for the WT-GMA simulation), which explains the increased density of contacts in those regions for those panels. Of the two α4-α5-biased systems (WT and G12D), the WT system exhibits this tendency more strongly, whereas the contacts of the GMA-biased system (WT-GMA) appear shifted with respect to the reference interface, but still clearly following the trend.

Figure 6c shows the interaction between K-Ras4B helices α4-α5 for the D154-contributing protomer and K-Ras4B helices α3-α4 for the other protomer. Figure 6d shows the interaction between the D154-contributing K-Ras4B protein of one protomer and the c-Raf protein of the other protomer with K-Ras4B helices α4-α5 interacting with the RBD domain. Figure 6e shows one minimum structure from the WT-GMA simulation that features interactions involving K-Ras4B residue K88, used for the validation of the helical stacking mechanism of the GMA model.[15] The α3 helix – and by extension residue K88 as well – is part of the interface which features the α3-α4 helices for one protomer and α3 helix and GTP region for the other.

**Comparison of the unbiased and biased simulations**

We also compared the similarity of the minima identified in the CG-PT-MetaD-WTE simulations with the representative structures extracted from the unbiased simulations by calculating the percentage of minima which shared a predetermined level of similarity (in terms of RMSD) with any of the unbiased simulation representative structures. The results of this analysis are shown in Figure S15. Unsurprisingly, the minima of the simulations with effectors tend to be more similar to the representative structures of the unbiased simulations, since those included the Raf effectors as well. Nowhere is this more apparent than the WT simulation where only two (6.5%) of the minima in the simulation without effectors resemble (RMSD < 10 Å) any of the unbiased simulation representative structures but 20 (45.6%) minima in the simulation with effectors do (similarity class high or acceptable, RMSD < 10 Å). Figure S16 highlights the high-similarity (RMSD < 5 Å) convergence points of the biased and unbiased simulations. Interestingly, none of these convergence points feature direct inter-protomer K-Ras4B interaction. Instead, in two of the three cases (Figure S16a,c) the complex comes together in a cross-like way, whereas in Figure S16b the four subunits are arranged in a linear fashion on top of the membrane with one K-Ras4B subunit interacting solely with the c-Raf subunit of its partner. Figure S17 shows all comparisons between the unbiased simulation representative structures and minima structures.

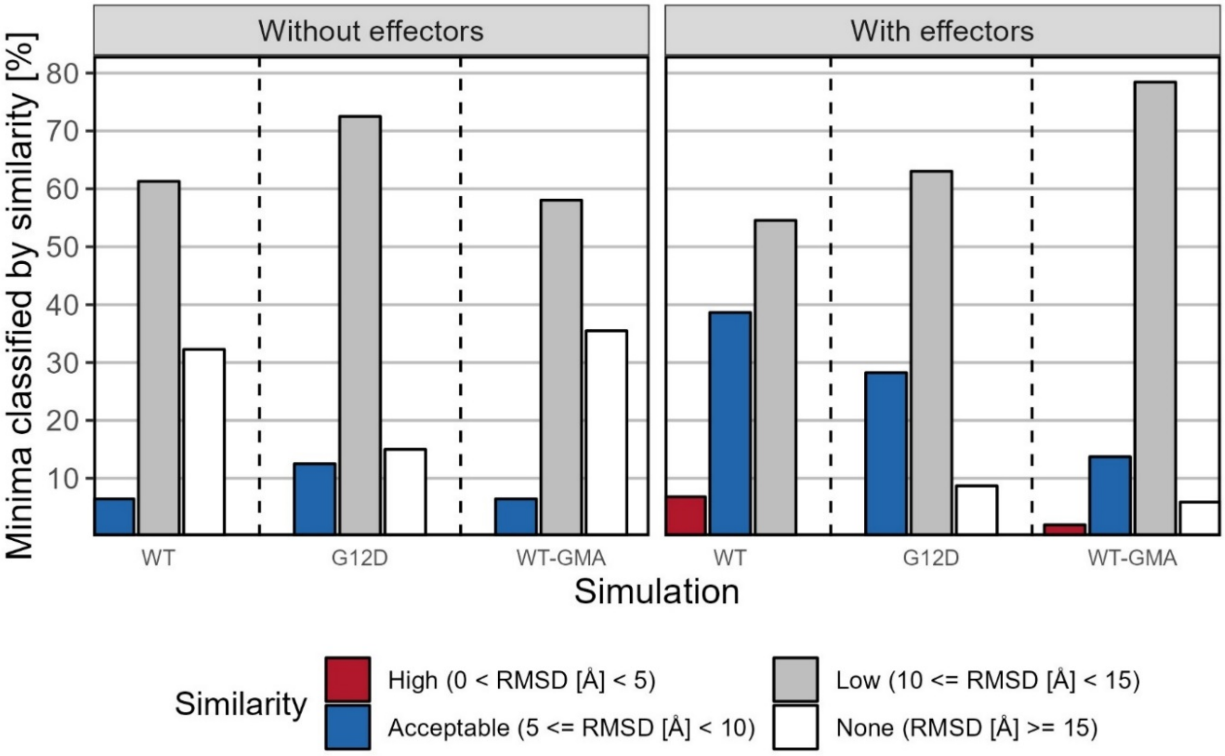


**Figure S15.** Percentage of minima belonging to each similarity class for all simulations after comparison with the unbiased simulation representative structures. The similarity class percentages are shown as bars colored according to their similarity class, with high, acceptable, low and no similarity, colored red, blue, light grey and white, respectively. Vertical dotted lines separate the simulation groups. Whenever a minimum structure had more than one matches among the unbiased simulation representative structures, only the higher-similarity one was kept for this analysis.

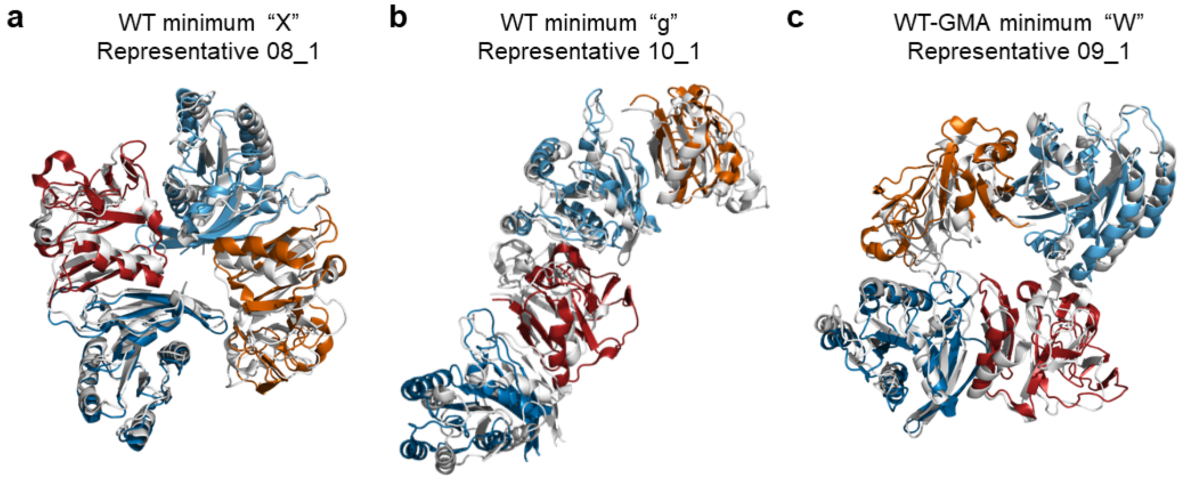


**Figure S16.** Selection of high-similarity minima and unbiased representative structure superimpositions. Panels a, b and c show top-side views of WT minimum "X", WT minimum "g" and WT-GMA minimum "W" superimposed on representative structures from the unbiased simulations replicas 8, 9 and 10, respectively. All unbiased representative structures are colored white and the minima structures are colored the same way as in Figures 2 and 5. K-Ras4B HVR residues, all ions and GTP cofactor not shown for clarity.

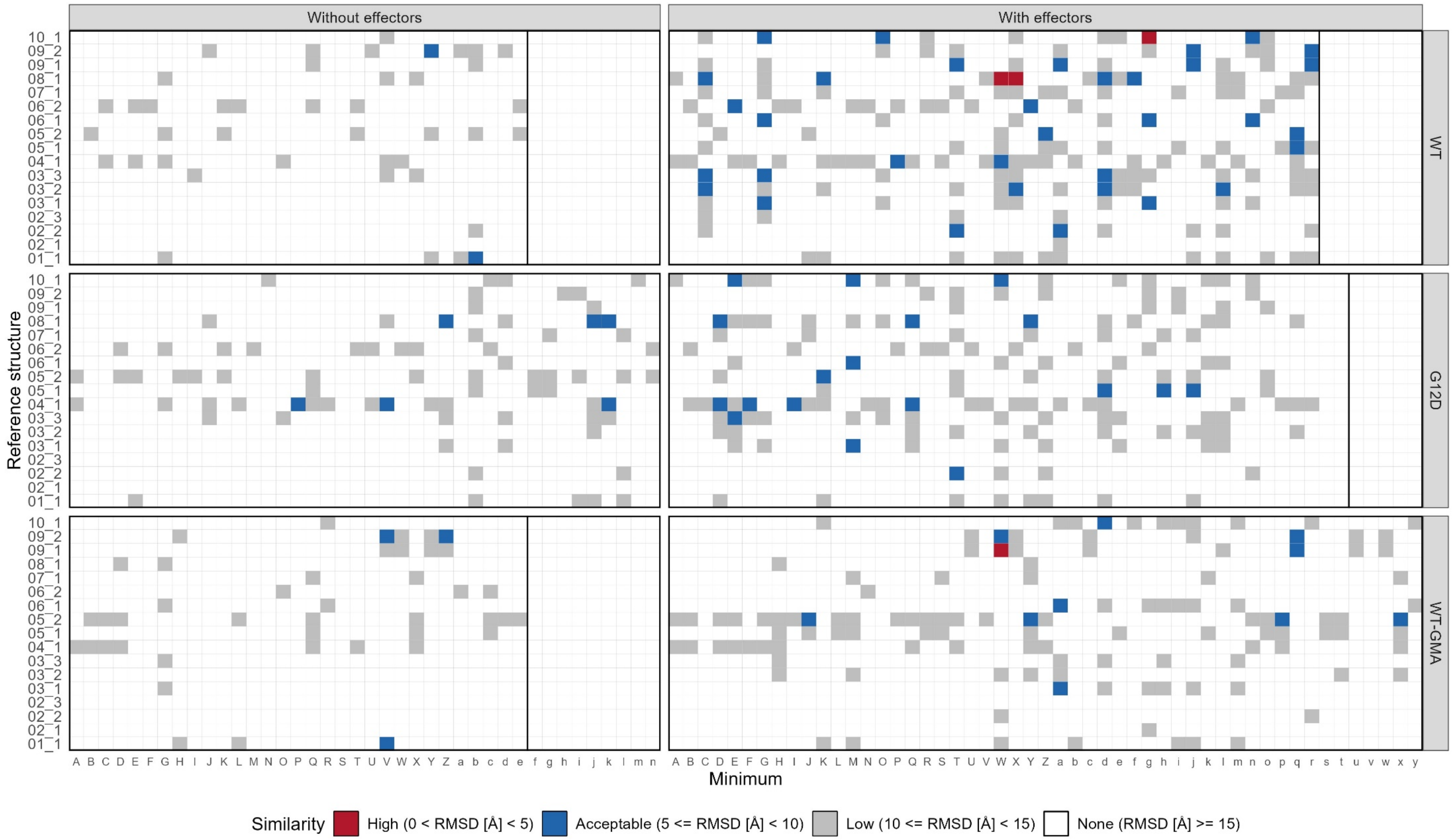


**Figure S17.** Similarity matrix of all representative minima structures against representative structures from the unbiased simulations. Each cell shows the similarity between an unbiased simulation representative structure (Y axis) and a minimum from each of the six simulations (X axis = minimum rank). Vertical black lines indicate the number of minima identified per simulation. Coloring is the same as for Figure S8.

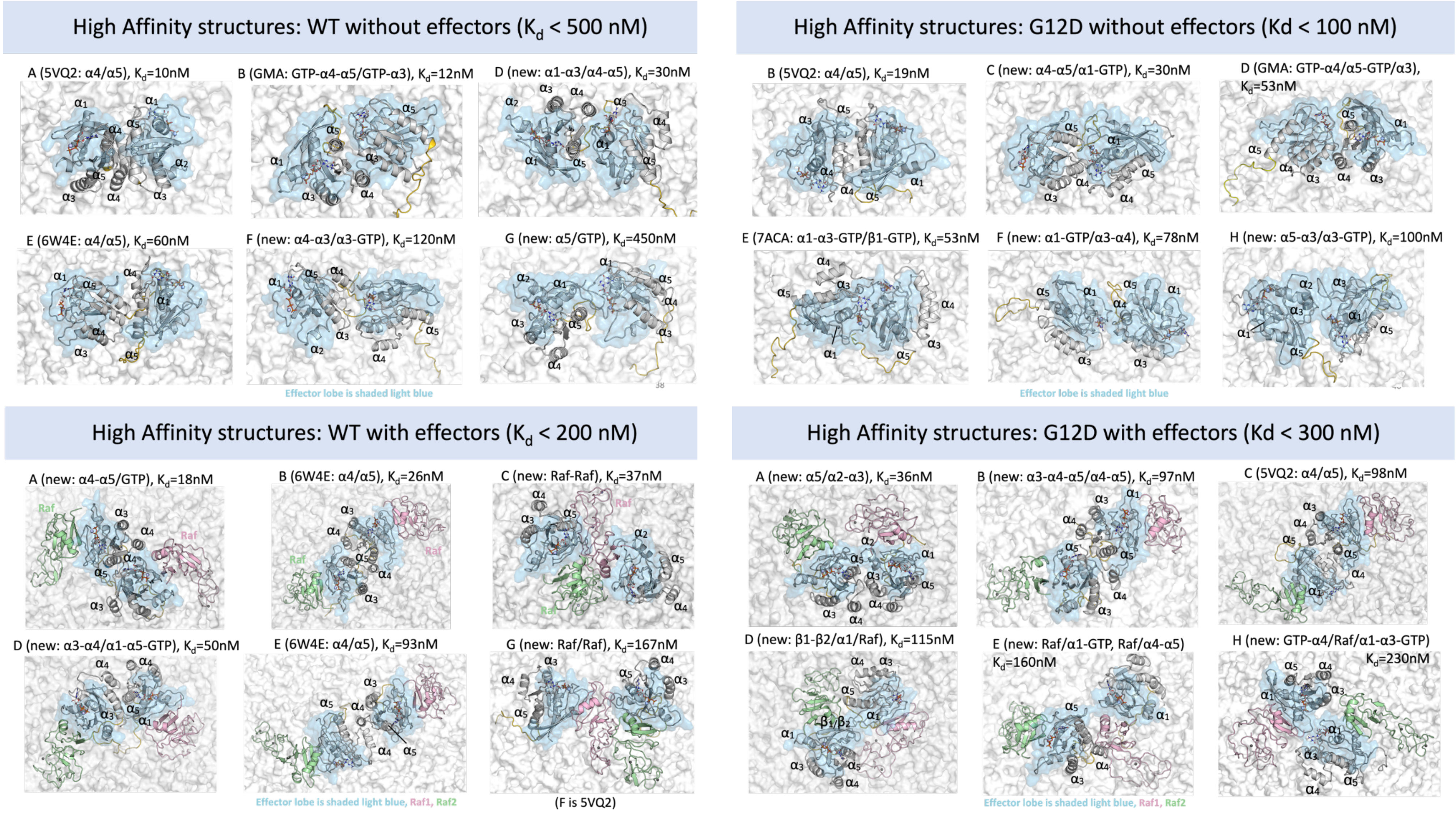


**Figure S18**. High affinity dimer structures for all simulations. Left column shows structures from the WT simulations with (bottom) and without (top) effectors. Right column shows structures from the G12D simulation with (bottom) and without (top) effectors. K-Ras4B helices of interest are labeled. K-Ras4B are shown as grey cartoons, Raf are shown as pink and green cartoons, GTP are shown sticks and $Mg^{2+}$ as light blue spheres. The affinity values reported in the figure correspond to the highest-affinity structure of each minimum. The structure corresponds to the representative structure of each minimum. PDB codes indicate the reference structure some of the minima resemble.

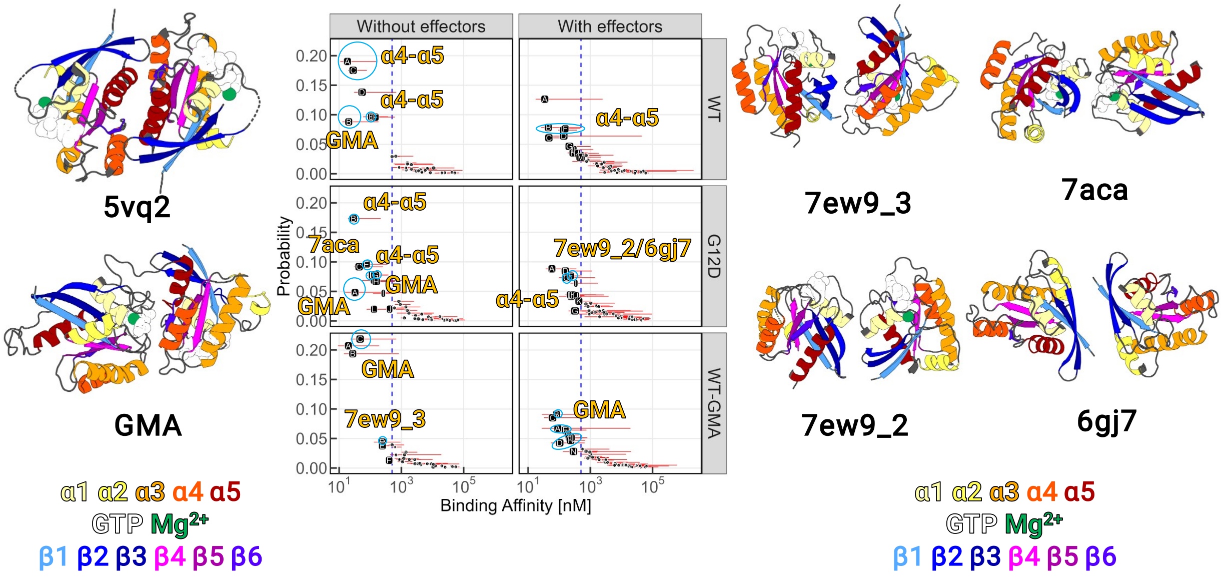


**Figure S19**. Selection of minima that resemble reference structures and whose affinity is ≤ 500 nM. The dashed blue line corresponds to a binding affinity of 500 nM, and is used to identify high-affinity minima (left of the line). Probability (Y axis) indicates the population of each cluster as a ratio of the total number of trajectory frames that were grouped in clusters. Cyan circles/ellipsoids indicate the minima that resemble the reference structure indicated in the gold label close to the indicated minima. The X axis for all panels is in a $\log_{10}$ scale. Cutoff for similarity was set to 6.5 Å.

**Protein-lipid interactions**

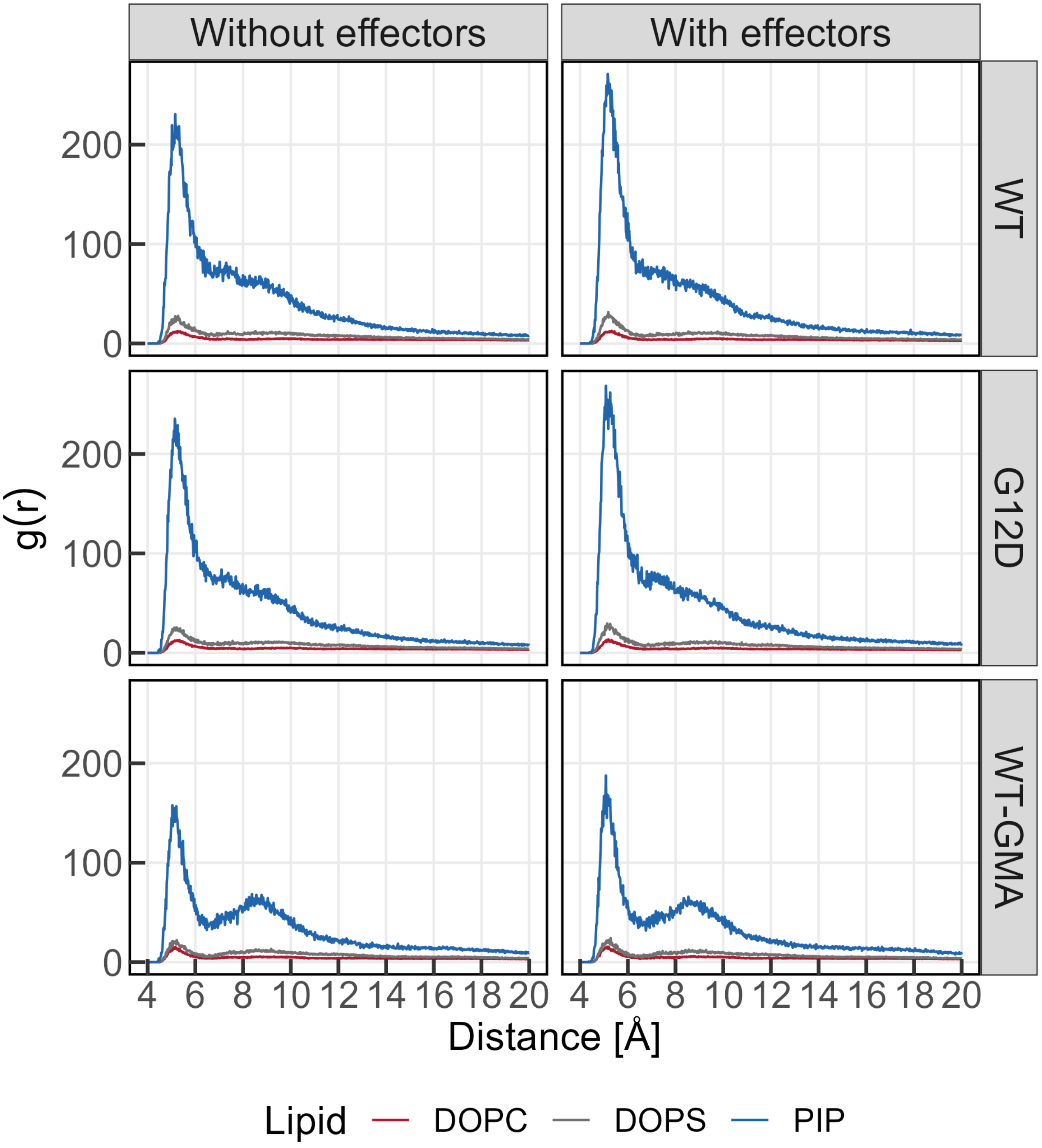


**Figure S20.** Radial distribution function distributions for the PO4 beads of DOPC, DOPS and $PIP_2$ lipids as calculated from the backbone bead of K-Ras4B T183. The distributions for DOPC, DOPS and $PIP_2$ are shown as red, grey and blue lines, respectively.

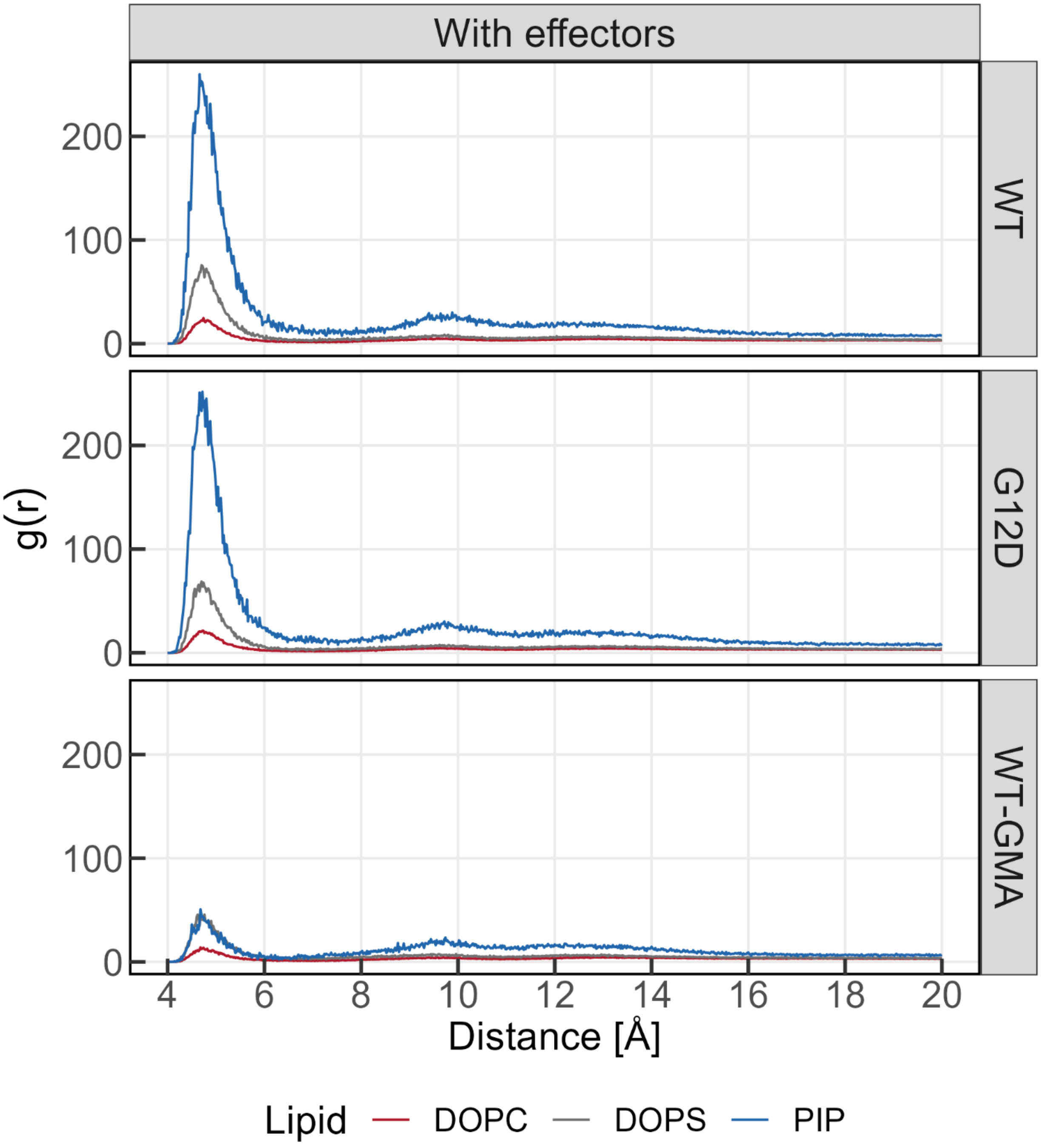


**Figure S21.** Radial distribution function distributions for the PO4 beads of DOPC, DOPS and $PIP_2$ lipids as calculated from the SC2 bead of c-Raf K148. Same coloring as for Figure S20.

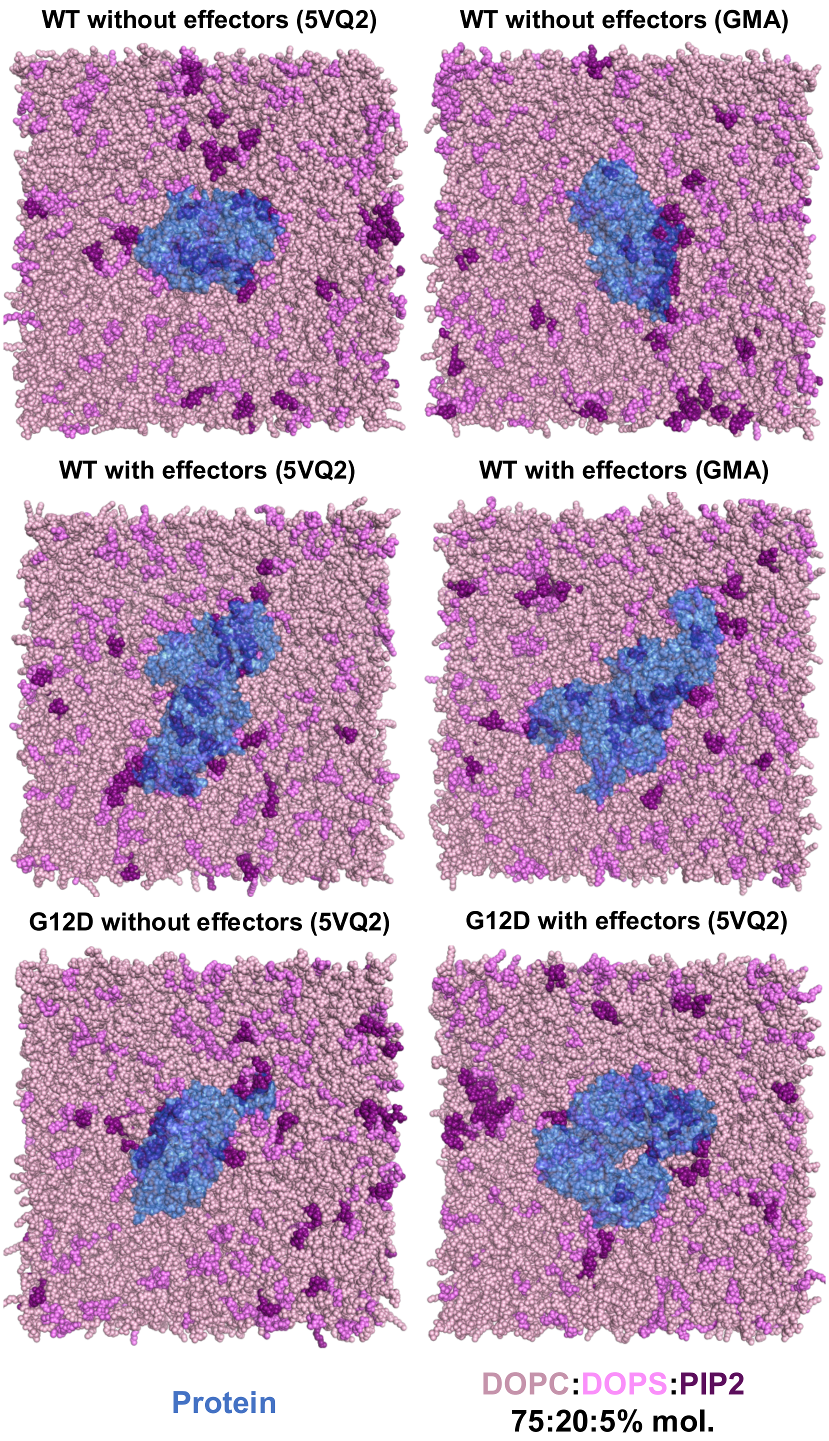


**Figure S22**. Snapshots of the deepest minimum structure of each simulation shown as coordinated by the different lipids on the model lipid bilayer. Brackets indicate the starting/biasing structure for each system.

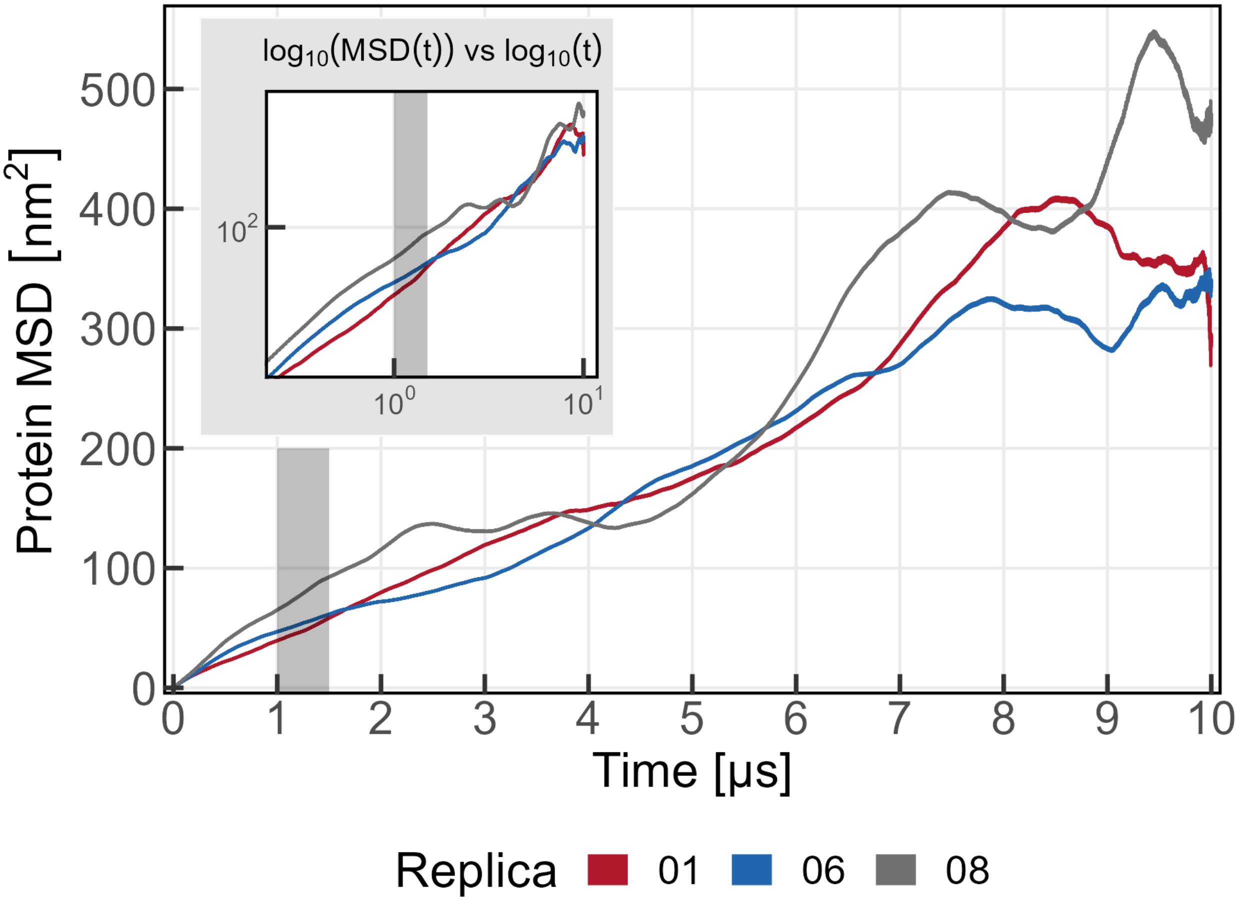


**Figure S23.** Mean squared displacement of the K-Ras4B-c-Raf monomers vs time for all replicas where protein monomers were observed (see Figure S2). The inset shows the $\log_{10}$ transformation of the plot to verify the linearity of the regions selected for diffusion coefficient calculation (shaded area between 1 and 1.5 μs). The MSD values for the three replicas are shown as red, blue and grey colored lines for replicas 1, 6 and 8, respectively.

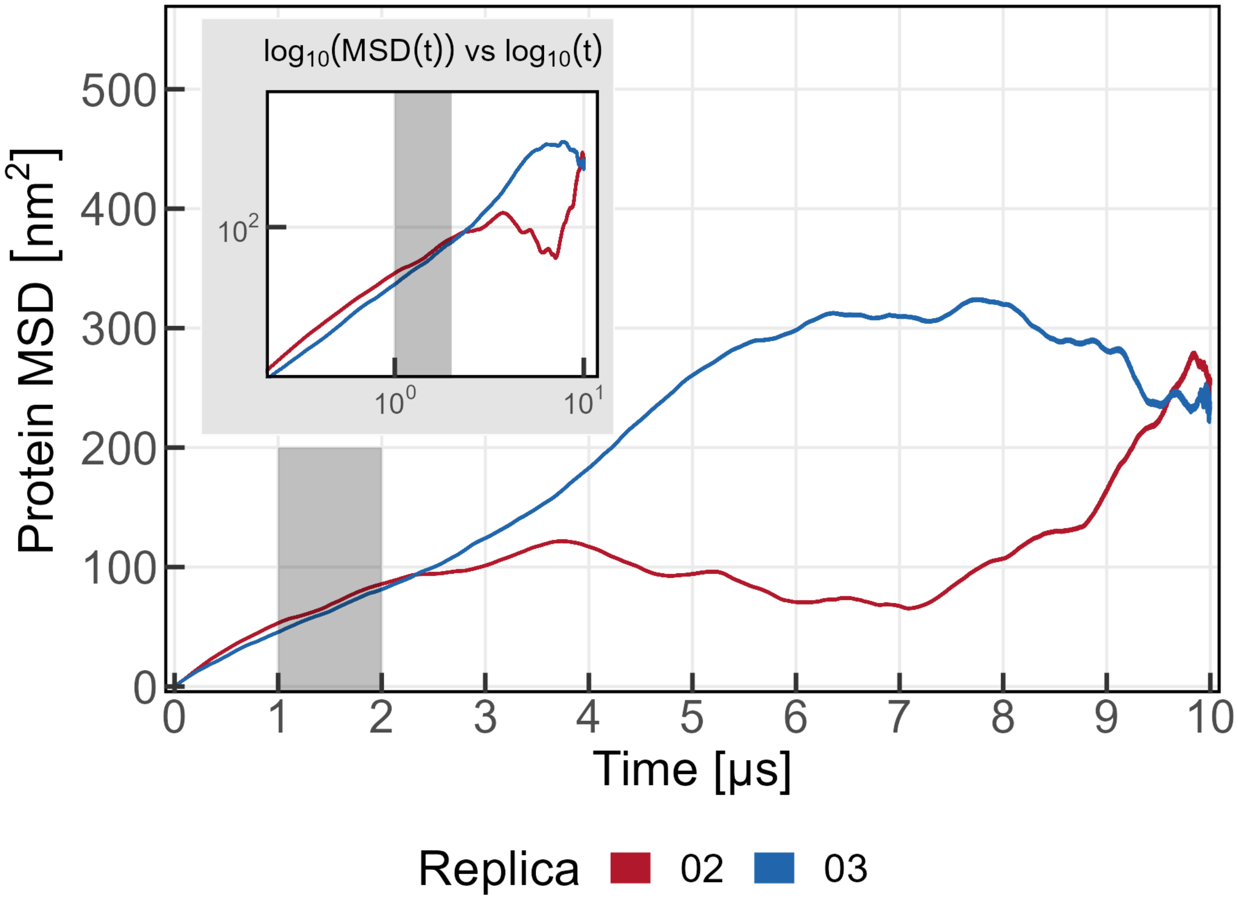


**Figure S24.** Mean squared displacement of the K-Ras4B-c-Raf unstable dimers vs time for all replicas where dimers undergoing multiple association-dissociation events were observed (see Figure S2). The inset shows the $\log_{10}$ transformation of the plot to verify the linearity of the regions selected for diffusion coefficient calculation (shaded area between 1 and 2 µs). The MSD values for the two replicas are shown as red and blue colored lines for replicas 2 and 3, respectively.

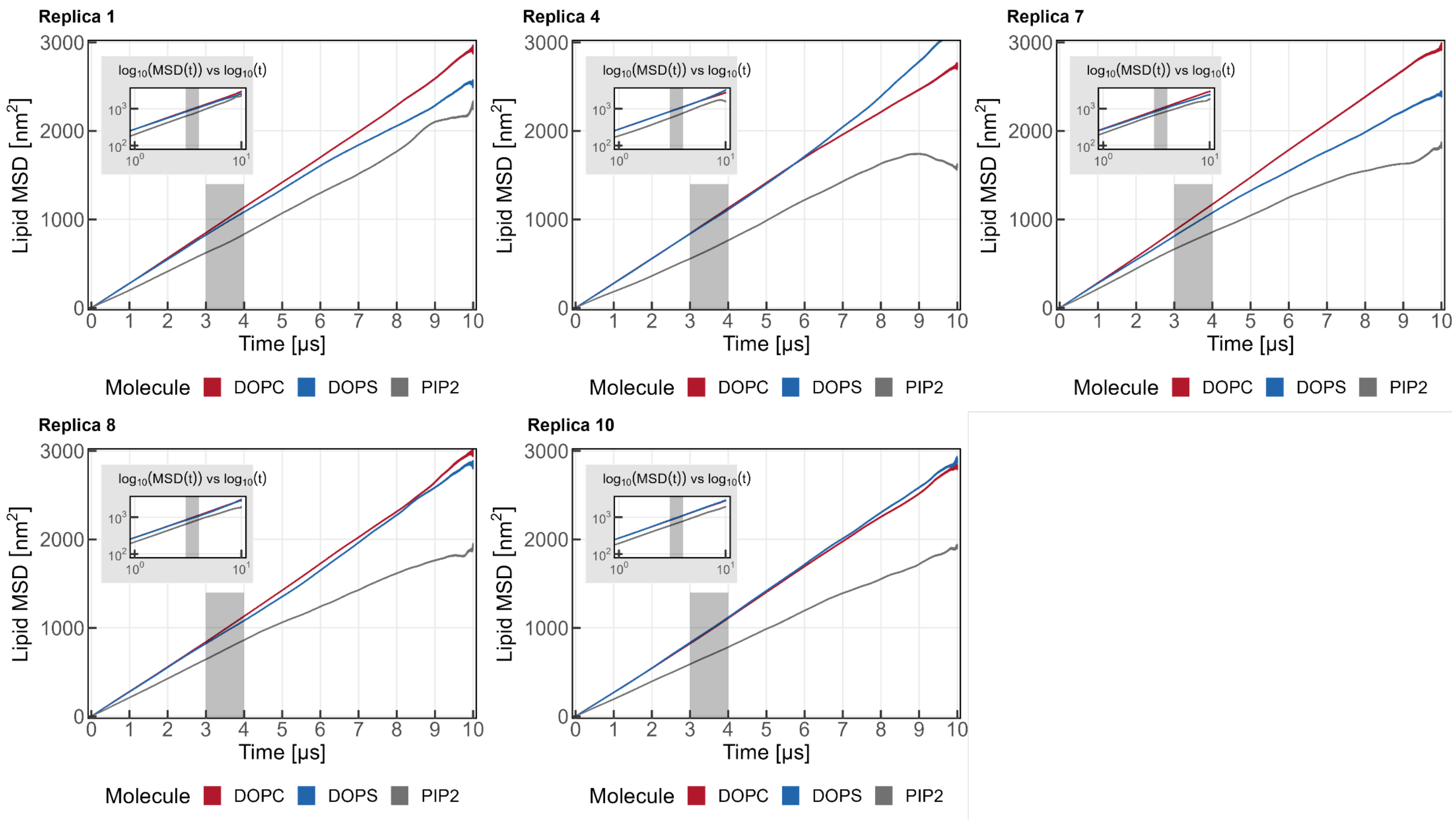


**Figure S25.** Mean squared displacement of the three lipid species vs time for all replicas where stable dimers were observed (see Figure S2). The inset shows the $\log_{10}$ transformation of the plot to verify the linearity of the regions selected for diffusion coefficient calculation (shaded area between 3 and 4 µs). The MSD values for different replicas are shown in separate panels with the panel label indicating the replica it corresponds to. MSD values for DOPC, DOPS and $PIP_2$ lipids are shown as red, blue and grey colored lines, respectively.

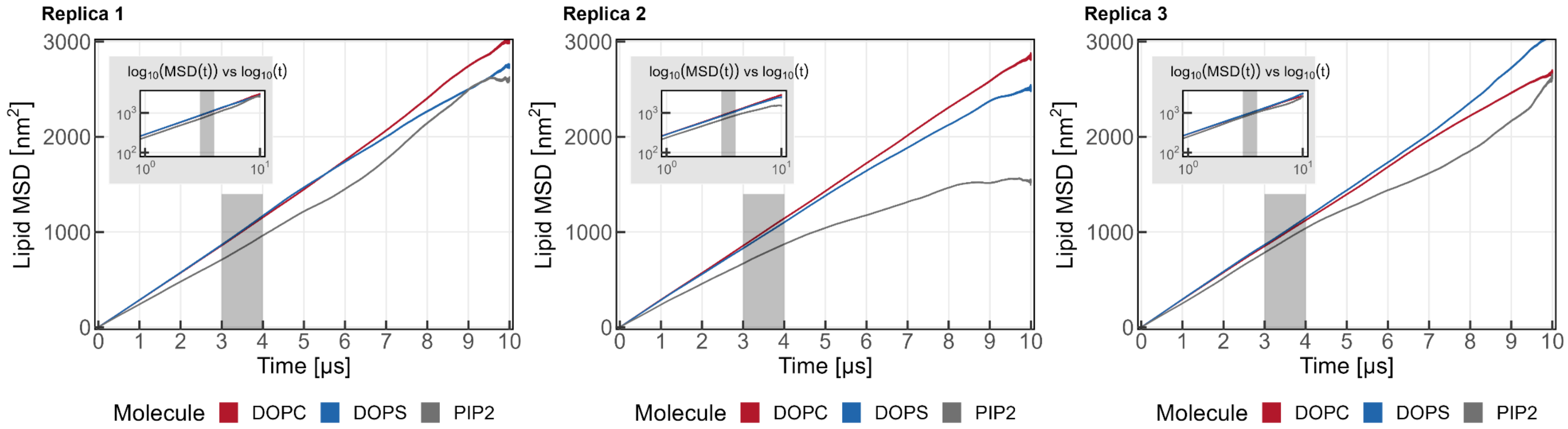


**Figure S26.** Mean squared displacement of the three lipid species vs time for three protein-free replicas. The inset shows the $\log_{10}$ transformation of the plot to verify the linearity of the regions selected for diffusion coefficient calculation (shaded area between 3 and 4 µs). The MSD values for different replicas are shown in separate panels with the panel label indicating the replica it corresponds to. MSD values for DOPC, DOPS and $PIP_2$ lipids are shown as red, blue and grey colored lines, respectively.

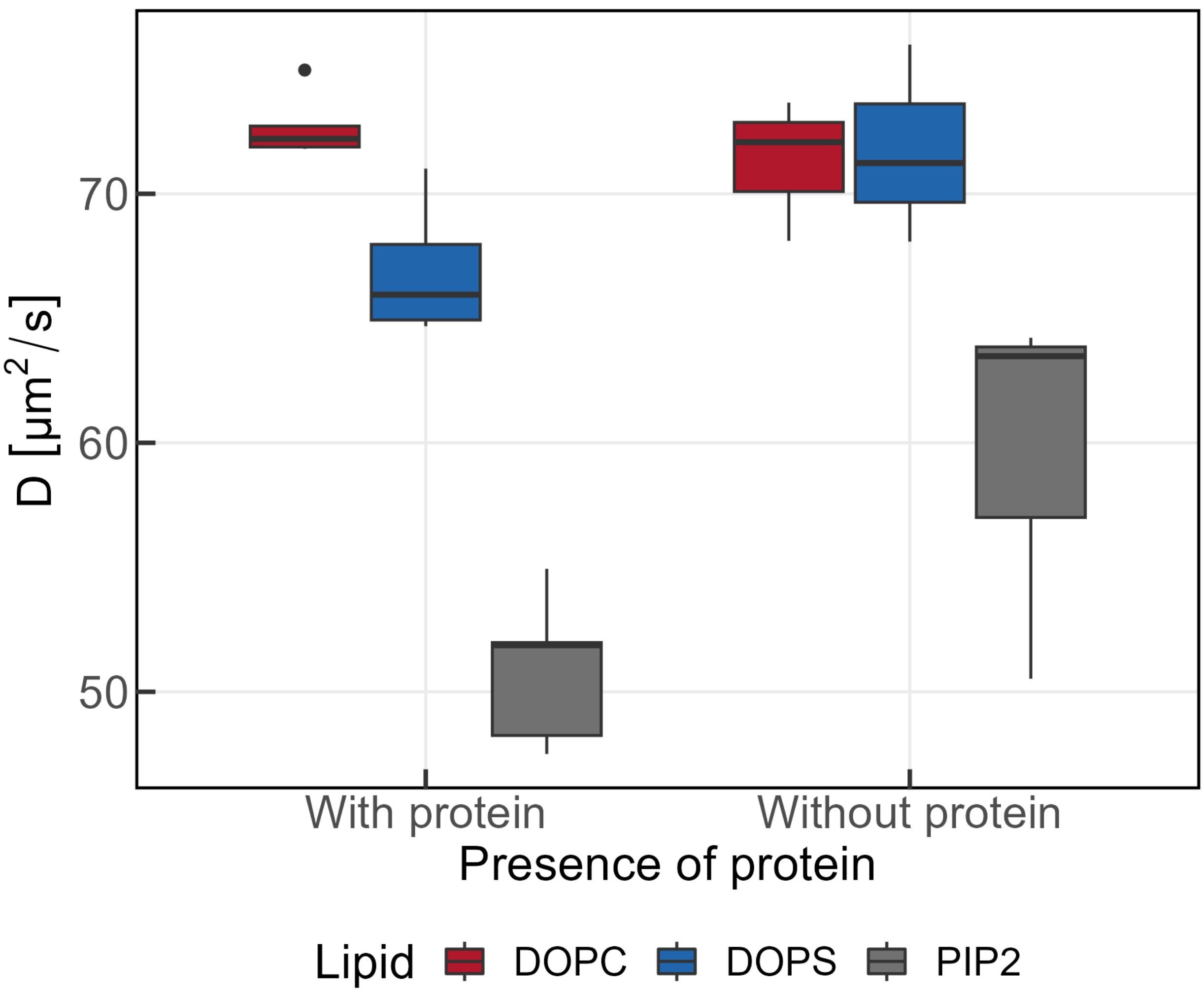


**Figure S27.** Boxplots of the diffusion coefficients calculated from the slope of the shaded regions of the plots shown in plots S25-S26.

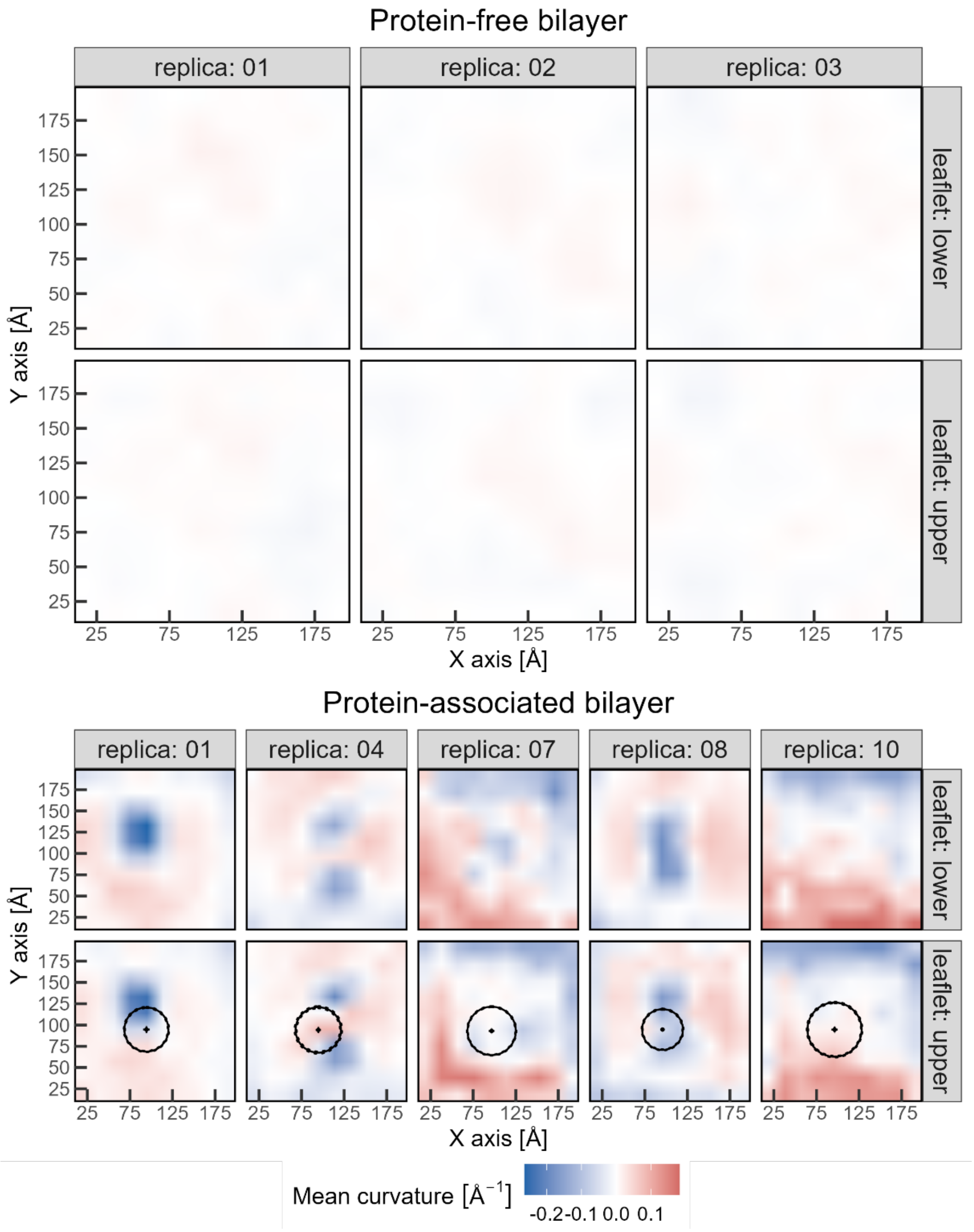


**Figure S28.** Mean curvature for the upper and lower bilayer leaflets in the absence (top panels) and presence (bottom panels) of the K-Ras4B-c-Raf protein complex. Each panel shows the mean curvature on a color scale of deep blue (negative values – concave) to light red (positive values – convex) through zero curvature (0 – flat). The circles in the bottom panels indicate the radius of gyration of the protein complex with the dashed circles corresponding to the average ± 1 standard deviation. The crosses in the center of the circles correspond to the position of the center of mass of the protein complex, with the length of each cross axis indicating the deviation along each dimension and the point where the two cross axes meet corresponding to the average coordinates. The off-white color of the protein-free bilayer panels indicates near-zero curvature.
